# Revealing the Biophysics of Lamina-Associated Domain Formation by Integrating Theoretical Modeling and High-Resolution Imaging

**DOI:** 10.1101/2024.06.24.600310

**Authors:** Monika Dhankhar, Zixian Guo, Aayush Kant, Ramin Basir, Rohit Joshi, Su Chin Heo, Robert L. Mauck, Melike Lakadamyali, Vivek B. Shenoy

## Abstract

The interactions between chromatin and the nuclear lamina orchestrate cell type-specific gene activity by forming lamina-associated domains (LADs) which preserve cellular characteristics through gene repression. However, unlike the interactions between chromatin segments, the strength of chromatin-lamina interactions and their dependence on cellular environment are not well understood. Here, we develop a theory to predict the size and shape of peripheral heterochromatin domains by considering the energetics of chromatin-chromatin interactions, the affinity between chromatin and the nuclear lamina and the kinetics of methylation and acetylation9in human mesenchymal stem cells (hMSCs). Through the analysis of super-resolution images of peripheral heterochromatin domains using this theoretical framework, we determine the nuclear lamina-wide distribution of chromatin-lamina affinities. We find that the extracted affinity is highly spatially heterogeneous and shows a bimodal distribution, indicating regions along the lamina with strong chromatin binding and those exhibiting vanishing chromatin affinity interspersed with some regions exhibiting a relatively diminished chromatin interactions, in line with the presence of structures such as nuclear pores. Exploring the role of environmental cues on peripheral chromatin, we find that LAD thickness increases when hMSCs are cultured on a softer substrate, in correlation with contractility-dependent translocation of histone deacetylase 3 (HDAC3) from the cytosol to the nucleus. In soft microenvironments, chromatin becomes sequestered at the nuclear lamina, likely due to the interactions of HDAC3 with the chromatin anchoring protein LAP2*β* ,increasing chromatin-lamina affinity, as well as elevated levels of the intranuclear histone methylation. Our findings are further corroborated by pharmacological interventions that inhibit contractility, as well as by manipulating methylation levels using epigenetic drugs. Notably, in the context of tendinosis, a chronic condition characterized by collagen degeneration, we observed a similar increase in the thickness of peripheral chromatin akin to that of cells cultured on soft substrates consistent with theoretical predictions. Our findings underscore the pivotal role of the microenvironment in shaping genome organization and highlight its relevance in pathological conditions.

## Introduction

Tissue development, maintenance, and healing necessitate cells to continuously assess their three-dimensional microenvironment and regulate their activity based on local biochemical and mechanical signals[1]. As the hub of cellular genetic information, the nucleus plays a vital role in dictating the destiny and function of cells by changing the packing of chromatin in response to these stimuli [2]. Chromatin is spatially organized into distinct domains: euchromatin, which is loosely packed and transcriptionally active, and heterochromatin, which is densely packed and transcriptionally silent [3]. Crucially, at the nuclear periphery, interactions between chromatin and the nuclear lamina lead to the formation of lamina-associated domains (LADs), which exhibit heterochromatic characteristics, play a pivotal role in functionally organizing the genome and silencing genes that belong to the wrong cell lineage during differentiation [4-6]. LADs appear as genomic features influenced by and influencing epigenomic states and the overall architecture of the genome. Despite previous research emphasizing the significance of LADs, the quantitative strengths of the interactions between chromatin and the nuclear lamina and how they are influenced by cellular microenvironments remain unknown.

To decipher the spatially heterogeneous biophysical mechanisms governing chromatin organization, with a specific focus on the strength of chromatin-lamina tethering and its modulation in response to the cell’s microenvironment, we introduce a multimodal framework. This framework integrates super-resolution microscopy with a mesoscale mathematical model that accounts for interactions between chromatin segments and chromatin-lamina, as well as the kinetics of histone methylation and acetylation. We show that chromatin-lamina interactions and histone methylation and acetylation reaction rates synergistically determine the morphology of heterochromatin domains in the nuclear interior as well as the periphery. By quantitatively analyzing super-resolution images of chromatin organization, we extract the nucleus-wide distribution of the strength of chromatin-lamina interactions. Along the lamina, we predict the presence of interspersed regions with very strong and vanishing chromatin tethering resulting in a bimodal distribution of chromatin-lamina affinity. We utilize our formulation to examine the evolution of peripheral heterochromatin organization in response to alterations in the cell microenvironment, as well as the effects of epigenetic drugs and pharmacological treatments influencing cell contractility. This information is subsequently utilized to predict the mechanisms mediating these interactions across diverse cellular microenvironments.

Cells adapt to the mechanical properties of their microenvironment by adjusting their internal force-generating machinery, particularly the actomyosin cytoskeleton. These cells enhance their contractility in stiffer microenvironments through mechanochemical signaling pathways [7]. Previous research has demonstrated that the shuttling of epigenetic regulators, such as histone deacetylase HDAC3 [8] and histone methyltransferase EZH2 [9], depends on cellular contractility. In this study, we investigate the formation of LADs (lamina-associated domains) and find that while both enzymes promote methylation, HDAC3, but not EZH2, affects the affinity between chromatin and the lamina. Additionally, inhibiting cellular contractility increases the nuclear levels of HDAC3, leading to an increase in peripheral heterochromatin. Based on these findings, we propose that diseases characterized by softening of the extracellular environment may exhibit increased chromatin condensation, both within the nuclear interior and at the periphery. To test this hypothesis, we analyze cells from healthy donors and patients with tendinosis using our theoretical framework. Our multifaceted integrative framework, which combines theoretical and imaging approaches, provides insights into specific mechanosensitive molecular alterations driving genome organization. Moreover, our framework showcases wider implications for translating these findings into understanding pathological conditions that impact the extracellular environment.

## 2 Materials and Methods

### 2.1 Description of chromatin organization in terms of the hetero and euchromatic phases, and the nucleoplasm

At the mesoscale, chromatin is organized as transcriptionally active euchromatin alongside tightly compacted domains of heterochromatin where genes are predominantly silenced. The heterochromatin domains can occur both in the interior of the nucleus as well as along its periphery as lamina-associated domains (LADs)[10] (Fig 1A). We mathematically model the interior and peripheral compartmentalized organization of chromatin as a far-from-equilibrium dynamic phenomenon governed by the coupled energetics of chromatin-chromatin and chromatin-lamina interactions. The spatiotemporal evolution of chromatin is governed by the kinetics of nucleoplasm diffusion and spatially varying active interconversion of acetylated and methylated histones mediated by epigenetic factors such as histone deacetylases (HDACs).

**Figure 1.**
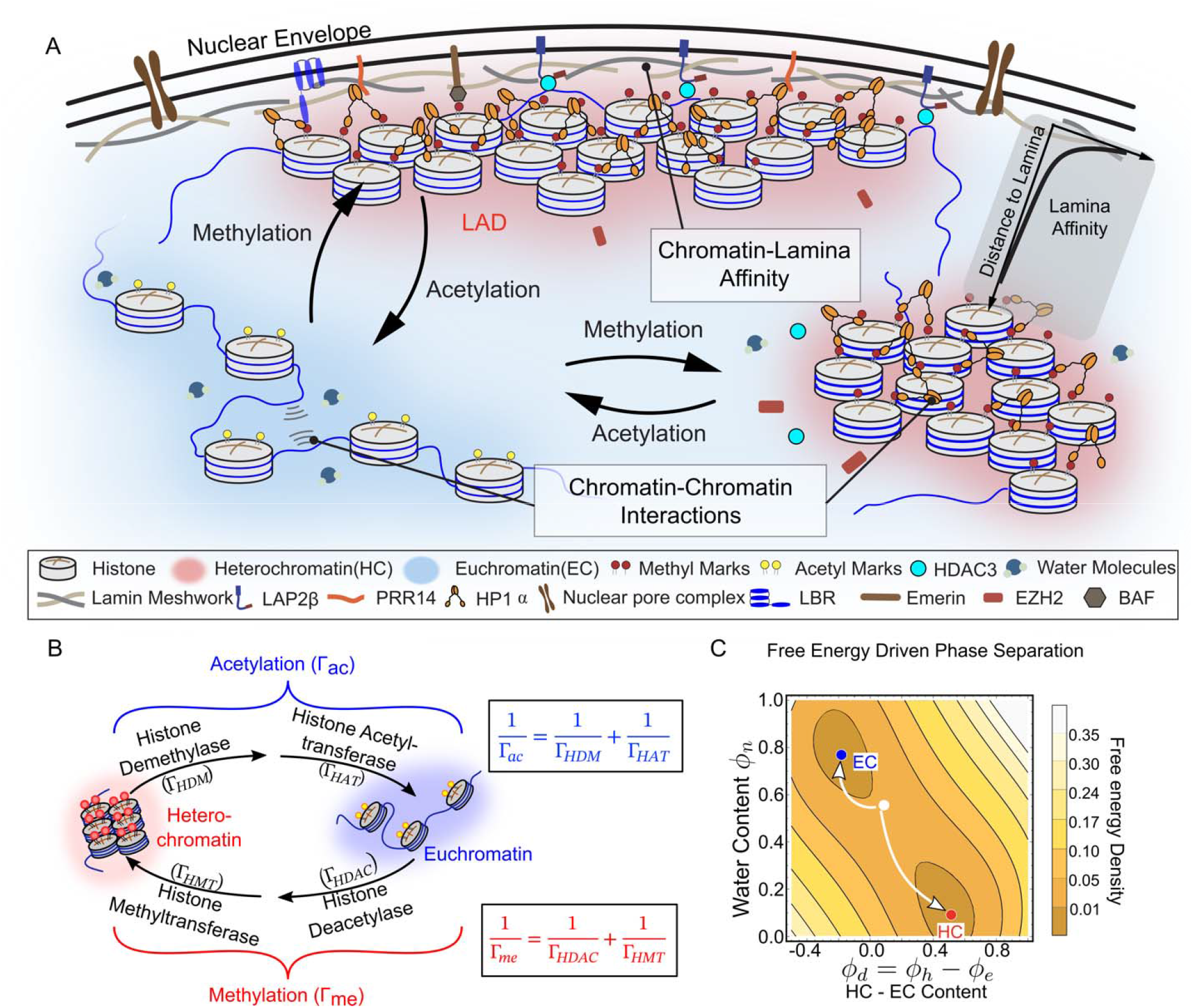
Schematic of the numerical model capturing the formation of interior HC domains and peripheral HC domains. (A) Chromatin-chromatin interactions result in segregation of chromatin into euchromatin and heterochromatin phases. These interactions incorporate chromatin-chromatin interactions, including those mediated by crosslinking molecules such as HP1α, as well as the direct interactions between segments of chromatin. The model also includes the interactions between chromatin and the lamina, mediated by anchoring proteins such as LAP2α/β and LBR. These chromatin-lamina interactions, localized at the nuclear periphery, result in the formation of heterochromatin rich lamina-associated domains (LADs). (B) Epigenetic factors, such as HDAC and HMT, regulate acetylation and methylation reactions that allow interconversion of heterochromatin and euchromatin phases, captured via first-order reaction kinetics. The diffusion of water and epigenetic markers is included in the continuum model. The anchoring of chromatin to the nuclear lamina is also mediated by HDAC3[10, 21]. (C) The contour plot of free energy density shows the two wells (local minima) corresponding to the two stable phases of chromatin – euchromatin (blue) and heterochromatin (red). We schematically show how an initial homogeneous distribution of chromatin (white circle) will evolve towards the two energy wells.

The nucleus comprises three constituents, namely, the two forms of chromatin – hetero (prominently methylated repressive form) and euchromatin (prominently acetylated) – and the nucleoplasm. Since our primary focus is on the key features of chromatin organization, all other nuclear constituents are included in the nucleoplasm. At any given location **x** and time *t*, we define the volume fraction of the three nuclear constituents as *ϕ*_*h*_ (**x**,*t*), *ϕ*_*e*_ (**x**,*t*), and *ϕ*_*n*_ (**x**,*t*), such that they add up to unity, *ϕ*_*h*_ +*ϕ*_*e*_ +*ϕ*_*n*_ =1. Thus, the nuclear composition at any point can be fully described by two independent variables. The local nuclear composition at each point can be equivalently defined in terms of (i) the volume fraction of the nucleoplasm *ϕ*_*n*_ and (ii) the difference between the volume fractions of heterochromatin and euchromatin *ϕ*_*d*_ = *ϕ*_*h*_ − *ϕ*_*e*_. These variables enable a natural definition of the evolution of nucleoplasm and the epigenetic marks between the two forms of chromatin. Furthermore, *ϕ*_*d*_ plays the role of an order parameter, such that *ϕ*_*d*_ > 0 corresponds to the heterochromatic phase, while when *ϕ*_*d*_ < 0, the euchromatic phase.

### 2.2 Driving forces for the spatiotemporal evolution of chromatin

The evolution of chromatin phases is determined by energetic driving forces along with the epigenetic reaction kinetics of histone methylation and acetylation. To capture the energetic driving forces, we construct the local free energy density at any point in space and any instant of time, in terms of the independent variables *ϕ*_*n*_ (*x*,*t*) and *ϕ*_*d*_ (*x*,*t*) as,

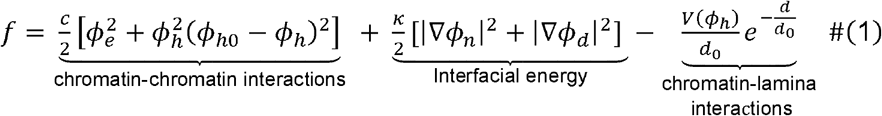

Here the first term represents a Flory–Huggins type of description for the free-energy of chromatin organization and arises from the competition between the enthalpy of nucleosome interactions and the entropy of mixing of the two phases. The enthalpic interactions between proximal nucleosomes could include electrostatic and hydrogen-bonding contributions associated with their epigenetic state. For instance, histone posttranslational modifications can alter the positive charge on histones, thereby modifying chromatin compaction [11].

Furthermore, cross-linking of methylated histones mediated by HP1α[12, 13] can increase chromatin-chromatin interactions and reduce overall enthalpy, leading to an energy minimum corresponding to compacted water-poor heterochromatin phase. On the other hand, the water-rich, loosely packed euchromatin phase increases the local entropy giving rise to second energy minimum. Moreover, differential levels of activity between the transcriptionally active euchromatin phase and the silenced heterochromatic phase have also been shown to alter the depth of the energy wells [14], thus contributing to energetic chromatin-chromatin interactions. Thus, the resulting double-well energy landscape comprises two minima (Fig 1C) corresponding to the euchromatin (*ϕ*_*h*_ = 0)and heterochromatin (*ϕ*_*h*_ = *ϕ*_*h0*_) phases. Physiologically the strength of chromatin-chromatin interactions is order of ∼ 2 − 4*k*_*B*_*T* ∼ 8 − 16 *pN. nm* [15] (details SI section S9). The parameter *c*, which characterizes these chromatin-chromatin interactions per unit volume, is on the order of ∼ 8 × 10 ^−3^ − 16 × 10^−3^ *pN /nm*^2^.

The second set of terms penalizes the formation of sharp gradients in the spatial organization of chromatin (refer SI, Section S1.2) which opposes the bulk energetic contributions from chromatin-chromatin interactions that try to segregate the dissimilar phases. The term *k* (∼0.8 *pN*, estimated in SI section S9) is the increase in the energy due to formation of a unit width of the interface. As *k* increases, there is a greater penalty on formation of sharp interfaces, resulting in more smooth interfaces. The width of the interface *l*_*int*_ is determined by the balance of the interfacial penalty and the bulk energy contributions, such that 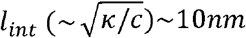 (see SI section S9 for details). We have previously reported that the experimentally observed change in chromatin density between hetero and euchromatin is indeed smooth[16] .

The last term captures the interaction between chromatin and the nuclear lamina, which leads to the formation of LADs along the nuclear periphery. In-vivo, the interactions between chromatin and the nuclear lamina are orchestrated by a diverse set of chromatin anchoring proteins, such as LAP2β, LBR, emerin and PRR14, associated with the inner nuclear membrane and the nuclear lamina[10, 17]. While these interactions can be individually incorporated by capturing the strength of each anchoring protein associated with the lamina and the nuclear membrane, as a first approximation we assume these to be the same.

The function *V* (*ϕ*_*h*_) measures the chromatin-lamina interaction energy per unit surface area of the nuclear envelope and considers all tethering mechanisms collectively contributing to the spatial organization of LADs. The magnitude of strength of chromatin-lamina interactions are dependent on the local chromatin state. In euchromatic phase (*ϕ*_*h*_ = 0) the interactions are mediated by proteins in the lamina that may tether euchromatin and have a strength *V*_*EC*_ = *V*(0).On the other hand, the tethering of methylated histones, mediated by proteins such as LAP2*β* has an interaction strength *V*_*HC*_ = *V*(*ϕ*_*h*0_). We define the preferential interactions of the lamina with heterochromatin over euchromatin as *V*_*LAD*_ = *V*_*HC*_ − *V*_*EC*_. As shown in the SI (section S5.1), the specific choice of *V*(*ϕ*_*h*_)only weakly impacts *V*_*LAD*_. Furthermore, the strength of chromatin-lamina interactions decays exponentially away from the nuclear periphery over a length-scale *d*_0_ comparable to the size of anchoring proteins (∼ 2.5 *nm*) [18]. Physically, the term *V* / *d*_0_ in Eq (1) measures the chromatin-lamina interactions per unit volume of region where the LADs are formed.

### 2.3 Spatiotemporal dynamics of chromatin evolution

To reduce the total free energy of the system, any random initial configuration (white dot) on the energy landscape (shown via a contour plot in Fig 1C) will be driven toward the two energy wells (red and blue dots). The temporal dynamics of energy-driven evolution are determined by the magnitudes of the gradients of the energy landscape, defined as chemical potentials 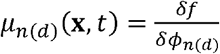 . Here, the functional derivative *δ* is a measure of the change in free energy density with respect to the volume fraction. The spatiotemporal evolution of nucleoplasm is diffusively driven by spatial heterogeneities in its chemical potential *μ*_*n*_ and can be written as,

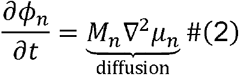

where *M*_*n*_ denotes the mobility of nucleoplasm in the nucleus. Thus, the passive diffusion of the nucleoplasm over time, via Eq 2, reduces the overall free energy of chromatin organization while conserving the net amount of nucleoplasm in the nucleus unless there is exchange of water with the cytosol (as shown in Fig S1) via appropriate boundary conditions, as discussed in the SI section S1.4.

The diffusion of nucleoplasm via Eq 2 captures only the movement of water, without changing the relative locations of acetylated and methylated marks on chromatin, as shown in Fig S1. Epigenetic marks can be actively added to or removed from histone tails via epigenetic reactions mediated by epigenetic factors such as HDACs, allowing interconversion between the heterochromatin and euchromatin (Fig S1). The conversion of euchromatin into heterochromatin phase requires two steps – removal of acetyl group via deacetylation followed by addition of methyl group via methyltransferase (HMT) activity. Thus, the overall rate of histone methylation can be written as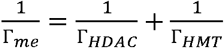 , as shown in Fig 1B. Similarly, the conversion of heterochromatin into euchromatin phase, which involves the activities of histone demethylase (HDM) and histone acetyltransferase (HAT) is 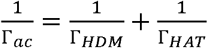 . The second term in Eq 3 represents these nonconservative reaction kinetics.

Additionally, the reaction kinetics can also be influenced by chromatin-chromatin interactions between spatially proximal nucleosomes. For instance, conversion of heterochromatin into euchromatin might be energetically less favorable in a heterochromatin-rich neighborhood than in a euchromatin-rich neighborhood (further discussion in the SI, section S1.4). The reaction-kinetic contributions arising from these neighborhood interactions can be shown to aptly mimic (see SI section S1.4 for the theoretical derivation) a diffusion-like evolution of epigenetic marks determined by neighborhood-dependent reaction kinetics. The diffusion-like and reactive kinetics together determine the spatiotemporal evolution of epigenetic marks as,

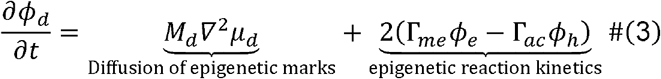

where *M*_d_ is the effective mobility of epigenetic marks in the nucleus (see SI section S1.3). Note that Γ_*me*_ and Γ_*ac*_ are not constant but spatially dependent on the non-homogeneous distribution of epigenetic regulators [9, 19] such that Γ_*me*_ = Γ_*me*_ (**x**,*t*) andΓ_*ac*_ = Γ_*ac*_ (**x**,*t*).

Eq 1-3 can be numerically solved to determine the spatiotemporal organization of chromatin in the nucleus. For the numerical solution, we rescale the equations using the intrinsic length and time scales, as described in the SI (section S1.5). A description of the model parameters, as well as the initial and boundary conditions used for the numerical solution is provided in the SI section S8.

### 2.4 Super-resolution STORM imaging and localization analysis of chromatin structures

STORM images were obtained from previously published work [9, 20] and re-analyzed using the theoretical framework developed here.

#### Cell culture and immunostaining

Human mesenchymal stromal cells (hMSCs) were isolated from fresh bone marrow obtained from human donors[22]. The cells were then plated and expanded on tissue-culture plastic in α-modified essential medium (α-MEM) supplemented with 10% fetal bovine serum (FBS), 1% penicillin–streptomycin, and 5 ng/ml basic fibroblast growth factor at 37°C and 5% CO2 until the colonies reached 80% confluency. Subsequently, the cells were stored in liquid nitrogen using a freezing medium composed of 95% FBS and 5% dimethylsulfoxide (DMSO). Throughout the expansion process, all hMSCs were cultured in standard growth medium consisting of α-MEM supplemented with 10% FBS and 1% penicillin– streptomycin. Human Tenocytes (hTCs) were isolated from the finger flexor tendon tissues of both young individuals and patients diagnosed with tendinosis[23]. The cells were then plated in basal growth medium, which consisted of high-glucose DMEM medium supplemented with 10% penicillin–streptomycin, L-glutamine, and 10% FBS.

hMSCs and hTCs were fixed using methanol-ethanol (1:1) at -20°C for 6 minutes, followed by blocking with a solution of 10%(w/v) bovine serum albumin (BSA) in phosphate-buffered saline (PBS) for 1 hour[3]. Subsequently, the cells were subjected to overnight incubation at 4°C with a 1:50 dilution of histone H2B anti-rabbit antibody (ProteinTech, #15857-1-AP). After thorough PBS washing, the samples were incubated with secondary antibodies labelled with Alexa Fluor 405 - Alexa Fluor 647[3].

Human fibroblasts (hFB) were grown in growth media (DMEM medium, 10% FBS, 1× non-essential amino acids, 1× Penicillin/Streptomycin and 1× GlutaMax) supplemented with 5 μM ethynil-deoxy-cytidine (EdC) for 96 hours[20]. The cells were fixed with 4% paraformaldehyde (PFA) diluted in PBS for 15 minutes at room temperature. Subsequently, they were permeabilized with 0.3% (v/v) Triton X-100 in PBS for 15 minutes at room temperature. Afterward, the cells were blocked using a solution containing 10% BSA (w/v) and 0.01% (v/v) Triton X-100 in PBS.

#### Confocal Airyscan imaging

Multi-channel confocal Airyscan images of hMSCs were acquired using the Zeiss LSM 900 with Airyscan 2 commercial super-resolution imaging platform (x63 oil immersion objective). The HDAC3 (Cell Signaling Technology, #3949, dilution 1:50) reporter dye Alexa Fluor 647 (Abcam, Cat: ab150115, dilution 1:50) was excited using 640 nm laser set at 5% power. The selective staining of DNA and F-actin were done by DAPI (Thermo Fisher Scientific, Cat: D1306) and conjugated Alexa Fluor 568 (Thermo Fisher Scientific, Cat: A12380). The DAPI and Phalloidin reporter dye were excited with 405 nm, and 560 nm laser at 2% power, respectively.

#### Super–resolution Imaging

Super-resolution imaging-based visualization of immunostained H2B was utilized to observe the detailed meso-scale organization of chromatin in the cell nuclei. Super-resolution images were acquired using the ONI (Oxford NanoImager) commercial STORM imaging platform. The imaging system was equipped with a ×100, 1.4 numerical aperture oil-immersion objective and a sCMOS Hamamatsu Orca Flash camera. To ensure optimal photo switching of Alexa Fluor 647, the imaging buffer followed standard guidelines[24], consisting of 10 mM cysteamine MEA in GLOX Solution: 0.5 mg ml^-1^ 1-glucose oxidase, 40 mg ml^-1^ 1-catalase and 10 % glucose in PBS. The reporter dye (Alexa Fluor 647) was excited using a 640 nm laser set at 40% power. Gradually, the power of the 405 nm laser was increased to reactivate Alexa Fluor 647. The camera’s exposure settings were configured to 15 ms, and a total of 30,000 frames were collected for each image using the ONI software. Additional details can be found in [9].

Super-resolution imaging of EdC labeled DNA in hFb cells were acquired using a commercial N-STORM microscope (Nikon) equipped with a CFI HP Apochromat TIRF 100 × 1.49 oil objective, an iXon Ultra 897 camera (Andor), and a Dual View system (Photometrics DV2 housing with a T647lpxr dichroic beam splitter from Chroma). A 647 nm laser was employed to excite the DNA labeled with AlexaFluor 647, with a power density of approximately 3 kW/cm. Alexa 647 was progressively reactivated with increasing 405 nm laser power during acquisition, up to a maximal power density of 0.020 kW/cm^2^. The imaging buffer consisted of 100 mM Cysteamine MEA, 5% glucose, 1% Glox, and 0.75 nM Imager strand (I2-560 Ultivue) in Ultivue Imaging Buffer. Additional details can be found in [20].

The subsequent post-analysis of STORM image localization was performed using custom-written MATLAB codes[9].

#### Identification and quantification of heterochromatin domains

MATLAB was utilized for the analysis of STORM images using an adapted Voronoi tessellation-based segmentation method[25] to construct Voronoi polygons of each fluorophore. This approach assigns a Voronoi polygon to each localization, where the polygon size is inversely proportional to the local localization density[20]. For each nucleus, the spatial distribution of localizations is represented by a collection of Voronoi polygons, where smaller polygon areas indicate higher density regions. The nuclear area is computed by summing the Voronoi polygon areas, excluding edge polygons due to their disproportionately large size resulting from edge effects. Localization density (total localizations per nuclear area) varies across nuclei due to differing mechanical and chemical treatments. To compare chromatin organization changes from different mechanical/chemical treatments, the localization density of each nucleus is normalized to its mean Voronoi polygon area, yielding a standard density unit. The inverse of the polygon area forms the reduced Voronoi density. Reduced Voronoi densities across nuclei and treatments were pooled and visualized as cumulative distribution plots. A threshold was set so that the 0-30 percentile of the density distribution represents sparse chromatin, while the 31-70 percentile represents dense chromatin. The Density-Based Spatial Clustering of Applications with Noise (DBSCAN) algorithm was then used to cluster neighboring and connected Voronoi polygons into distinct domains, ensuring at least three localizations per cluster.

#### Identification and quantification of lamina-associated domains

We further distinguished LADs from interior heterochromatin domains based on their proximity to the nuclear lamina. For each heterochromatin cluster, we quantified the distance of each localization to nuclear envelope. If any localization lies within the predefined threshold distance, we classified that cluster as a LAD. The threshold was set at 2.5% of nuclear radii from nuclear envelope.

To quantify the thickness of LADs, we divided the nuclear boundary into n segments. Within each segment, we computed the cumulative area covered by LADs and divided it by the segment length to determine the thickness of the LAD domain. In our analysis, we selected n=50 as the number of segments. However, we also conducted tests up to n=100 and observed no significant changes in the results.

Note that the quantification data obtained from STORM images, coupled with the theory, will be used to extract the genome-wide distribution of chromatin-lamina interactions and methylation rates. In the following section, we will first discuss the scaling relation predicated by theory, followed by the integrative approach used to obtain the biophysical parameters.

## 3 Results

### 3.1 Interplay of epigenetic reactions, histone diffusion and chromatin-lamina interactions determine the size of interior domains and LADs

#### Model predicts the formation of stable interior domains and LADs observed using super-resolution imaging

The mathematical model developed in Section 2 (Eq 1-3) is numerically solved to predict chromatin distribution in the nucleus (details in the SI section S8). We observe the segregation of water-poor, condensed heterochromatin region (corresponding to the heterochromatic energy well (red) in Fig 1C from a distinctly water-rich euchromatin region (euchromatic energy well (blue) in Fig 1C). The nascent nucleated domains of heterochromatin grow, attaining a quasi-periodic distribution both in the interior and at the nuclear periphery, as shown in Fig 2A, which is stable against further coarsening. Consistent with our simulations, histone density mapping of nuclear super-resolution STORM imaging on 2D substrates [9] reveals that chromatin organizes into compacted heterochromatin, characterized by high histone density, which is clearly distinguishable from the low-density euchromatin region (Fig 2B). We observe heterochromatin domains localized within the interior of the nucleus and along the nuclear periphery (LADs) in both simulations and high-resolution STORM images (Fig 2A, B). Our model predicts that at steady-state, the size distribution of the heterochromatin domains in the interior of the nucleus shows a characteristic mean (Fig S11), in excellent agreement with the size-distribution of condensed chromatin domains observed via STORM imaging (Fig 3B). Importantly, the model also predicts that near the nuclear periphery, LADs form either discrete domains or continuous layers (Fig 2a) and show a size distribution that peaks at a characteristic length-scale, as shown in (Fig S11). The characteristic size-scale of the LADs is also confirmed from super-resolution imaging of cells (Fig 3C).

**Figure 2.**
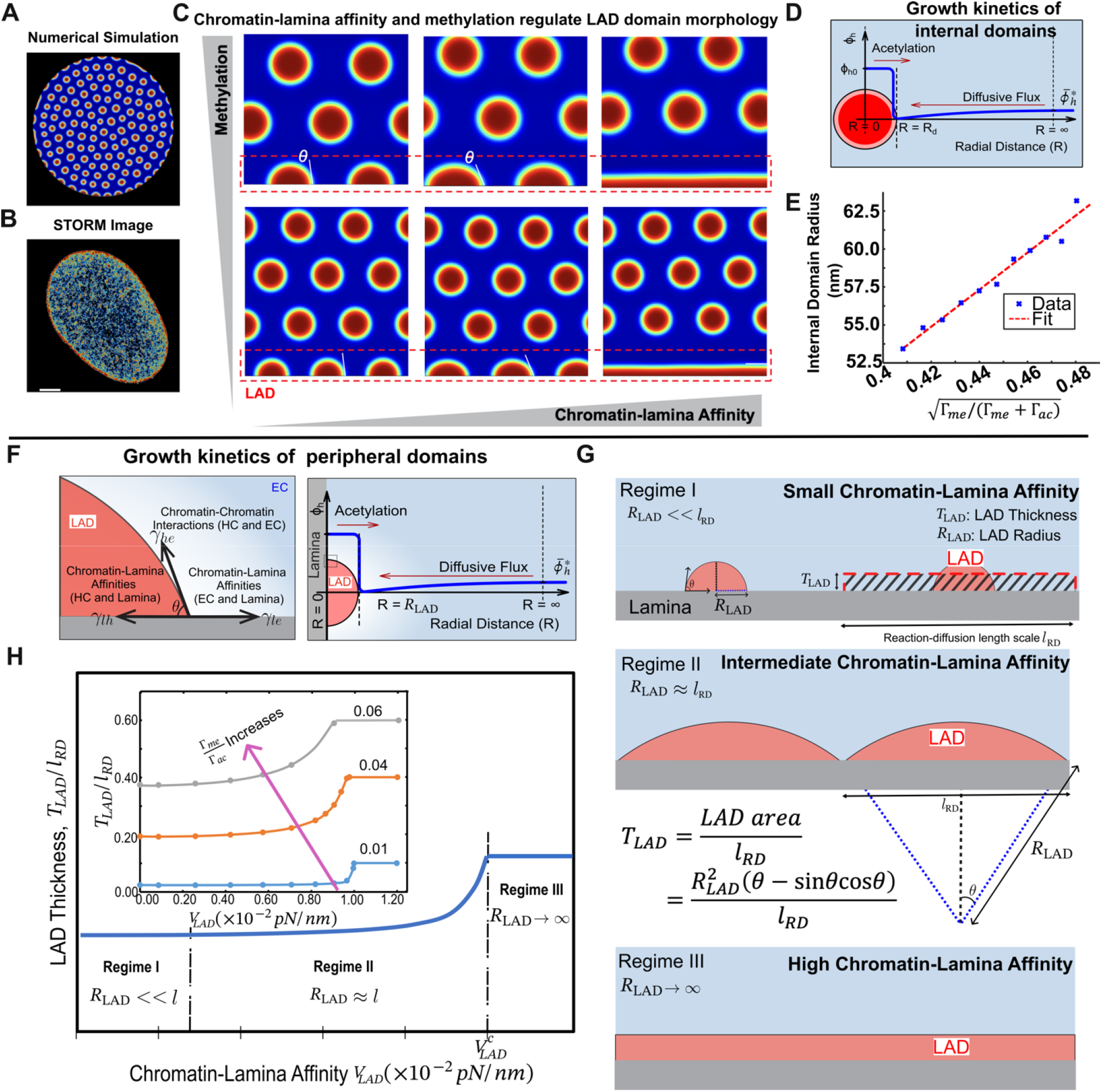
The morphology of interior heterochromatin domains and LADs is regulated by the methylation levels and the strength of chromatin-lamina affinity. (A) Steady-state chromatin organization predicted by numerical simulations showing phase separation into compacted heterochromatin domains (red) and loosely packed euchromatin domains (blue) and the formation of heterochromatin-rich LADs at the nuclear periphery. (B) Representative Voronoi density rendering of H2B STORM images, color-coded to indicate density levels (blue for low density and red for high density). (C) Numerical simulations showing the synergistic effects of chromatin-lamin affinity and methylation rates on LADs and inner heterochromatin domain morphology. At low levels of chromatin-lamina affinity, as the methylation level increases, the radius of the discrete LADs increases at a fixed value of the contact angle . The morphology of LADs is determined by both the level of methylation and chromatin-lamin affinity. Low affinity results in the formation of isolated LADs, while at greater levels of affinity LAD spread along the lamina. (D) Kinetic balance determines the steady-state size of interior HC domains. The diffusion of methylate histones from the EC region towards the HC domains and histone methylation drive their growth, counteracted by the opposing kinetics of histone acetylation. (E) The scaling relationship between the size of interior HC domains and the level of methylation. (F) (Left) At the interface of HC-rich LAD, EC-rich environment, and lamina, a balance of interfacial interaction between the two phases of chromatin (*γ*_*he*_ ) and the strength of chromatin-lamina affinities (*γ*_*lh*_ ) gives rise to a stable morphology of LADs, which have a characteristic contact angle with the lamina(*θ*). (Right) The formation of LADs in steady state requires a balance between diffusion-driven heterochromatin influx (into the LAD) and acetylation-driven conversion of heterochromatin into euchromatin. (G) Schematics illustrating the morphology of LAD at varying levels of chromatin-lamina affinity. (H) The scaling relationship between the thickness of LAD and chromatin-lamina affinity at a given methylation level. The inset shows the scaling relation at varying methylation levels.

**Figure 3.**
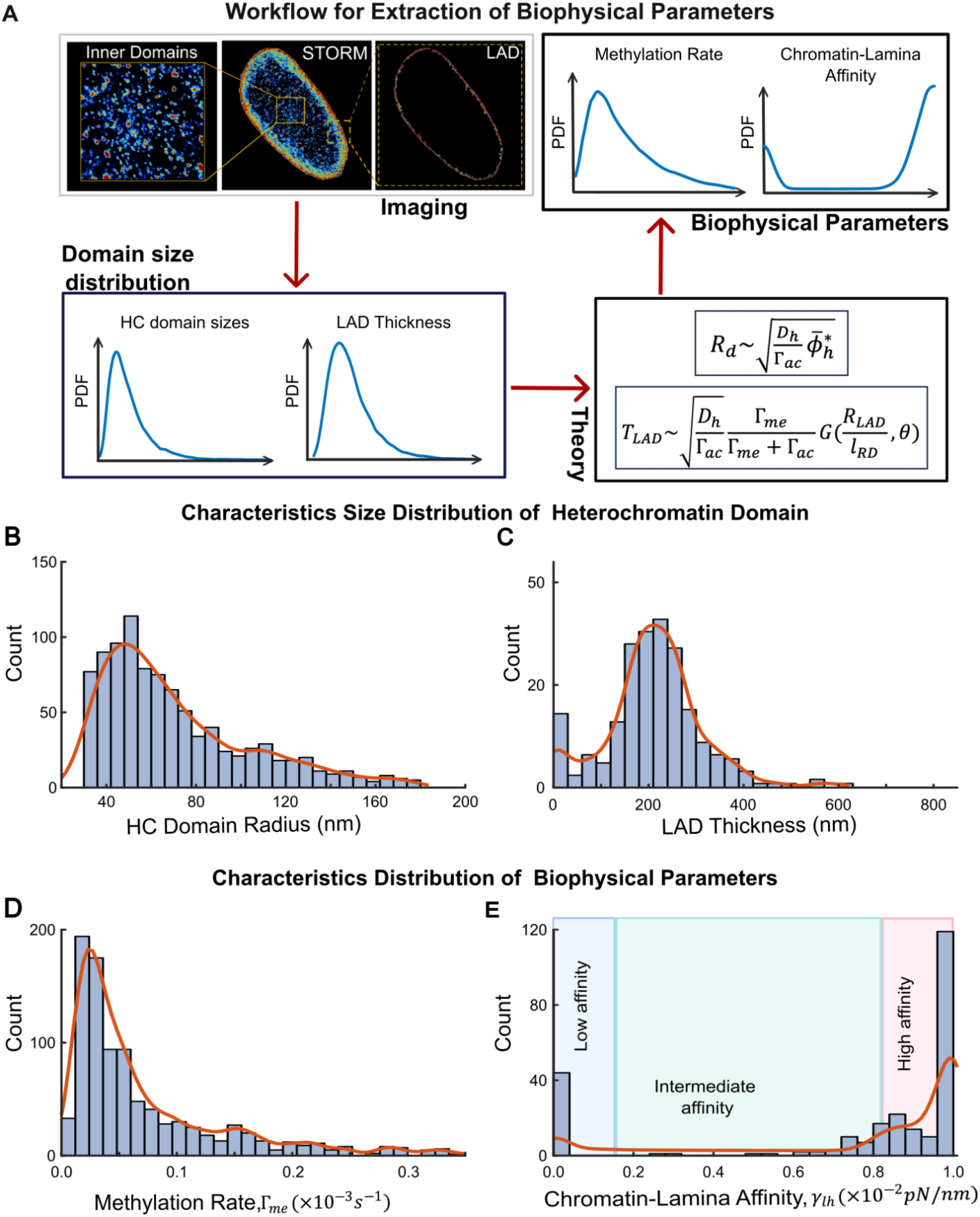
(A) Workflow of the theoretical framework for extracting methylation rates and chromatin-lamina affinity by integrating super resolution STORM images of DNA with theoretical analyses. Within the interior of the nucleus, STORM image analysis gives the distribution of heterochromatin domain radii, and at periphery, it provides the distribution of LAD thickness. The combination of these distributions with the theoretical framework predicts the corresponding distribution of methylation rates and the chromatin-lamina affinities. The characteristic distribution of interior HC domains sizes *R*_*d*_ (B), LAD thickness *T*_*LAD*_ (C) obtained from STORM images of control nuclei. The theoretical framework then predicts the nuclear wide distributions of (D) histone methylation rate Γ_*me*_ and (E) chromatin-lamina interaction affinity *V*_*LAD*_. Notably the distribution of *V*_*LAD*_ is bimodal. The red curve in all plots shows the smoothed density plot. All source data are provided as a Source Data file.

Upon compaction, chromatin can be partitioned into heterochromatin domains within the nucleus interior or LADs along the nuclear periphery. Thus, the total heterochromatin content of the nucleus is shared between the interior domains and LADs such that,

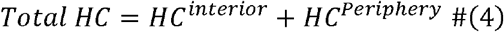

In Eq 4, ‘Total HC’ is calculated as the average heterochromatin content in the nucleus 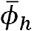 times the total nuclear area. In the presence of epigenetic reactions, the average amount of heterochromatin is determined by the epigenetic reaction rates, 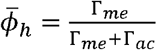 (detailed derivation in SI section S2); the average amount of heterochromatin in the nucleus increases with the repressive methylation rate. Note here that the repressive methylation rate comprises deacetylation of histones via HDACs and the addition of repressive methyl groups (e.g. H3K9me2/3 and H3K27me3) by HMTs (as shown in Fig 1B). Next, we determine the biophysical parameters that govern the size of heterochromatin domains in the interior and at the periphery of the nucleus.

#### Sizes of the interior heterochromatin domains are determined by the balance between epigenetic reactions and diffusion kinetics

We begin by considering the growth of the heterochromatin domains in the interior of the nucleus, away from the periphery. As the overall levels of HC and EC phases in the nucleus 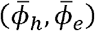 do not lie in either of the energy wells (white circle in Fig 1C), the free energy can be lowered by nucleating heterochromatin domains (*ϕ*_*h*_ = *ϕ*_*h*0_, red circle in Fig 1C) surrounded in the immediate vicinity by euchromatin (*ϕ*_*h*_ = 0, blue circle in Fig 1C). Far from the nucleated domain, the concentration of heterochromatin 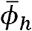 is determined by epigenetic reactions. However, when there is significant peripheral sequestration of methylated histones, the far-field concentration of heterochromatin 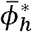 (Fig 2D) decreases from the average concentration 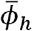. The spatial gradient in chromatin concentration (blue curve in Fig 2D) leads to a diffusive flux of methyl marks (Fig 2D) towards the domain, driving its growth (refer SI section S3 for quantitative derivation). Inside the domain (*ϕ*_*h*_ = *ϕ*_*h*0_), acetylation converts the heterochromatic methylated histones into euchromatic form, which are then driven outward due to their preference for like-marked histones. As depicted in Fig 2D, the effective outflux of methylated histones opposes the diffusion gradient-driven influx of methyl marks. At steady state, the balance between the opposing fluxes stabilizes the domain (refer SI section S3 for derivation) with its radius scaling as,

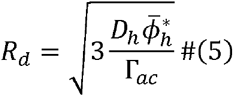

A detailed description of the steps involved in the growth and stabilization of the inner heterochromatin domains is provided in the SI (section S3), along with the derivation of Eq 5. The size scale of the interior heterochromatin domains predicted theoretically by Eq 5 makes it immediately apparent that the balance between the heterochromatin influx due to diffusion kinetics (the 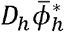 term) and the acetylation driven outflux (the Γ_*ac*_ term), independent of energetic considerations, determines the size of the interior heterochromatin domains.

Next, we relate the far-field heterochromatin concentration 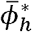 to the average heterochromatin content in the nucleus, 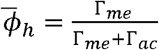 via Eq 4 and 5 (for a detailed derivation see SI Section S4) as,

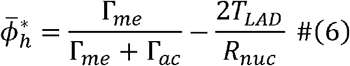

Here,*T*_*LAD*_ is the size of heterochromatin accumulating at the lamina and *R*_*nuc*_ is the radius of the nucleus (see SI section S5.2). Note that when the LAD thickness is very small (*T*_*LAD*_ ≪ *R*_*nuc*_), the influence of LADs on the interior domains is insignificant, and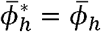. The additional term 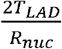 in Eq 6 arises from interdependence of interior and peripheral domains via Eq 4.

Consistent with our theoretical analysis, simulations show that as methylation increases, the heterochromatin domains grow, and vice versa (Fig 2C). As a further validation, the numerically observed dependence of mean size of domains on epigenetic reactions also closely follows the theoretically predicted scaling (Fig 2E). Next, we determine the size scaling of the LADs at the nuclear periphery with respect to the histone methylation rate and the strength of chromatin-lamina interactions.

#### LAD shapes are determined by chromatin-lamina affinity while their sizes are regulated by epigenetic reactions

The presence of chromatin-anchoring proteins such as LAP2*β* , HDAC3 and LBR [6, 10, 26] that sequester chromatin to the lamina is captured in our model through the chromatin-lamina interaction energy per unit surface area of nuclear lamina, *V*(*ϕ*_*h*_ )Eq 1). Notably, the chromatin-lamina interaction energy reflects the collective impact of several chromatin-anchoring proteins, and any alterations in it signify either a change in the number or affinity of these proteins. These interactions lead to the localization of chromatin to the periphery, resulting in the formation of a euchromatin-heterochromatin (he) interface, a lamina-heterochromatin (lh) interface and a lamina-euchromatin (le) interface, as shown in Fig 2F (left panel). The surface tension *γ* of each of these interfaces is defined as the energetic cost of forming a unit area of the interface. The balance of the three surface tensions governs the extent of peripheral localization of chromatin, analogous to the wall-wetting phenomenon described by the well-known Young’s equation, *γ*_*lh*_ = *γ*_*he*_ *cosθ* + *γ*_*le*_, where *θ* is the contact angle of the LAD with respect to the nuclear lamina, which determines the morphology of the LAD. Here, *γ*_*lh*_ and *γ*_*le*_ represents the surface tension at lamina-heterochromatin and lamina-euchromatin interface respectively, *γ*_*he*_ denotes the surface tension corresponding to heterochromatin-euchromatin interface (Fig 2F). Thus, the contact angle of LAD with respect to the nuclear lamina *θ*, and consequently the LAD morphology, is determined by the energetic competition between chromatin-chromatin interactions and chromatin-lamina interactions as,

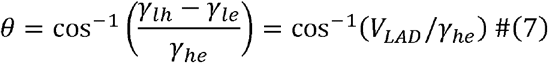

Here, *V*_*LAD*_ / *γ*_*he*_ is the strength of chromatin-lamina interactions relative to the strength of chromatin-chromatin interactions, where *V*_*LAD*_ (= *V*_*HC*_ − *V*_*EC*_ ∼ *γ*_*lh*_ − *γ*_*le*_) quantifies the preferential interaction of lamina with heterochromatin over euchromatin (detailed derivation in SI section S5.1). Thus, using Eq 7, the morphology of the LADs can be directly used as a readout of the chromatin-lamina affinity. However, when quantifying LADs observed experimentally via STORM imaging (Methods section 2.4), resolution limits can introduce errors in the measurement of individual contact angles for each LAD. Thus, we define the average thickness of LAD *T*_*LAD*_ as its height averaged over its span as shown in Fig 2G (top panel, mathematically defined in the SI, Section S5.2). As for the interior domains, balancing the diffusive flux of methyl marks into the LADs and the conversion of methyl marks to acetyl marks in the LADs, we show in SI section S5.2 that the average LAD thickness can be related to the reaction rates, diffusion constant and contact angle through the relation,

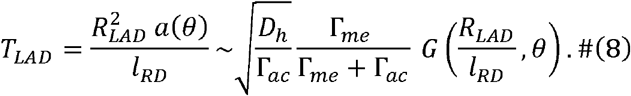

Here, the function *a*(*θ*) quantifies the area of the LAD at steady state (Eq S34), *l*_*RD*_ (∼ 300 *nm*) is the characteristic length scale determined by reaction-diffusion kinetics (SI section S9) and *G*(*R*_*LAD*_, *θ*) is a non-linear function that measures the total diffusive influx of methyl marks (inclusive of the competition between interior domains and LADs) and is contingent on both the LAD-lamina contact angle *θ* and the LAD radius *R*_*LAD*_. The precise dependence of LAD thickness *T*_*LAD*_ on histone methylation rate Γ_*me*_ and chromatin-lamina affinity relative to chromatin-chromatin interaction strength *V*_*LAD*_ (∼ *γ*_*he*_ cos *θ*)can be obtained numerically by determining the shape of LAD when the heterochromatin influx and outflux are exactly balanced (for the simulation methodology and numerical results refer SI section S8) as shown in Fig 2H. We find that as the histone methylation rate increases (or conversely, the acetylation rate decreases) the LAD thickness increases monotonically, for a given chromatin-lamina affinity as shown in the inset to Fig 2H. On the other hand, the dependence of *T*_*LAD*_ on *V*_*LAD*_ is non-linear and exhibits three distinguishable regimes with increase in the strength of chromatin-lamina interactions, as described in Section S5 (denoted in Fig 2H). In Regime I, vanishing chromatin-lamina affinity leads to sparsely distributed, bead-like heterochromatin domains with minimal LAD thickness (Fig 2G, top). As affinity increases in Regime II, these HC domains spread and elongate, increasing in thickness and decreasing the spacing between them (Fig 2G, middle). In Regime III, high affinity results in a near-continuous HC layer along the lamina (Fig 2G, bottom). In all three regimes, increasing methylation rate enhances heterochromatin content in the nucleus, amplifying LAD thickness. In the three regimes, we theoretically and numerically confirm that the thickness of the LADs follows distinct scaling with epigenetic reactions and chromatin-lamina affinity as discussed in detail in the SI (Section S7).

To summarize, we have theoretically and numerically determined the length scales of the heterochromatin domains within the nuclear interior (Eqs 5, 6) and at the periphery (Eqs 7, 8) taking into account their interdependence (Eq 4). We find that the heterochromatin domain sizes are self-consistently determined by the biophysical parameters governing epigenetic reaction kinetics ( ) and the strength of chromatin-lamina interactions . Thus, using the distribution of the interior domain radius and LAD thickness obtained from the quantification of STORM images (discussed in section 2.4), Eqs 5-8 can be solved simultaneously to determine the nucleus-wide distribution of biophysical parameters, as we discuss next.

### 3.2 Extracting biophysical parameters from super-resolution images using theory

In this section, we use the theoretical relations derived in Section 3.1 to extract the epigenetic reaction rates of methylation Γ_*me*_ and the strength of the chromatin-lamina interactions relative to chromatin-chromatin interactions *V*_*LAD*_ (∼ *γ*_*lh*_ *γ*_*le*_ ∼ *γ*_*he*_ cos *θ*) from STORM images. The intranuclear environment is inherently heterogeneous due to spatial variations in epigenetic regulators, chromatin anchoring proteins, and the presence of other nuclear constituents. For instance, the Lamin B proteins, which are involved in heterochromatin association with the lamina, are heterogeneously distributed in the nuclear envelope [10]. Additionally, nuclear membrane structures such as nuclear pore complexes[27] may exhibit relatively weaker binding affinity for chromatin segments. Similarly, spatial variations in the availability of epigenetic factors like HDAC (Fig 6F, 7A) [28] can result in heterogeneous histone methylation rates.

We adopt a two-step procedure (Fig 3A) to extract the distribution of histone methylation rate Γ_*me*_ and strength of chromatin-lamina interactions relative to the chromatin-chromatin interactions *V*_*LAD*_ from STORM images. The theoretical expressions derived in Section 3.1 (which assume homogeneous histone methylation rate and chromatin-lamina affinity) are taken to hold locally in the interior and the nuclear periphery. To visualize the spatial chromatin organization within cells, we generate nuclear-wide H2B density heatmaps via Voronoi tessellation-based segmentation of super-resolution images [9, 20]. By analyzing these images (as described in Section 2.4), we obtain the statistical distributions of the interior domain radii *R*_*d*_ and the LAD thickness *T*_*LAD*_ . The obtained interior domain and LAD sizes are locally related to the methylation rate Γ_*me*_ via Eqs 5 and 6, allowing us to extract the nucleus-wide distribution of Γ_*me*_. Using the extracted distributions of _Γ*me*_ and LAD thickness, we use Eq 8 (numerically depicted in Fig 2H) to deduce the distribution of chromatin-lamina affinity relative to chromatin-chromatin interaction strength *V*_*LAD*_ along the nuclear periphery.

As a second step, we validate that the distributions of _Γ*me*_ and *V*_*LAD*_ extracted using the theoretical framework are accurate despite the simplifying assumptions involved in the theoretical derivation of Eqs 4-8. The mean values of the extracted parameters Γ_*me*_ and *V*_*LAD*_ are used as inputs to simulate chromatin organization in the nucleus via numerical solutions of phase-field equations (Eqs 1-3) using COMSOL Multiphysics. The pharmacological or biophysical perturbation driven changes in the distributions of biophysical parameters, extracted from the theoretical framework, are incorporated into the simulation by a proportionate change in the mean of the parameters Γ_*me*_ and *V*_*LAD*_. By quantifying the simulation predicted chromatin reorganization due to changes in parameters we assess the relative changes in the LAD and interior domain size-scales, validating these changes against those observed in cells. Notably, numerical simulations allow the spatially heterogeneous biophysical parameters, spatially sampled from a normal distribution with a mean comparable to that obtained from parameter extraction. Furthermore, the heterochromatin domains in the simulation can freely interact with their neighbors, reflecting their intrinsically emergent behavior.

Thus, our theory-based image analysis framework leverages the mechanistic principles regulating chromatin organization to utilize super-resolution imaging of chromatin as a readout of the underlying biophysical parameters.

### 3.3 Chromatin-lamina affinity extracted from STORM images shows a bimodal distribution

We first implement the integrated theoretical framework outlined in section 3.2 to analyze chromatin organization in hMSC nuclei cultured on glass under control conditions [9], thereby extracting the distributions of Γ_*me*_ and *V*_*LAD*_. The quantification of heterochromatin domain size scales reveals a skewed distribution of the interior domain radii *R*_*d*_ (Fig 3B, Fig S13) and LAD thickness *T*_*LAD*_ (Fig 3C, Fig S13). Across all nuclei the distributions show a comparable characteristic mean (Fig S13). Next, we employ the theoretical relations between the heterochromatin domain size scales and the biophysical parameters methylation rate Γ_*me*_and chromatin-lamina affinity *V*_*LAD*_ for both the interior domains and the LADs (Eq 5-8). The distribution of Γ_*me*_across all control-treated nuclei exhibits a similar skewness (Fig 3D, Fig S13) with a comparable mean value (Fig S13).

Notably, the distribution of chromatin-lamina affinity *V*_*LAD*_ is bimodal, with two distinctive peaks at very low and very high affinities (Fig 3E). The peak associated with vanishing chromatin-lamina affinity likely corresponds to the regions along nuclear periphery deficient in chromatin binding proteins, such as nuclear pore complex regions which are not known to interact strongly with heterochromatin[29, 30]. Spatially, regions with low chromatin-lamina affinity typically correspond to lamina regions with sparse H2B localizations. The second peak occurs at a very high chromatin-lamina interaction strength, comparable to the chromatin-chromatin interactions, attributable to the presence of chromatin anchoring proteins with a strong affinity for methylated histones. In STORM images, such regions spatially correlate with dense H2B localizations, and high LAD thickness along the lamina. Between the two peaks, we observe a spectrum of chromatin-lamina affinities where the chromatin-lamina interactions are weaker but still nonzero.

The predicted spectrum of chromatin-lamina affinities aligns with a range of chromatin-LaminB1 interaction strengths observed experimentally using LaminB1-chromatin immunoprecipitation (LB1 ChIP) in multiple cell types [31]. Regions along the chromatin polymer exhibiting strong LaminB1 association are classified as Type 1 LADs, resembling the peak of high chromatin-lamina affinity we observe. Furthermore, as we predict, these LADs have highly compacted chromatin apparent from their very low chromatin accessibility [31]. Regions of chromatin polymer exhibiting a weaker (but nonzero) LaminB1 association, with marginally increased chromatin accessibility are classified as Type 2 LADs [31] and resemble the range of low to intermediate chromatin-lamina affinities we observe.

In the subsequent sections, we will examine chromatin reorganization following known pharmacological treatments and biomechanical stimuli. We use the STORM images with our theoretical framework to predict perturbations in the spatial distribution of the histone methylation rate and chromatin-lamina affinity. Our generalized framework remains blind to the specific treatments the nuclei undergo, ensuring that the analysis is not influenced by prior knowledge across datasets. Using numerical simulations, we then confirm that the extracted distributions of the biophysical parameters do indeed drive the observed chromatin reorganization.

### 3.4 Model captures the differential LAD regulation by distinct pharmacological treatments

Next, we comparatively analyze chromatin organization in control nuclei and those with pharmacologically perturbed biophysical parameters. From our theoretical findings in Section 3.1, we see that LAD morphologies are regulated synergistically by histone methylation and chromatin-lamina interactions. By targeting the epigenetic reactions, we previously inhibited the activities of enhancer of zeste homologue 2 (EZH2, a histone methyltransferase) via GSK343 treatment (Fig 4A)[32, 33], and histone deacetylase (HDAC) via pan-HDAC inhibitor trichostatin A (TSA, Fig 5A)[3, 20, 34]. Since histone deacetylase and methyltransferase contribute to the overall rate of histone methylation in our model (Fig 1B), we extract the resulting change in Γ_*me*_. Moreover, HDAC3 also interacts with the inner nuclear membrane protein LAP2*β* in tethering LADs [21]. Hence, we hypothesize that TSA treatment may also alter chromatin-lamina affinity. Thus, the comparison of differential biophysical parameter modulations after GSK and TSA treatment serves to validate the accuracy of our parameter extraction procedure.

**Figure 4.**
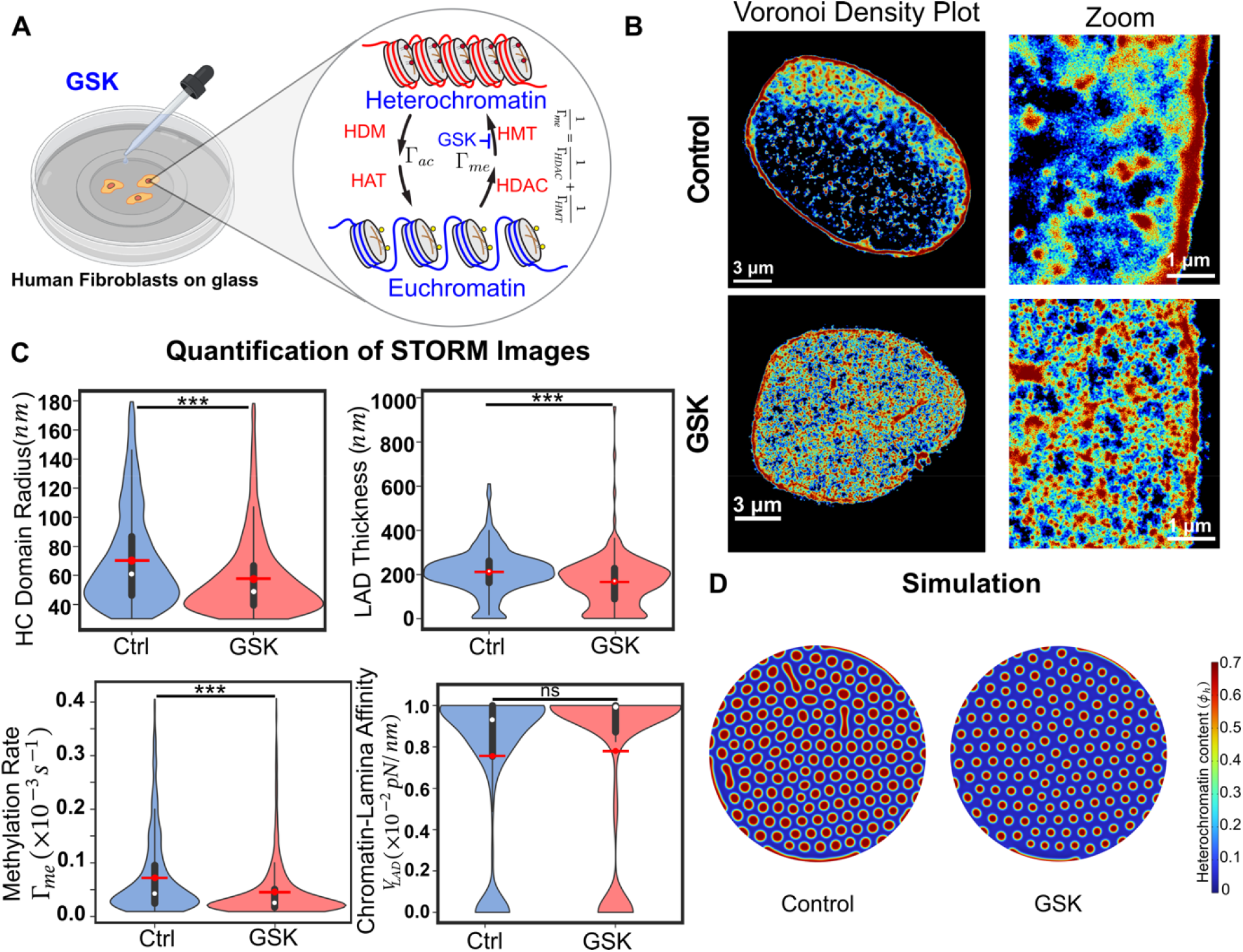
GSK treatment inhibits histone methyltransferase EZH2 resulting in smaller LADs. (A) A schematic showing the role of GSK, which inhibits histone methyltransferases (HMTs) and induces chromatin decompaction. (B) Super-resolution STORM images revealing the reorganization of chromatin upon GSK treatment. (Left panels) Voronoi density plots representing the extent of spatial compaction of chromatin in control (top) and GSK treated nuclei (bottom) along with a zoomed-in view (right panels). (C) Quantification of STORM images in the control an GSK treated nuclei showing the distribution of interior domain radii (left top) and LAD thickness (right top). Distributio of the corresponding methylation rate and the chromatin-lamina affinity (bottom panels) obtained by our integrativ theoretical framework. After GSK treatment, the methylation rate decreases although there is no effect on chromatin-lamina affinity (Control nuclei: n=5; GSK-treated nuclei: n=4; unpaired two tail test, *p<0.05, **p<0.01, ***p<0.001). All violin plots show a symmetric kernel density estimate (outline), the quartiles (black boxplot), the median (white dot), and the mean (red dot with line). (D) Numerical predictions of the size of heterochromatin (in red) in the control and GSK-treated nuclei using the extracted biophysical parameters. All source data are provided as a Source Data file.

**Figure 5.**
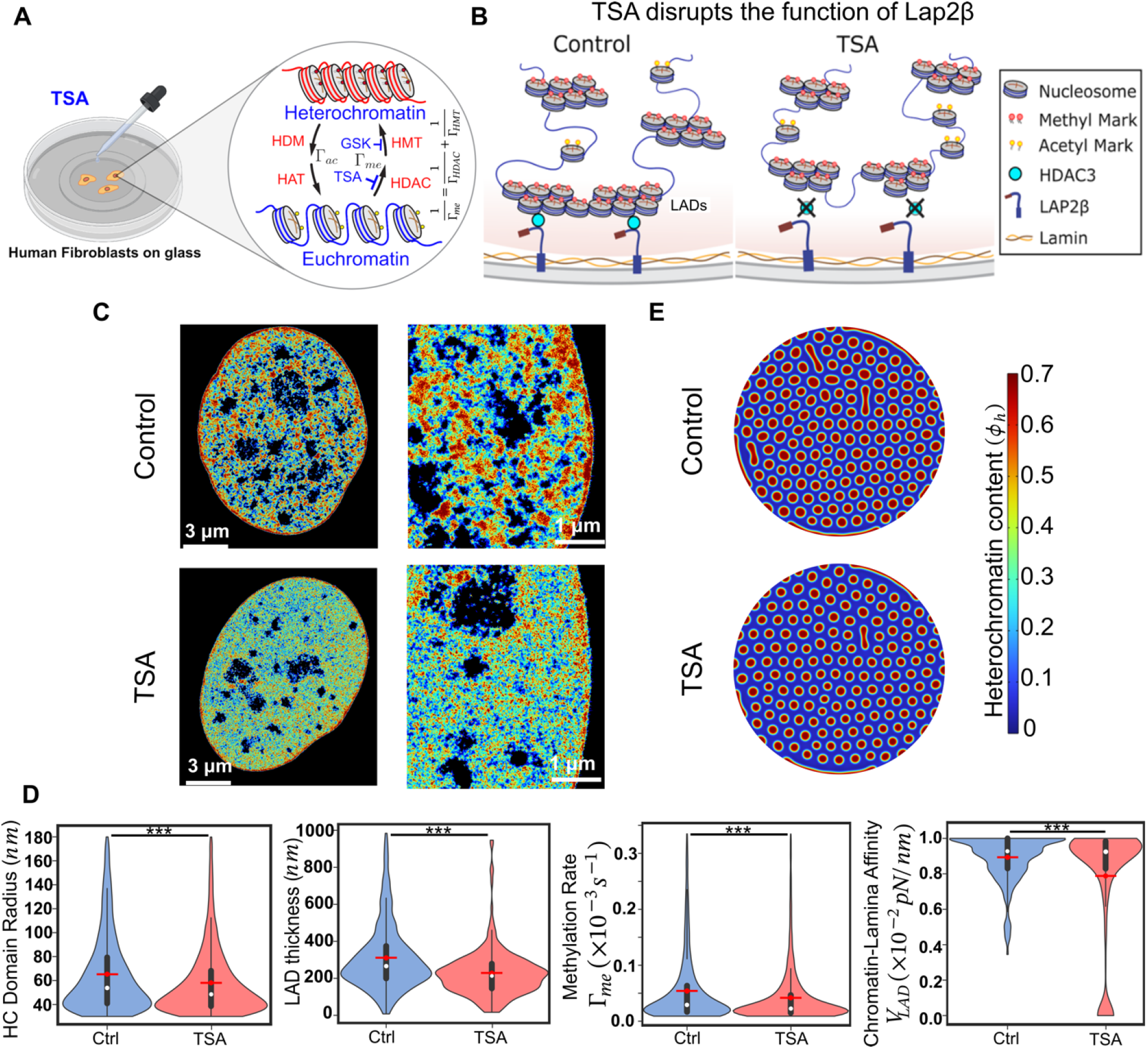
TSA treatment inhibits HDAC, resulting in reduction of methylation rate and chromatin-lamina affinity. (A) A schematic showing the role of TSA, which inhibits histone deacetylase (HDAC) and causes chromati decompaction. (B) A schematic showing the mechanisms of tethering chromatin to the nuclear periphery via HDAC3 and nuclear envelope proteins (e.g., LAP2β). TSA treatment disrupts the association of HADCs with LAP2β and causes the detachment of LADs thus decreasing chromatin-lamina affinity. (C) Super-resolution STORM images reveal the reorganization of chromatin upon TSA treatment. (Left panels) Voronoi density plots representing the extent of spatial compaction of chromatin in the control (top) and TSA-treated nuclei (bottom), along with a zoomed-in view (right panels). (D) Our theoretical framework reveals that upon TSA treatment, the interior heterochromatin domain radii, LAD thickness, methylation rate and chromatin-lamina affinity all decrease. (Control nuclei: n=6; TSA-treated nuclei: n=5, unpaired two tail test, *p<0.05, **p<0.01, ***p<0.001). All violin plots show a symmetric kernel density estimate (outline), the quartiles (black boxplot), the median (white dot), and the mean (red dot with line). (E) Numerical prediction using the extracted biophysical parameters as input showing smaller HC domains in the interior and periphery in TSA-treated nuclei. All source data are provided as a Source Data file.

**Figure 6.**
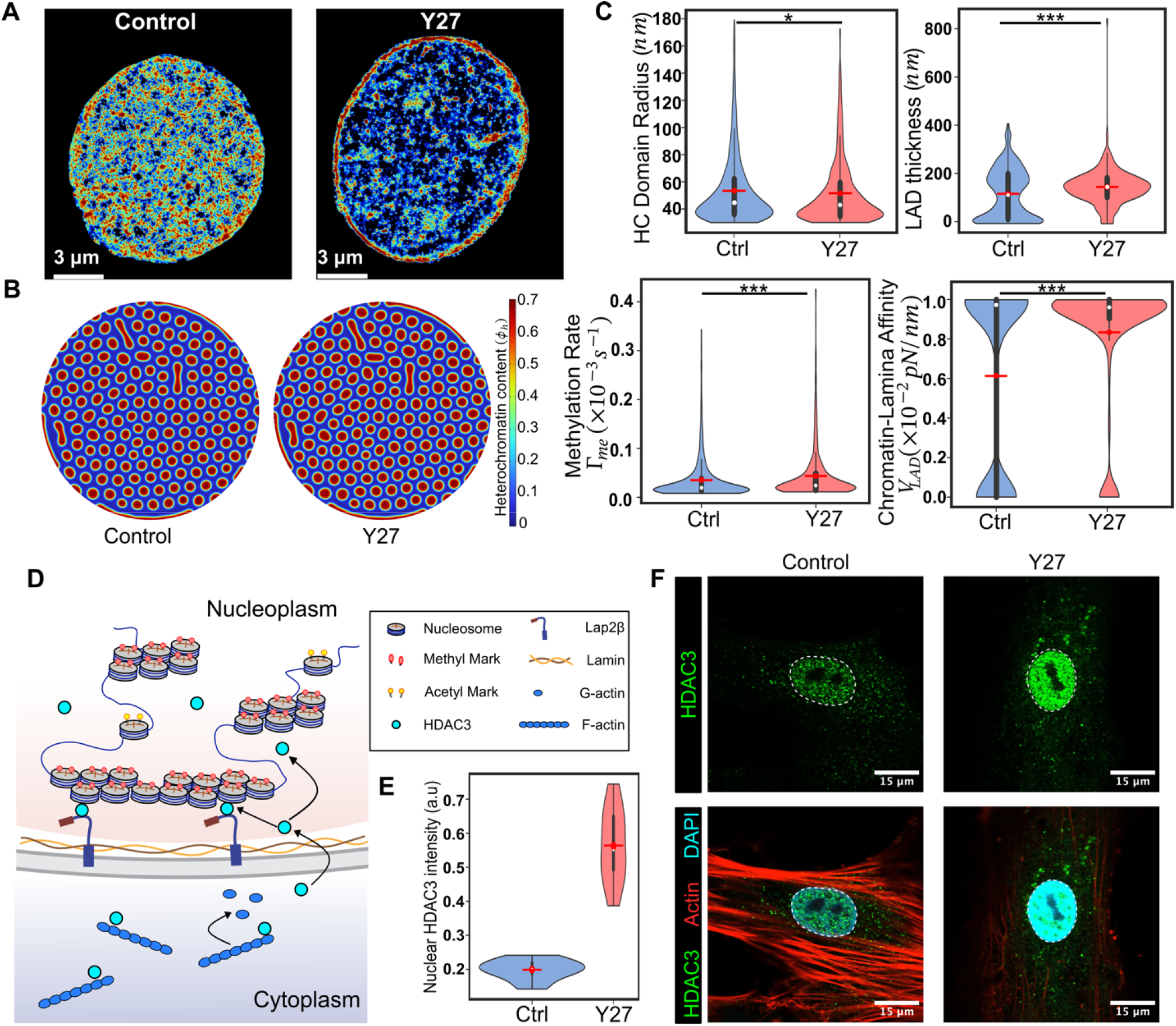
Inhibition of cellular contractility increases the nuclear localization of HDAC3, leading to a higher methylation rate and chromatin-lamina affinity, consequently resulting in larger LADs. A reduction in cellular contractility induces the nuclear translocation of HDAC3, leading to chromatin compaction. (A) Voronoi density plots representing the spatial reorganization of chromatin in the control and Y27-treated cells. (B) Numerical simulatio showing increased domain size and LAD thickness in Y27-treated nuclei. The methylation rate and chromatin-lamin affinity extracted from the STORM images are used as inputs in the model. (C) Quantitative analysis of STORM images reveals a decrease in the mean heterochromatin domain radius and an increase in LAD thickness, methylation rate, and chromatin-lamina affinity upon Y27 treatment. (Control nuclei: n=4; Y27-treated nuclei: n=5, unpaired two tail test, *p<0.05, **p<0.01, ***p<0.001). All violin plots show a symmetric kernel density estimate (outline), the quartiles (black boxplot), the median (white dot), and the mean (red dot with line). (D) A schematic showing the nuclear localization of HDAC3 in response to contractility abrogation. (E) HDAC3 fluorescence intensity quantification, showing increase in nuclear HDAC3 upon Y27 treatment. (F) Representative immunofluorescenc images for HDAC3 and actin in control and Y27-treated nuclei. All source data are provided as a Source Data file.

#### HMT inhibition affects only histone methylation and not chromatin-lamina affinity

EZH2 promotes the transfer of methyl groups to histone H3 at lysine 27 (H3K27) [35, 36]. We observe that its inhibition by GSK343 in hMSCs results in genome-wide chromatin decompaction compared to untreated control cells (Fig 4B)[9]. From the STORM images of H2B localizations (Fig 4B), we quantify the nuclear-wide distribution of heterochromatin domain sizes (as described in section 2.4) after control and GSK treatments (Fig 4C). Upon GSK treatment the mean interior heterochromatin domain radius *R*_*d*_ decreases by approximately 18% (Fig 4C). Next, by identifying the heterochromatin domains near the nuclear periphery (Section 2.4) we obtain the distribution of their average thickness *T*_*LAD*_ (as defined in SI section S5.2). GSK treatment results in decreased LAD thickness, with a mean LAD thickness approximately 20% lower than that of control-treated nuclei.

Having measured the sizes of the interior domains and LADs, we quantify the distribution of histone methylation rate, Γ_*me*_ and chromatin-lamina affinity, *V*_*LAD*_ in GSK and control treated nuclei (Fig 4C). Our theoretical framework predicts an approximately 36% lower mean methylation rate in GSK-treated cells than in untreated cells (Fig 4C). This is expected since GSK inhibits EZH2, a histone methyltransferase that contributes to methylation kinetics in our model (Fig 1B). However, we do not observe a significant change in chromatin-lamina affinity after GSK treatment (Fig 4C), indicating that inhibition of EZH2 does not influence the binding of chromatin to lamina according to our predictive framework.

Using the extracted changes in the parameters Γ_*me*_ and *V*_*LAD*_ we simulate the change in chromatin organization using Eq (1-3). The numerical predictions of chromatin reorganization (Fig 4D) in GSK-treated and control nuclei closely match the observed in vitro chromatin reorganization (Fig 4B), validating our parameter extraction framework. The numerically observed mean heterochromatin domain sizes in GSK-treated nuclei decreased by approximately 20% and 30% in the interior and at the periphery of the nucleus, respectively (Fig S12), comparable to the findings in cells.

#### HDAC inhibition reduces overall histone methylation and chromatin-lamina affinity

Similar to HMTs, HDACs play an important role in chromatin organization and gene expression via removal of acetyl groups from histones promoting chromatin condensation (Fig 5A) [37]. However, unlike EZH2, HDAC3 also contributes to the tethering of chromatin to the nuclear periphery via interactions with LAP2β, a nuclear envelope protein (Fig 5B)[21]. Using super-resolution imaging of histones and DNA, we previously showed that inhibition of HDAC via TSA treatment in human fibroblast (hFb) nuclei results in reduced chromatin compaction compared to that in untreated control nuclei (Fig 5C) [20]. Quantitative analysis of domain sizes after TSA treatment confirmed these results, revealing reductions of approximately 11% and 26% in the interior domain sizes and LAD thicknesses, respectively (Fig 5D). We next extract the changes in histone methylation rate and chromatin-lamina affinity (Fig 5D) using the theoretical framework developed in section 3.2. Our analysis predicts that the histone methylation rate is reduced by approximately 22% in TSA-treated nuclei compared to that in untreated nuclei (Fig 5D). The TSA-treatment mediated inhibition of histone deacetylation, which is a sub step of the histone methylation reaction in our model (Fig 1B) explains the predicted reduction in .

Additionally, the chromatin-lamina affinity in TSA-treated nuclei reduced by approximately 11% compared to that in untreated nuclei (Fig 5D). The decrease in chromatin-lamina affinity explains why the reduction in sizes of peripheral LADs is more pronounced than that in the interior domains. HDAC3 at the nuclear periphery interacts with chromatin anchoring protein LAP2 and mediates gene repression [21]. Abrogation of HDAC enzymatic activity via TSA treatment has been shown to release specific endogenous gene loci from the inner nuclear membrane mediating their movement toward the nucleus interior [38-40]. Thus, the predicted decline in chromatin-lamina affinity, observed using our theoretical framework, can be ascribed to the involvement of HDACs, such as HDAC3 which is involved in mediating the interactions between heterochromatin and the nuclear lamina[21]. The numerical simulations (Fig 5E), using the predicted changes in parameters, replicate the chromatin reorganization observed in cells upon TSA-treatment (Fig 5C) with reductions of approximately 11% and 22% in the interior domain and LAD sizes, respectively (Fig S12).

Overall, by quantitative analysis of STORM images, our theoretical framework accurately predicts the changes in biophysical parameters governing chromatin organization after GSK and TSA treatments. Notably, we predict that both TSA and GSK treatments reduce histone methylation rates, as expected in our model where deacetylation and methyltransferase activity contribute to the overall histone methylation rate (Fig 1B). However, TSA treatment, but not GSK treatment, reduces the strength of chromatin-lamina interactions. Indeed, EZH2, which is inhibited by GSK, is not a known contributor to the binding of chromatin to the lamina. Thus, solely based on STORM images, the model is able to account for the distinct actions of biochemical mechanisms altered during the GSK and TSA treatments without any prior knowledge of the specific pharmacological treatments applied. Next, we test our theory on upstream pharmacological perturbations that influence chromatin organization.

### 3.5 Abrogation of cell contractility alters chromatin-lamina interactions and LAD formation

The epigenetic regulators targeted in Section 3.4, EZH2 and HDAC3 exhibit a mechanosensitive nucleocytoplasmic partitioning [41]. For instance, mechanical stresses such as compressive or fluid shear stresses have been shown to alter the nuclear availability of EZH2 [42]. Furthermore, we have previously shown that cytoskeletal contractility alters the nucleocytoplasmic shuttling of HDAC3 such that under low contractility conditions, the nuclear localization of HDAC3 increases (Fig 6D) [8]. Thus, to study how upstream cellular perturbations such as cell contractility can alter chromatin organization, we analyzed STORM images of hMSCs treated with Y27632 (Y27; a specific inhibitor of rho-associated protein kinase II (ROCK II))[9].

As we previously demonstrated [9], STORM imaging of H2B localization in Y27-treated nuclei reveals global chromatin reorganization (Fig 6A). The quantification of domain sizes reveals that the mean radius of interior domains in Y27-treated nuclei was approximately 4.3% lower than that in control nuclei (Fig 6C). On the other hand, the mean LAD thickness increased by approximately 19.4% after Y27 treatment. The abrogation of contractility upon Y27 treatment enhances the nuclear localization of HDAC3 (Fig 6E and 6F), which is consistent with previous observations[8]. The increase in HDAC3 increases the rate of histone deacetylation, thereby amplifying the overall rate of methylation (as defined in Section 2.3, Fig 1B) and therefore the heterochromatin content. However, as discussed in Section 3.1, significant sequestering of heterochromatin along the periphery leaves less heterochromatin for the nucleus interior, leading to a reduction in the size of the interior domains. To confirm this interplay, we quantify the ratio of peripheral to total localization of heterochromatin in the nuclei from the STORM images. We find that Y27-treated nuclei exhibit significantly more peripheral heterochromatin (Fig S14) than control nuclei, suggesting a stronger sequestration of chromatin to the lamina. Thus, our model explains how a pronounced increase in LAD thickness after Y27 treatment drives a reduction in the size of interior heterochromatin domains, despite an increase in global chromatin condensation.

Using our theoretical framework, we next extract the distributions of histone methylation rate and chromatin-lamina affinity. We observe that the mean histone methylation rate in Y27-treated nuclei increases by approximately 12% compared to untreated nuclei (Fig 6C). Moreover, the chromatin-lamina affinity of Y27-treated nuclei was predicted to be approximately 26% higher compared to untreated nuclei (Fig 6C). The increase in nuclear HDAC3 due to the loss of contractility (Fig 6E) results in increased deacetylation of histones, which drives the predicted increase in the histone methylation rate in our model. Moreover, HDAC3 plays an additional role in tethering heterochromatin to the lamina via interactions with the nuclear membrane protein LAP2*β* [38, 43]. Thus, our framework predicts that the nuclear localization of HDAC3 in response to contractility abrogation amplifies not only the histone methylation rate, but also the chromatin-lamina affinity. Numerical simulations (Fig 6B) incorporating the spatially heterogeneous biophysical parameters extracted from the theoretical framework recapitulate the in vitro chromatin reorganization (Fig 6A). We observe an approximate 7% and 15% increase in domain sizes in the nuclear interior and at the periphery, respectively (Fig S12).

Altogether, our theoretical framework is able to predict the changes in histone methylation rate and chromatin-lamina affinity even when chromatin organization is perturbed via upstream modulations such as abrogation of contractility. As a further examination of the framework and to demonstrate its robustness beyond pharmacological treatments, we next perturb the cytoskeletal contractility via biophysical mechanisms such as altering the biomechanical environment of the cell.

### 3.6 Lower substrate stiffness leads to an increase in HDAC3 mediated chromatin-lamina affinity

The biomechanical environment, which is altered after disease, injury, or age-dependent tissue - degeneration, drives cytoskeletal remodeling in cells. For instance, in stiff environments, cell contractility increases due to stable substrate adhesions and stress fibers[44]. We have previously demonstrated the regulatory role of in-vitro substrate stiffness on chromatin compaction in the interior and periphery of hMSC nuclei [9]. The heatmaps of H2B localization densities, obtained via STORM imaging of hMSCs cultured on glass ( 70GPa), stiff (30 kPa), and soft (3 kPa) hydrogel substrates are shown in Fig 7A. Substrate-dependent chromatin remodeling affects both interior domains and LADs (Fig 7A, left panels). Quantitative analysis reveals an approximately 6.8% and 8% decrease in the mean interior domain radius in nuclei on stiff and glass substrates, respectively, compared to those on soft substrates (Fig 7B). At the nuclear periphery, compared to nuclei on soft substrates in nuclei on stiff and glass substrates the thickness of LADs reduced by approximately 14% and 21.5% respectively (Fig 7B). Thus, an increase in substrate stiffness results in reduced heterochromatin domain sizes in the interior and periphery of the nucleus.

**Figure 7.**
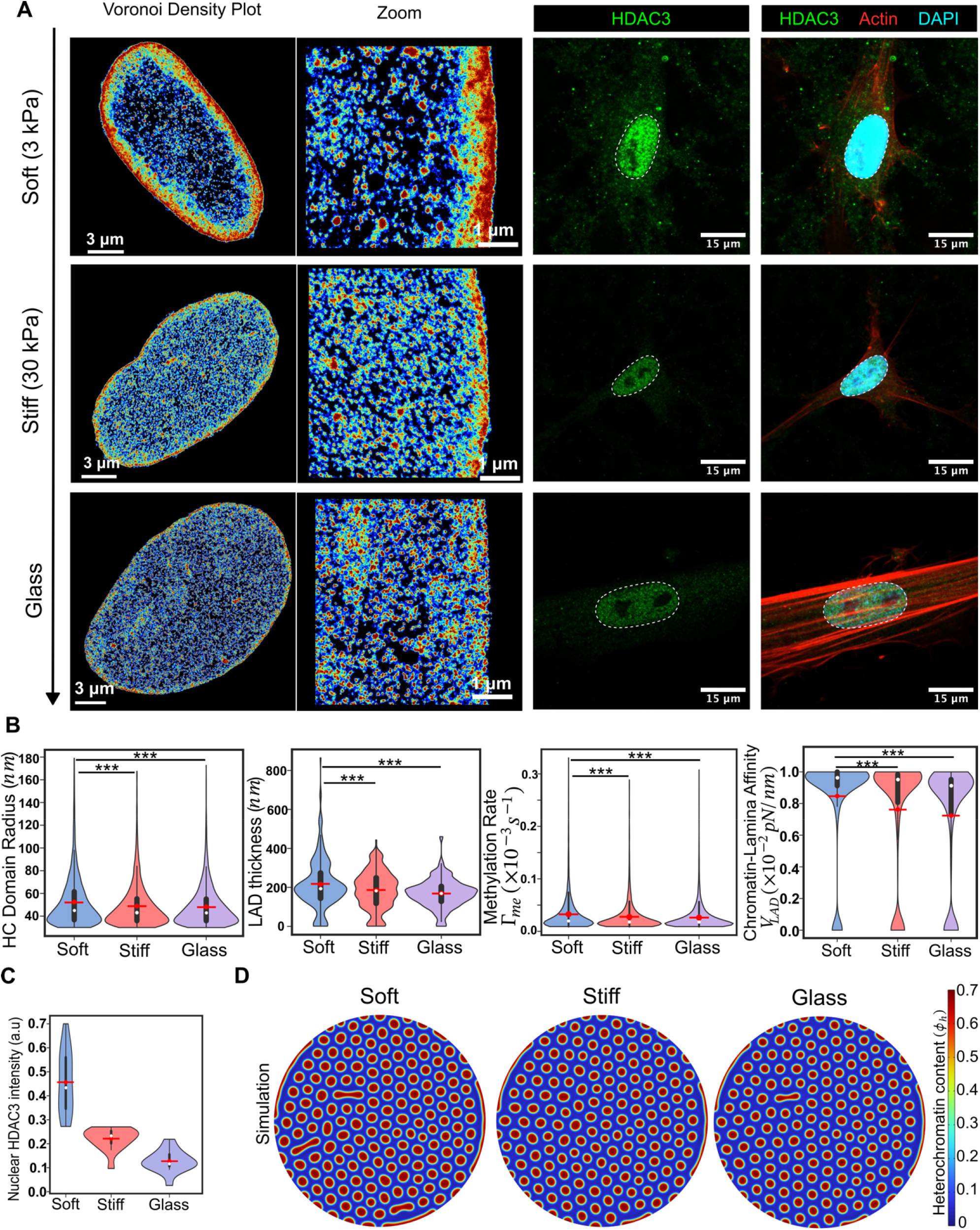
Higher substrate stiffness elevates cellular contractility resulting in smaller LADs. (A) (Left panels) Voronoi density plots of STORM images of hMSCs representing the spatial compaction of chromatin on soft (3kPa), stiff (30kPa) and glass (rigid) substrates, with an enlarged view. Cellular contractility is elevated on stiffer substrates, leading to the translocation of HDAC3 into the cytoplasm from the nucleus, which results in chromatin decompaction. Similarly, there is an increase in HDAC3 and EZH2 levels on softer substrates, which drives chromatin condensation and the attachment of peripheral heterochromatin from the lamina. (A) (Right panels) Representative immunofluorescence images for HDAC3 and actin on varying substrate stiffness (B) Quantitative analysis of the STORM images, integrated with theory, showing the distribution of heterochromatin domain radii, LAD thickness, methylation rates, and chromatin-lamina affinity on substrates of different stiffnesses. (Soft nuclei: n=5; stiff nuclei: n=5; glass nuclei: n=4, unpaired two tail test, *p<0.05, **p<0.01, ***p<0.001). All violin plots show a symmetric kernel density estimate (outline), the quartiles (black boxplot), the median (white dot), and the mean (red dot with line). (C) HDAC3 fluorescence intensity quantification, showing decrease in nuclear HDAC3 with increasing substrate stiffness. (D) Numerical simulation, using the biophysical parameters from the STORM images, showing a decreasing trend in the HC domain (in red) size with increasing substrate stiffness. All source data are provided as a Source Data file.

Next, using our theoretical framework, we extract the nucleus-wide distribution of histone methylation rate and chromatin-lamina affinity. Compared to nuclei cultured on soft substrates, we predict that those on stiff substrates exhibited a 17% decrease in the mean methylation rate. In the nuclei on glass the mean methylation rate was further reduced, showing a 19.9% decrease compared to that in the nuclei on soft substrates (Fig 7B). Furthermore, we observe a 10% and 14.5% lower chromatin lamina affinity on stiff and glass substrates, respectively, than on soft substrates (Fig 7B). The increase in stiffness from soft to stiff to glass substrates drives a progressive increase in cell contractility thereby reducing the intranuclear HDAC3 (Fig 7A, right panels). Hence, the loss of nuclear HDAC3 due to increase in stiffness (Fig 7C) results in reduced deacetylation of histones, thereby reducing the parameter Γ_*me*_ in our model. At the nuclear periphery, as discussed in the previous section, the loss of HDAC3 also results in a decrease in heterochromatin tethering to the nuclear lamina effectively reducing chromatin-lamina affinity.

Previously, it has been shown that via anchoring proteins such as emerin, Lamin A/C is implicated in heterochromatin tethering to the lamina [45, 46] and that stiff tissue environment drives an increase in Lamin A/C [47]. However, we observe a lower chromatin lamina affinity on stiffer substrates. Thus, our model suggests HDAC3 plays a relatively more prominent role in the formation of LADs as compared to Lamin A/C, at least in the cell-culture conditions described in Section 2.4. In agreement with this conclusion, previous work showed that knockdown of Lamin proteins does not lead to changes in LADs in mouse embryonic stem cells [48]. Finally, the distribution of the extracted methylation rate and chromatin-lamina affinity is used in our numerical simulations to predict the changes in chromatin reorganization (Fig 7D). The numerical simulations show an approximately 7% (10%) and 15% (25%) decrease in the interior and LAD sizes on stiff (glass) substrate respectively, compared to those on soft substrate (Fig S12), in agreement with the cell measurements (Fig 7B).

Thus, our theoretical framework accurately predicts changes in biophysical parameters associated with chromatin reorganization in cells cultured in vitro on substrates of varying stiffness. This validation leads us to enquire whether the in vivo chromatin reorganization due to tissue degenerative diseases such as tendinosis can be assessed with our theoretical framework to extract the modulation of biophysical parameters.

### 3.7 Tendinosis induces chromatin reorganization by altering microenvironmental stiffness

Injury and disease cause pathological degeneration of the fibrous extracellular matrix, altering the chemo-mechanical microenvironment of cells. Previous studies have shown that tissue degeneration in tendinosis reduces local tissue stiffness [49, 50]. We have demonstrated that human tenocyte (hTC) nuclei isolated from patients with tendinosis exhibit aberrant chromatin reorganization akin to that of healthy tenocytes cultured in a soft matrix [9]. Super-resolution STORM images of H2B localization densities in diseased hTC nuclei exhibit a global chromatin organization distinct from healthy nuclei (Fig 8a). We observe approximately 5.8% larger interior domains and 10.2% higher LAD thickness in diseased hTCs than in healthy cells (Fig 8B). Thus, tendinosis correlates with increased heterochromatin domain sizes both in the nucleus interior and periphery, resembling cells on soft substrates (Section 3.6).

**Figure 8.**
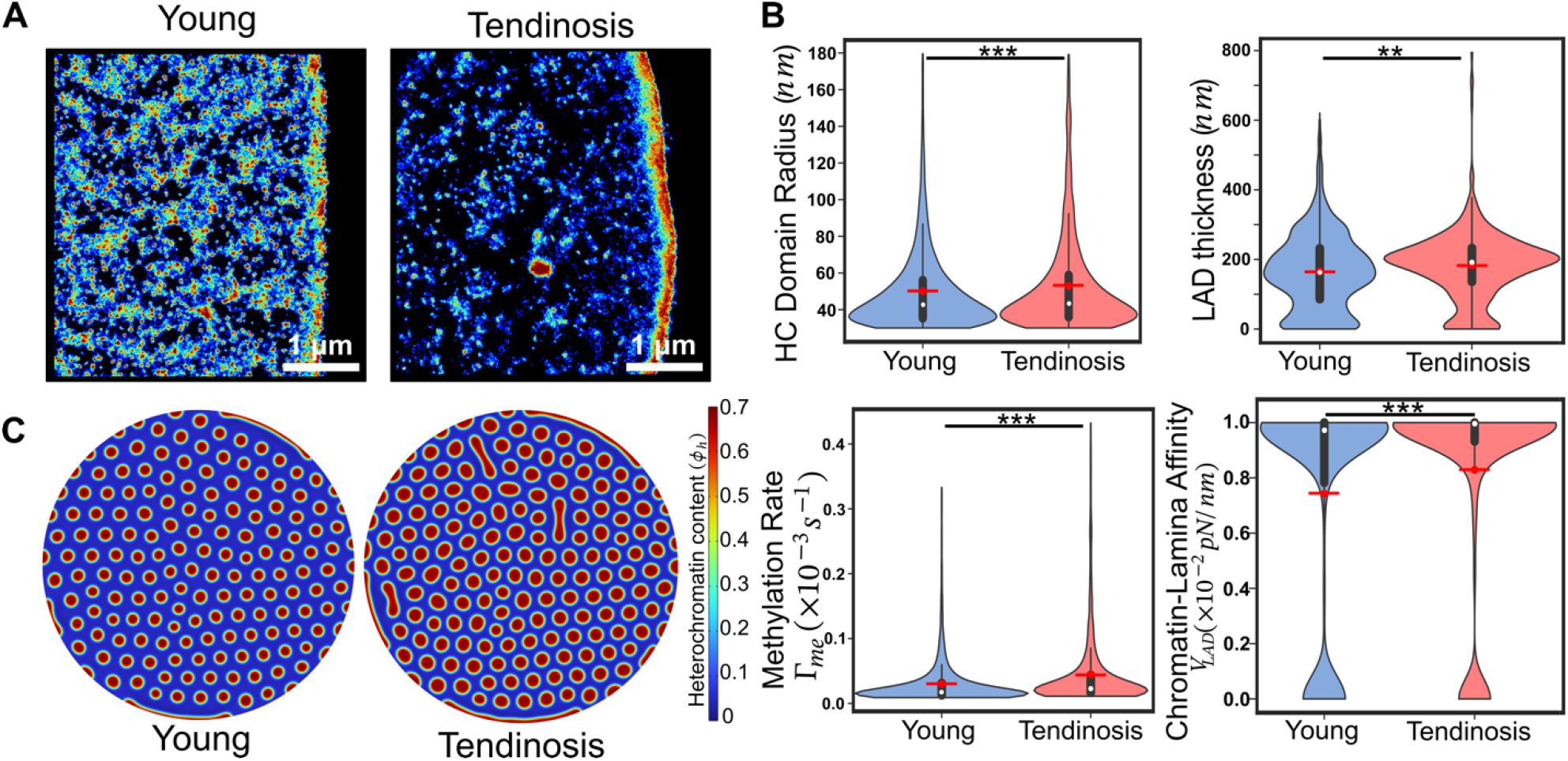
Tendinosis induces soft-like phenotype leading to larger LADs. (A) Enlarged images of the Voronoi density plot for young and diseased (tendinosis) human tenocytes (hTCs), showing changes in chromati condensation. (B) Quantitative analysis reveals an increase in the mean heterochromatin domain radius, LAD thickness, methylation rate and chromatin-lamina affinity. (Healthy nuclei: n=10; Diseased nuclei: n=9, unpaired two tail test, *p<0.05, **p<0.01, ***p<0.001). All violin plots show a symmetric kernel density estimate (outline), th quartiles (black boxplot), the median (white dot), and the mean (red dot with line). (C) Numerical simulation, employing biophysical parameters extracted from the integration of STORM images and theory as inputs, shows an increase in both the HC domain size and the LAD thickness. All source data are provided as a Source Data file.

We next quantify the distributions of histone methylation rate and chromatin lamina affinity in healthy and diseased tenocytes. Our theoretical framework reveals that in nuclei with tendinosis the mean histone methylation rate increases by approximately 30% while the chromatin-lamina affinity rises by 11.6% compared to that in healthy counterparts (Fig 8B). Note that although the methylation rate increased significantly, the increase in the size of interior heterochromatin domains was relatively low. This is due to the significant sequestering of heterochromatin to the nuclear periphery driven by the increase in chromatin-lamina affinity, as discussed in Section 3.1. This is further discussed in detail in the SI (Section S10). Thus, the changes in biophysical parameters after in-vivo tissue degeneration mirror those in cells cultured in vitro on soft substrates (Section 3.6). The induction of a soft phenotype in the tenocytes can be attributed to tendinosis driven collagen degeneration resulting in a reduction in tissue stiffness, as discussed in section 3.5. As in the section 3.6, although a reduction in microenvironmental stiffness is expected to result in loss of Lamin A/C, we instead observe an increase in peripheral LAD thickness and the extracted chromatin-lamina affinity. Thus, our model again suggests that HDAC3 and LAP2 play a relatively prominent role in driving LAD formation. Incorporating the extracted changes in the histone methylation rate and chromatin-lamina affinity in our numerical simulations, we observe that the mean interior domain radius increases by 16%, while the mean LAD thickness increases by 28% (Fig S12) comparable to what was observed in vitro (Fig 8B).

Altogether we find that our theoretical framework offers a methodology to accurately extract the changes in biophysical parameters in a versatile range of nuclear perturbations, not only in vitro but also after in vivo pathological degeneration, such as in tendinosis.

## 4. Discussion

In this study, we integrate theoretical and numerical modeling with super-resolution microscopy to introduce a multimodal framework for analyzing chromatin organization images and extracting the distribution of chromatin-lamina interaction strengths along the nuclear periphery. Such interactions are pivotal in regulating gene expression and spatial chromatin organization during cell development, differentiation, and disease [4, 51]. While the involvement of nuclear envelope proteins in mediating genome tethering to the lamina is known, understanding the variation in interaction strength along the periphery and across different cellular microenvironments remains unclear. Quantitatively evaluating these interactions necessitates analyzing super-resolution images of peripheral chromatin with a predictive biophysical model, which has been a challenge. We present a mesoscale mathematical model predicting the nucleus-wide chromatin organization in response to changes in the cell microenvironment, epigenetic drugs, and pharmacological treatments affecting cell contractility. Our model captures the emergence of densely compacted heterochromatin domains of characteristic sizes in both the nucleus interior and at the periphery. Leveraging these observations, our model predicts alterations in spatial chromatin organization in disease states where the extracellular matrix degradation leads to softening. Building upon our theoretical model, we develop a predictive framework that utilizes the morphology of peripheral heterochromatin domains observed via super-resolution microscopy as an indicator of chromatin-lamina interaction strength distribution. This data enables us to evaluate the roles of molecular mechanisms mediating these interactions in diverse cellular microenvironments.

At the nuclear periphery, we observe a spectrum of morphologies in heterochromatin domains, ranging from small, isolated clusters to elongated formations extending along the lamina. Our theoretical analysis identifies two key biophysical factors — local chromatin-lamina affinity and epigenetic reaction rates — as pivotal in determining the shapes and sizes of these peripheral domains. We demonstrate quantitatively that increased chromatin-lamina affinity leads to preferential spreading of heterochromatin domains along the nuclear lamina, forming elongated lamellar structures. Conversely, elevated histone methylation rates result in larger peripheral domains, albeit with minimal impact on their shape. Consequently, we predict that the average thickness of peripheral heterochromatin domains, indicative of both size and shape, is a composite outcome of chromatin-lamina interactions and histone methylation rates, as illustrated in a phase diagram predicted by our model (Fig S8). Within the nucleus interior, we find that the steady-state size of heterochromatin domains is uniquely regulated by epigenetic reaction kinetics. Increasing (decreasing) histone methylation rates lead to the formation of larger (smaller) domains (Fig 2E). Thus, given a distribution of epigenetic reaction rates and chromatin-lamina affinities, our model can predict the nucleus-wide size distribution of interior and peripheral domains. The size distribution of heterochromatin domains can be directly inferred from STORM images [3, 9, 20]. By inversely employing theoretically derived size-scaling relationships, we extract the nucleus-wide distribution of epigenetic reaction rates and chromatin-lamina affinity (Fig 3A). Our predictive framework, encompassing image quantification and subsequent parameter-extraction algorithms, is not limited to STORM and can be easily adapted to other super-resolution techniques enabling the visualization of heterochromatin domains.

A notable outcome arising from our predictive framework is the observation that the distribution of the chromatin-lamina interaction strength exhibits a bimodal pattern, characterized by two distinct peaks (Figure 9B). One peak signifies minimal interaction strengths, while the other denotes robust interactions akin to those observed between segments of chromatin. This spatial heterogeneity in chromatin-lamina affinity distribution likely stems from the considerable spatial variability in the composition of the lamina meshwork, alongside the presence of diverse nuclear membrane proteins such as ion channels, nuclear pores, and chromatin-tethers like LBR and LAP2β [52]. While certain nuclear membrane proteins like LBR, LAP2β, emerin, and HDAC3 actively tether heterochromatin, other functional membrane components like ion channels, pumps and nuclear pore complexes only interact very weakly with chromatin. This inherent variability in chromatin-lamina interactions is effectively captured in our model, resulting in the bimodal distribution of nucleus-wide chromatin-lamina affinity across all nuclei analyzed (Fig 3C, S13). In STORM images, the peak representing a high degree of chromatin-lamina interactions typically correlates with dense H2B localizations, manifesting as large peripheral heterochromatin domains extensively distributed along the lamina. Conversely, regions along the periphery corresponding to the second peak, characterized by minimal chromatin-lamina interactions, exhibit little to no presence of peripheral heterochromatin domains. Between these extremes, a spectrum of intermediate chromatin-lamina interactions is observed, with small and thin peripheral heterochromatin domains indicative of regions with relatively weaker but still perceptible affinity for heterochromatin. Thus, the extracted distribution of chromatin-lamina affinity provides insights into the distinct mechanisms involved in peripheral chromatin sequestration.

**Figure 9.**
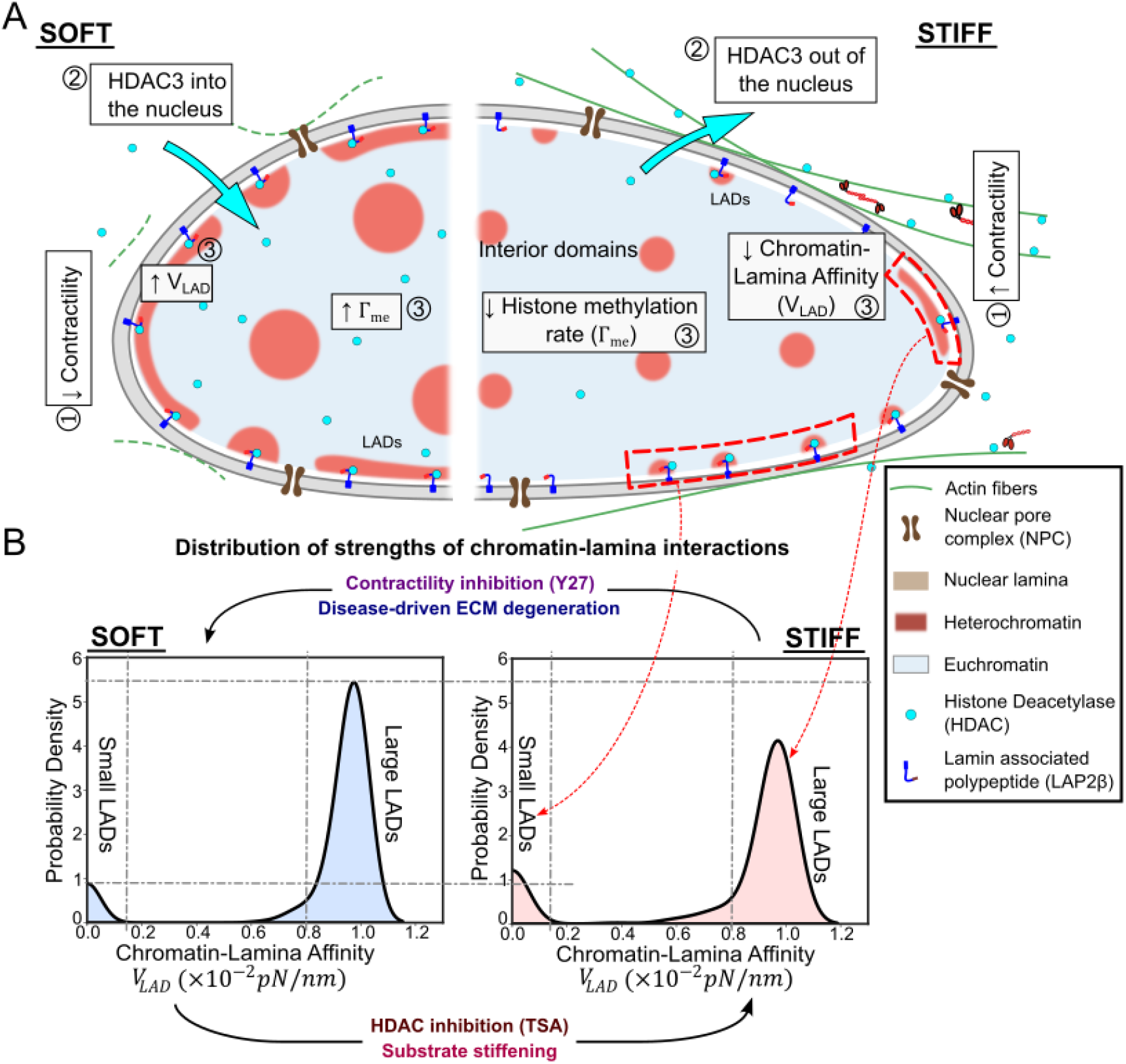
A summary of the proposed model for synergistic regulation of LAD organization by epigenetic reactions and the strength of chromatin-lamina interactions. (A) Schematic depicting the shuttling of HDAC3 between the cytoplasm and the nucleus depending on the cellular contractility. The right panel shows the organization of chromatin on stiff substrates, where higher contractility leads to reduced intranuclear HDAC3 levels and hence also lower chromatin-lamina affinity. The left panel shows chromatin organization in cells on soft substrates with higher methylation levels and lamina affinity. (B) The nucleus-wide distribution of the strength of chromatin-lamina affinity extracted from our framework shows a bimodal distribution with one peak at vanishing chromatin-lamina interactions with small discrete LADs (labelled small LADs) and another at strong chromatin-lamina interactions, comparable to the chromatin-chromatin interactions, with large continuous LADs (labelled large LADs). On a stiff substrate reduced nuclear HDAC3 contributes to decreasing the chromatin-lamina interaction strength. Note that the peak corresponding to large LADs decreases. (B) also summarizes the effects of pharmacological perturbations (TSA, Y27 treatments), change in substrate stiffness and in-vivo tendinopathic ECM degeneration on the distribution of the chromatin-lamina interaction strengths.

Our model predictions closely align with the heterogeneity observed in Lamin B1 enrichment in genomic regions across multiple cell types, as unveiled by LB1 ChIP-seq [31]. The peak of high chromatin-lamina affinity predicted by our model likely corresponds to genomic regions with high Lamin B1 occupancy (classified as Type 1 LADs). Conversely, the predicted range of weaker but nonzero chromatin-lamina affinities may resemble genomic regions exhibiting relatively lower Lamin B1 occupancy (Type 2 LADs) in LB1 ChIP-seq [31].

The cell microenvironment can intricately regulate the nucleus-wide distribution of epigenetic reaction rates and chromatin-lamina affinity in a mechanosensitive manner (Fig 9A). Environmental factors like stiffness can modulate the nuclear localization of HDAC3, influencing histone deacetylation rates [53]. Additionally, mechanical forces such as compression and shear loads inhibit EZH2 activity, a histone methyltransferase [42, 54, 55]. To mimic these mechanosensitive variations, we analyze chromatin organization in cells pharmacologically treated with HDAC and EZH2 inhibitors using TSA and GSK, respectively. The effective methylation rate is determined by both histone deacetylation and methyltransferase activities, as illustrated in Fig. 1B, expressed as 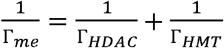. Consequently, both TSA and GSK treatments reduce the overall methylation reaction rate, impacting the size-scale of both the interior and peripheral heterochromatin domains. Our analysis reveals that TSA treatment not only perturbs epigenetics but also reduces chromatin-lamina affinity, particularly affecting peripheral domains. Conversely, GSK treatment does not significantly alter chromatin-lamina interactions. Thus, our quantitative assessment of nuclei after in vitro treatments implicates HDAC3, and not EZH2, in chromatin sequestering at the lamina. Consistent with our model predictions, HDAC inhibition via TSA treatment and RNA-induced inhibition can detach gene loci from the nuclear lamina, relocating them to the nucleus interior [38-40, 56]. This finding confirms the role of HDAC3 in tethering chromatin to the lamina, particularly evident near the nuclear periphery where HDAC3 interacts with LAP2β and emerin, suppressing gene expression [21, 43]. Overall, our model accurately predicts methylation rate distribution and chromatin affinity to the nuclear periphery, effectively distinguishing between pharmacological treatments that impact one or both of these processes.

To explore how methylation rates and the affinity of chromatin to the nuclear periphery depend on mechanochemical signaling, we analyzed chromatin organization on substrates of varying stiffness. We found that compared to nuclei on a stiff hydrogel substrate, nuclei on a soft hydrogel substrate exhibited an increase in the size-scales of both the interior and peripheral heterochromatin domains. Utilizing our model to extract biophysical parameters, we predicted that on a soft substrate, both the histone methylation rate and chromatin-lamina affinity are elevated compared to a stiff substrate. To elucidate the amplification of both parameters, we observed an augmented nuclear accumulation of HDAC3, accompanied by F-actin depolymerization on soft substrates compared to stiff and glass substrates. These findings align with our prior reports that on stiff substrates, increased cytoskeletal contractility results in the shuttling of HDAC3 from the nucleus into the cytoplasm in mouse fibroblasts [8, 53]. To validate that the loss of contractility on soft substrates indeed contributes to the altered chromatin organization, we also pharmacologically disrupted cell contractility using Y27 treatment, after which we observed that both the histone methylation rate and chromatin-lamina affinity increase—similar to nuclei on soft substrates. However, after Y27 treatment the increase in chromatin-lamina affinity is more pronounced than on a soft substrate resulting in excessive sequestering of chromatin to the lamina reducing the size of the interior domains (as discussed in Section S10). This may be mediated by altered chromatin-binding after Y27 treatment due to mechanisms in addition to HDAC3 [6, 57]. While this requires further investigation, it can be easily incorporated into the model by suitably adapting the mechanosensitive chromatin-lamina interactions as a function of such binding mechanisms.

Thus, our observations strongly suggest that via the shuttling of epigenetic factors like HDAC3, cytoskeletal contractility chemo-mechanically transduces environmental stiffness stimuli to the nucleus and drives chromatin reorganization. Changes in environmental stiffness are observed in vivo due to pathological degeneration of the extracellular matrix, such as in tendinosis resulting from overuse and repetitive stress [49, 58]. Investigating the aberrant chromatin reorganization in tenocytes due to tendinosis, our theoretical framework predicts that both the histone methylation rate and chromatin-lamina interactions are increased, mirroring the biophysical parameters extracted for in vitro nuclei exhibiting a soft phenotype.

The predictive framework developed here for extracting the nucleus-wide distribution of biophysical parameters through quantitative inference from image analysis is unbiased and does not rely on prior knowledge of the stimulus imparted on the nucleus. Thus, in situations where the underlying mechanism of chromatin remodeling is unknown, our model can elucidate the integrative roles of epigenetic reactions and chromatin-lamina interactions in regulating heterochromatin domain formation in both the interior and the periphery of the nucleus. Our numerical simulations can also capture the time-dependent dynamics of epigenetic regulation and chromatin-lamina interactions, enabling prediction of parameter changes governing dynamic chromatin remodeling in response to temporally varying chemo-mechanical stimuli. Moreover, our model can be refined in combination with polymer modeling to depict genome organization at the scale of individual chromatin compartments, focusing on the expression and dynamics of specific gene loci. When coupled with imaging techniques such as FISH and Oligopaint probes and super-resolution imaging [59, 60], our framework can identify genes affected by epigenetic regulation or chromatin-lamina interactions. These diagnostic capabilities hold promising applications in identifying broad therapeutic targets to modulate gene expression near the periphery or within the interior of the nucleus. Such predictions could reverse the effects of environmental stimuli or favorably modulate chromatin organization. Thus, our multi-modality-based parameter extraction framework establishes a fundamental framework for better understanding the mechanisms through which environmental stimuli drive chemo-mechanical nucleo-cytoskeletal signaling, thereby altering mesoscale chromatin organization in the nucleus both internally and at the periphery of the nucleus.

## Data availability

The data supporting the findings of this study are available from the corresponding author upon request. The data generated in this study are provided in the Source Data file.

## Code availability

The code used for measurement of sizes of heterochromatin domain obtained from STORM imaging is freely available through github (https://github.com/ShenoyLab/STORM_Analysis_Parameter_Extraction).

## Acknowledgements

This work was supported by NIH Award U54CA261694 (V.B.S.); NCI Awards R01CA232256 (V.B.S.); NSF CEMB Grant CMMI-154857 (V.B.S.); NSF Grants MRSEC/DMR-1720530 and DMS-1953572 (V.B.S.); NIBIB Awards R01EB017753 and R01EB030876 (V.B.S.). We gratefully acknowledge the valuable comments and suggestions from Dr. Rajan Jain.

## REFERENCES

1. Du, H., et al., Tuning immunity through tissue mechanotransduction. Nature Reviews Immunology, 2023. 23(3): p. 174–188.

2. Wang, N., J.D. Tytell, and D.E. Ingber, Mechanotransduction at a distance: mechanically coupling the extracellular matrix with the nucleus. Nature reviews Molecular cell biology, 2009. 10(1): p. 75–82.

3. Ricci, M.A., et al., Chromatin fibers are formed by heterogeneous groups of nucleosomes in vivo. Cell, 2015. 160(6): p. 1145–1158.

4. Alagna, N.S., et al., Choreography of lamina-associated domains: structure meets dynamics. FEBS Lett, 2023. 597(22): p. 2806–2822.

5. Buchwalter, A., J.M. Kaneshiro, and M.W. Hetzer, Coaching from the sidelines: the nuclear periphery in genome regulation. Nat Rev Genet, 2019. 20(1): p. 39–50.

6. Manzo, S.G., L. Dauban, and B. van Steensel, Lamina-associated domains: Tethers and looseners. Curr Opin Cell Biol, 2022. 74: p. 80–87.

7. Discher, D.E., P. Janmey, and Y.-l. Wang, Tissue cells feel and respond to the stiffness of their substrate. Science, 2005. 310(5751): p. 1139–1143.

8. Alisafaei, F., et al., Regulation of nuclear architecture, mechanics, and nucleocytoplasmic shuttling of epigenetic factors by cell geometric constraints. Proceedings of the National Academy of Sciences, 2019. 116(27): p. 13200–13209.

9. Heo, S.-J., et al., Aberrant chromatin reorganization in cells from diseased fibrous connective tissue in response to altered chemomechanical cues. Nature biomedical engineering, 2023. 7(2): p. 177–191.

10. Briand, N. and P. Collas, Lamina-associated domains: peripheral matters and internal affairs. Genome Biol, 2020. 21(1): p. 85.

11. Bowman, G.D. and M.G. Poirier, Post-translational modifications of histones that influence nucleosome dynamics. Chemical reviews, 2014. 115(6): p. 2274–2295.

12. Canzio, D., et al., Chromodomain-mediated oligomerization of HP1 suggests a nucleosome-bridging mechanism for heterochromatin assembly. Molecular cell, 2011. 41(1): p. 67–81.

13. Sanulli, S., et al., HP1 reshapes nucleosome core to promote phase separation of heterochromatin. Nature, 2019. 575(7782): p. 390–394.

14. Grosberg, A.Y. and J.-F. Joanny, Nonequilibrium statistical mechanics of mixtures of particles in contact with different thermostats. Physical Review E, 2015. 92(3): p. 032118.

15. Moller, J., J. Lequieu, and J.J. de Pablo, The free energy landscape of internucleosome interactions and its relation to chromatin fiber structure. ACS central science, 2019. 5(2): p. 341–348.

16. Kant, A., et al., Active transcription and epigenetic reactions synergistically regulate meso-scale genomic organization. Nature Communications, 2024. 15(1): p. 4338.

17. Kiseleva, A.A. and A. Poleshko, The secret life of chromatin tethers. FEBS Lett, 2023. 597(22): p. 2782–2790.

18. Harris, C.A., et al., Structure and mapping of the human thymopoietin (TMPO) gene and relationship of human TMPO β to rat lamin-associated polypeptide 2. Genomics, 1995. 28(2): p. 198–205.

19. Jain, N., et al., Cell geometric constraints induce modular gene-expression patterns via redistribution of HDAC3 regulated by actomyosin contractility. Proc Natl Acad Sci U S A, 2013. 110(28): p. 11349–54.

20. Otterstrom, J., et al., Super-resolution microscopy reveals how histone tail acetylation affects DNA compaction within nucleosomes in vivo. Nucleic acids research, 2019. 47(16): p. 8470–8484.

21. Somech, R., et al., The nuclear-envelope protein and transcriptional repressor LAP2beta interacts with HDAC3 at the nuclear periphery, and induces histone H4 deacetylation. J Cell Sci, 2005. 118(Pt 17): p. 4017–25.

22. Gardner, O.F., M. Alini, and M.J. Stoddart, Mesenchymal stem cells derived from human bone marrow. Cartilage tissue engineering: methods and protocols, 2015: p. 41–52.

23. McBeath, R., et al., Tendinosis develops from age-and oxygen tension-dependent modulation of Rac1 activity. Aging Cell, 2019. 18(3): p. e12934.

24. Zanacchi, F.C., et al., A DNA origami platform for quantifying protein copy number in super-resolution. Nature methods, 2017. 14(8): p. 789–792.

25. Levet, F., et al., SR-Tesseler: a method to segment and quantify localization-based super-resolution microscopy data. Nature methods, 2015. 12(11): p. 1065–1071.

26. Harr, J.C., A. Gonzalez-Sandoval, and S.M. Gasser, Histones and histone modifications in perinuclear chromatin anchoring: from yeast to man. EMBO reports, 2016. 17(2): p. 139–155.

27. Belmont, A.S., Y. Zhai, and A. Thilenius, Lamin B distribution and association with peripheral chromatin revealed by optical sectioning and electron microscopy tomography. J Cell Biol, 1993. 123(6 Pt 2): p. 1671–85.

28. Broide, R.S., et al., Distribution of histone deacetylases 1–11 in the rat brain. Journal of Molecular Neuroscience, 2007. 31: p. 47–58.

29. Gozalo, A., et al., Core Components of the Nuclear Pore Bind Distinct States of Chromatin and Contribute to Polycomb Repression. Mol Cell, 2020. 77(1): p. 67–81 e7.

30. Pascual-Garcia, P. and M. Capelson, The nuclear pore complex and the genome: organizing and regulatory principles. Curr Opin Genet Dev, 2021. 67: p. 142–150.

31. Shah, P.P., et al., An atlas of lamina-associated chromatin across twelve human cell types reveals an intermediate chromatin subtype. Genome Biol, 2023. 24(1): p. 16.

32. Heo, S.-J., et al., Biophysical regulation of chromatin architecture instills a mechanical memory in mesenchymal stem cells. Scientific reports, 2015. 5(1): p. 16895.

33. Sato, T., et al., Transcriptional selectivity of epigenetic therapy in cancer. Cancer research, 2017. 77(2): p. 470–481.

34. Tóth, K.F., et al., Trichostatin A-induced histone acetylation causes decondensation of interphase chromatin. Journal of cell science, 2004. 117(18): p. 4277–4287.

35. Dudakovic, A., et al., Epigenetic control of skeletal development by the histone methyltransferase Ezh2. Journal of Biological Chemistry, 2015. 290(46): p. 27604–27617.

36. Gan, L., et al., Epigenetic regulation of cancer progression by EZH2: from biological insights to therapeutic potential. Biomarker research, 2018. 6: p. 1–10.

37. Gallinari, P., et al., HDACs, histone deacetylation and gene transcription: from molecular biology to cancer therapeutics. Cell research, 2007. 17(3): p. 195–211.

38. Zullo, J.M., et al., DNA sequence-dependent compartmentalization and silencing of chromatin at the nuclear lamina. Cell, 2012. 149(7): p. 1474–87.

39. Pickersgill, H., et al., Characterization of the Drosophila melanogaster genome at the nuclear lamina. Nature genetics, 2006. 38(9): p. 1005–1014.

40. Zink, D., et al., Transcription-dependent spatial arrangements of CFTR and adjacent genes in human cell nuclei. The Journal of cell biology, 2004. 166(6): p. 815–825.

41. Song, Y., et al., Cell engineering: biophysical regulation of the nucleus. Biomaterials, 2020. 234: p. 119743.

42. Maleszewska, M., et al., The decrease in histone methyltransferase EZH2 in response to fluid shear stress alters endothelial gene expression and promotes quiescence. Angiogenesis, 2016. 19: p. 9–24.

43. Poleshko, A., et al., Genome-Nuclear Lamina Interactions Regulate Cardiac Stem Cell Lineage Restriction. Cell, 2017. 171(3): p. 573–587 e14.

44. Yeung, T., et al., Effects of substrate stiffness on cell morphology, cytoskeletal structure, and adhesion. Cell motility and the cytoskeleton, 2005. 60(1): p. 24–34.

45. Lee, K.K., et al., Distinct functional domains in emerin bind lamin A and DNA-bridging protein BAF. Journal of cell science, 2001. 114(24): p. 4567–4573.

46. Reddy, K., et al., Transcriptional repression mediated by repositioning of genes to the nuclear lamina. Nature, 2008. 452(7184): p. 243–247.

47. Swift, J., et al., Nuclear lamin-A scales with tissue stiffness and enhances matrix-directed differentiation. Science, 2013. 341(6149): p. 1240104.

48. Amendola, M. and B. van Steensel, Nuclear lamins are not required for lamina-associated domain organization in mouse embryonic stem cells. EMBO reports, 2015. 16(5): p. 610–617.

49. Arya, S. and K. Kulig, Tendinopathy alters mechanical and material properties of the Achilles tendon. Journal of applied physiology, 2010. 108(3): p. 670–675.

50. Finnamore, E., et al., Transverse tendon stiffness is reduced in people with Achilles tendinopathy: A cross-sectional study. PLoS One, 2019. 14(2): p. e0211863.

51. Bellanger, A., et al., Restructuring of lamina-associated domains in senescence and cancer. Cells, 2022. 11(11): p. 1846.

52. Turgay, Y., et al., The molecular architecture of lamins in somatic cells. Nature, 2017. 543(7644): p. 261–264.

53. Damodaran, K., et al., Compressive force induces reversible chromatin condensation and cell geometry–dependent transcriptional response. Molecular biology of the cell, 2018. 29(25): p. 3039–3051.

54. Li, Q., et al., EZH2 reduction is an essential mechanoresponse for the maintenance of super-enhancer polarization against compressive stress in human periodontal ligament stem cells. Cell death & disease, 2020. 11(9): p. 757.

55. Munne, P.M., et al., Compressive stress-mediated p38 activation required for ERα+ phenotype in breast cancer. Nature communications, 2021. 12(1): p. 6967.

56. Milon, B.C., et al., Role of histone deacetylases in gene regulation at nuclear lamina. PloS one, 2012. 7(11): p. e49692.

57. Czapiewski, R., M.I. Robson, and E.C. Schirmer, Anchoring a Leviathan: How the Nuclear Membrane Tethers the Genome. Front Genet, 2016. 7: p. 82.

58. Cho, Y., et al., CTRP3 exacerbates tendinopathy by dysregulating tendon stem cell differentiation and altering extracellular matrix composition. Science Advances, 2021. 7(47): p. eabg6069.

59. Beliveau, B.J., et al., Versatile design and synthesis platform for visualizing genomes with Oligopaint FISH probes. Proceedings of the National Academy of Sciences, 2012. 109(52): p. 21301–21306.

60. Park, D.S., et al., High-throughput Oligopaint screen identifies druggable 3D genome regulators. Nature, 2023. 620(7972): p. 209–217.

