## Supplementary Material for "Revealing the Biophysics of Lamina-Associated Domain Formation by Integrating Theoretical Modeling and High-Resolution Imaging"

For

### S1 Extended Methods

#### *S1.1 Mathematical description of chromatin organization in the nucleus*

To investigate the organization of chromatin in the nucleus, we develop a mathematical model for the phase separation of heterochromatin and euchromatin considering the chromatin-chromatin interactions, chromatin-lamina interactions, and epigenetic regulation of chromatin via histone acetylation or methylation. The nucleus is conceptualized as comprising three components: the nucleoplasm, heterochromatin, and euchromatin. At a given spatial coordinate $\boldsymbol{x}$ and time t the nuclear constitution is characterized by local volume fractions: $\phi_h(\boldsymbol{x},t)$ for heterochromatin, $\phi_e(\boldsymbol{x},t)$ for euchromatin and, $\phi_n(\boldsymbol{x},t)$ for nucleoplasm, with a stipulation that $\phi_e + \phi_h + \phi_n = 1$. This constraint ensures conservation of mass within the system. The physical state at any given location is thus discernible through two independent parameters: the volume fraction of nucleoplasm $\phi_n(\boldsymbol{x},t)$, and the net difference in volume fractions between heterochromatin and euchromatin, denoted as $\phi_d(\boldsymbol{x},t) = \phi_h(\boldsymbol{x},t) - \phi_e(\boldsymbol{x},t)$. The variable $\phi_d$ serves as an order parameter, signaling the dominance of either phase: negative values indicate a prevalence of euchromatin, while positive values denote a higher concentration of heterochromatin.

#### *S1.2 Free energy landscape of the nucleus*

In terms of the independent variables $\phi_d(\boldsymbol{x},t)$ and $\phi_n(\boldsymbol{x},t)$, the free energy density at any point $\boldsymbol{x}$ in the nucleus can be expressed as $f(\phi_n, \phi_d, \nabla\phi_n, \nabla\phi_d)$. This formulation incorporates the energetic considerations associated with the interfaces between phases through the gradients of the volume fractions. Specific form of the free energy can be invoked by considering the various energetic contributions in the nucleus such as,

$$f = \underbrace{\frac{c}{2}\left[\phi_e^2 + \phi_h^2(\phi_{h0} - \phi_h)^2\right]}_{\text{chromatin-chromatin interactions}} + \underbrace{\frac{\kappa}{2}\left[|\nabla\phi_n|^2 + |\nabla\phi_d|^2\right]}_{\text{Interfacial energy}} - \underbrace{\frac{V(\phi_h)}{d_0}\phi_h e^{-\frac{d}{d_0}}}_{\text{chromatin-lamina interactions}} \qquad \text{(S1)}$$

The first term in Eq S1 emerges from the interplay between the entropy and the enthalpy of mixing for the heterochromatin and euchromatin phases, conferring a double-well configuration to the free energy landscape, as shown in the contour plot in Fig 1C. The two wells, shown as blue and red dots, are the energy minima corresponding to the two stable phases of chromatin - a water-rich, loosely packed euchromatin phase ($\phi_h = 0$) or a compacted water-poor heterochromatin phase ($\phi_h = \phi_{h0}, \phi_n \sim 0$). Here $\phi_{h0}$ is maximum heterochromatin amount and denotes the extent of compaction in the heterochromatin phase. An initial chromatin state, indicated by a white dot, will naturally phase separate into domains of heterochromatin and euchromatin (white arrows).

The second term denotes the energy penalty associated with forming phase boundaries between the euchromatin and heterochromatin as they separate. The term $\kappa$ is the increase in the energy due to formation of a unit width of the interface. As $\kappa$ increases, there is a greater penalty on formation of sharp interfaces, resulting in more smooth interfaces which are wider. Thus $\kappa$ directly control the width and the energy of the phase boundaries. Note that the term $|\nabla\phi_{n,d}|$ is the magnitude of the slope of the interface.

The last term captures the interactions between the chromatin and the lamina via chromatin anchoring proteins (HDAC3, LAP2$\beta$, emerin)[1, 2] with function $V(\phi_h)$ (refer section S5.1 for details) denoting the strength of these anchoring interactions per unit area. These interactions are strongest at the nuclear periphery (distance from lamina $d = 0$) and vanish exponentially over a length scale $d_0$, comparable to size of anchoring proteins.

The total free energy of the nucleus can be written as,

$$\Pi[\phi_n, \phi_d] = \int_{V_n} f(\phi_n, \phi_d, \nabla\phi_n, \nabla\phi_d) dV - \int_{\partial V_p} \bar{\mu}_n I^n dA \tag{S2}$$

Here, $V_n$ is the volume of nucleus and $\partial V_p$ denotes the surface of the nucleus where water exchange between nucleus and cytoplasm takes place. $\bar{\mu}_n$ is external chemical potential and $I^n$ is the volume of water entering per unit surface area into the nucleus.

The energy landscape described by Eq 1 guides the chromatin's temporal dynamics from an initial configuration (white dot in Fig 1C) into the two energy minima, corresponding to the two chromatin phases (red and blue dot). The spatiotemporal evolution of nucleoplasm and chromatin occurs such that the free energy of chromatin organization is driven towards the energy minima. The driving forces for this evolution are called the chemical potential ($\mu_n$ and $\mu_d$) of nucleoplasm and epigenetic marks respectively and are quantitatively dependent on the gradients in the energy landscape. Using variational principles, these chemical potentials are written as,

$$\mu_n(\boldsymbol{x}, t) = \frac{\delta f}{\delta \phi_n} = \frac{\partial f}{\partial \phi_n} - \boldsymbol{\nabla} \cdot \left(\frac{\partial f}{\partial \nabla \phi_n}\right)$$
$$\mu_d(\boldsymbol{x}, t) = \frac{\delta f}{\delta \phi_d} = \frac{\partial f}{\partial \phi_d} - \boldsymbol{\nabla} \cdot \left(\frac{\partial f}{\partial \nabla \phi_d}\right) \tag{S3}$$

Here, $\mu_n$ is the chemical potentials of nucleoplasm driving its kinetics as described in section S1.3 and $\mu_d$ is the chemical potential for the order parameter $\phi_d$ evolving the epigenetic marks in a conserved manner as discussed in the section S1.3 and S1.4.

*S1.3* *<u>Diffusion and reaction kinetics</u>*

The dynamic evolution of the nucleus is governed by both diffusion and reaction kinetics. First, we consider the nucleoplasm, the inert content within the nucleus. Its conservative evolution at all points in the nucleus is dependent on the local flux of nucleoplasm. This flux, as governed by Fick's first law, is directly proportional to the gradient of its chemical potential as $\boldsymbol{\mathcal{J}}^n = -M_n \nabla \mu_n$, where $M_n$ is the mobility of nucleoplasm in the nucleus. Thus, we can express the change in nucleoplasm volume fraction over time as a result of diffusion as

$$\frac{\partial \phi_n}{\partial t} = \underbrace{M_n \nabla^2 \mu_n}_{\text{diffusion}} \tag{S4}$$

The physical interpretation of Eq S4 which regulates the local redistribution of water particles is discussed in Section S1.4. On the other hand, the spatiotemporal evolution of chromatin – which distinguishes between the heterochromatin and the euchromatin phases – is governed by epigenetic reaction kinetics. The reactions facilitate a non-conservative regulation of the overall levels of histone methylation and acetylation. Heterochromatic phase, which is prominently methylated is converted into euchromatin phase via acetylation reaction involving the removal of methyl groups called demethylation followed by addition of acetyl group on the histone tails via acetyltransferase activity. The overall rate of acetylation reaction $\Gamma_{ac}$ is a combination of its substeps such that $\frac{1}{\Gamma_{ac}} = \frac{1}{\Gamma_{HDM}} + \frac{1}{\Gamma_{HAT}}$ (Fig 1B). Similarly, the euchromatin phase is converted into heterochromatin via methylation reaction incorporating histone deacetylation and methyltransferase activity such that $\frac{1}{\Gamma_{me}} = \frac{1}{\Gamma_{HDAC}} + \frac{1}{\Gamma_{HMT}}$ (Fig 1B). The overall rate of change of heterochromatin phase to euchromatin phase is captured via a first order kinetics of the reaction such that,

$$\left.\frac{\partial \phi_d}{\partial t}\right|_{\text{epigen}} = 2(\Gamma_{me}\phi_e - \Gamma_{ac}\phi_h) \tag{S5}$$

However, the rates of reactions ($\Gamma_{me}$ and $\Gamma_{ac}$) are not constant – but are dependent on the strength of interactions between neighboring nucleosomes. Briefly, the chromatin-chromatin energy landscape results in like-marked (heterochromatin-heterochromatin or euchromatin-euchromatin) neighbors being more stable than unlike-marked (heterochromatin-euchromatin) neighbors. Thus, reactions will preferentially occur if they lead to the formation of more like-marked nucleosome neighbors. The role of chromatin-chromatin interaction energetics in driving the spatially heterogenous reaction kinetics is discussed in more detail in Section S1.4. Effectively, such neighborhood dependent reaction kinetics emulates a diffusion-like conservative evolution of epigenetic marks – as if the epigenetic marks are spatially diffusing resulting in coarsening of the heterochromatin and euchromatin phases (as shown schematically in Fig S1 and explained in Section S1.4). This effectively conservative evolution can be written as,

$$\left.\frac{\partial \phi_d}{\partial t}\right|_{\text{cons}} = M_d \nabla^2 \mu_d \tag{S6}$$

Here, $M_d$ is the mobility of epigenetic marks in nucleus. The effective chemical potential of epigenetic marks that drives the diffusion-like epigenetic evolution is $\mu_d$ and is dependent on the free energy of chromatin organization, as described in Eq S3.

Combining these dynamics (Eq S5-S6), the overall evolution of the order parameter $\phi_d$ can be written as,

$$\frac{\partial \phi_d}{\partial t} = 2\underbrace{(\Gamma_{me}\phi_e - \Gamma_{ac}\phi_h)}_{\text{epigentic regulation}} + \underbrace{M_d \nabla^2 \mu_d}_{\text{diffusion}} \tag{S7}$$

This comprehensive framework of equations (S3), (S4), and (S7) provides a complete understanding of spatio-temporal evolution of chromatin organization within the nucleus.

#### *S1.4 Physical interpretation of diffusion and reaction kinetics*

As discussed in Section S1.3, we have incorporated the diffusion kinetics of nucleoplasm as well as the reaction kinetics of epigenetic regulation in our model. Mathematically, these are described by Eqs S4 and S7. In this section we the physical interpretation of the kinetics underlying diffusion and reaction driven chromatin evolution.

The nucleoplasm kinetics, given by Eq S4 is diffusion driven and conservative, i.e. unless there is a flux of water into or out of the nucleus, the total amount of nucleoplasm in the nucleus remains constant. The favorable energetic interactions, such as those mediated by nucleosome bridging proteins such as HP1$\alpha$ [3], results in methylated histones coming together. This compaction of chromatin requires local movement of water molecules away from the condensing heterochromatin domains, as shown in Fig S1. However, this spatiotemporal evolution does not alter the total number of water molecules. This energetic driven local conservative movement of water molecules is captured via the diffusion of nucleoplasm in Eq S4.

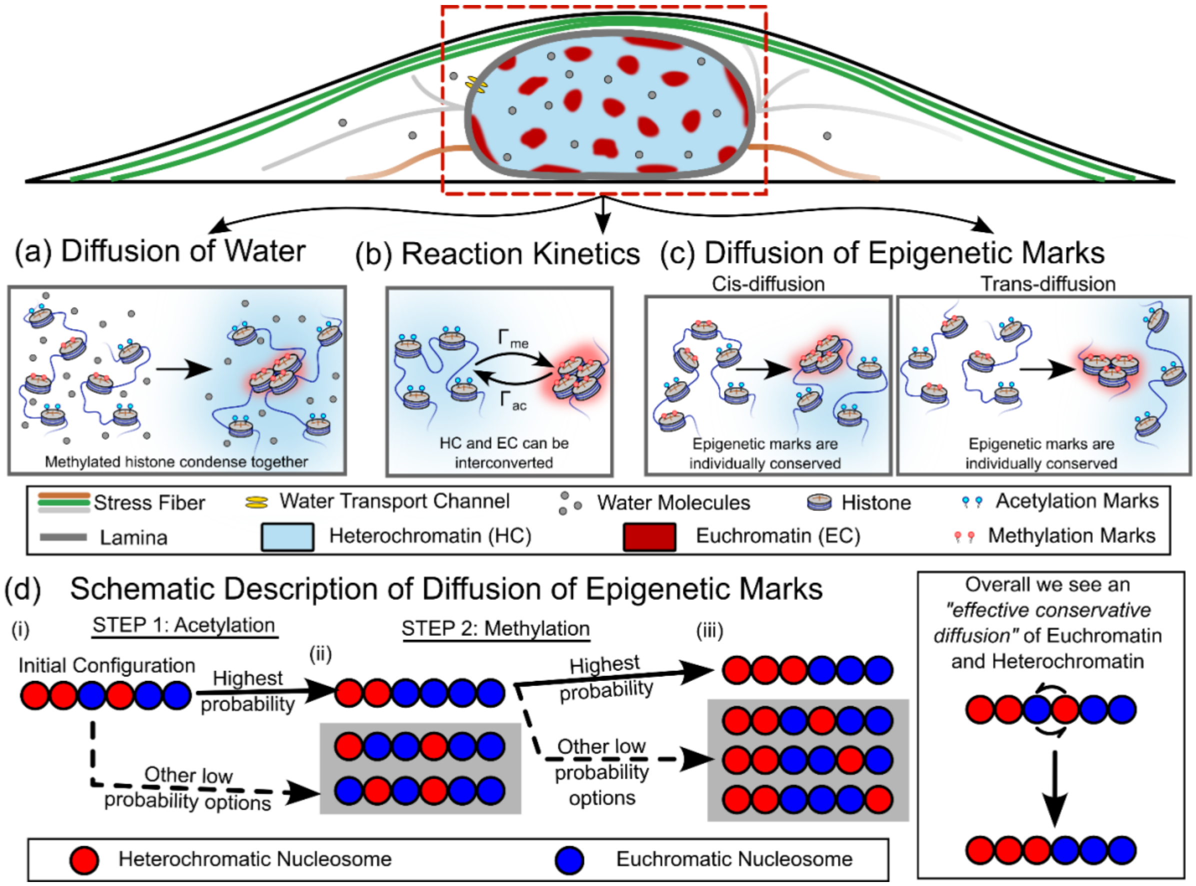


**Figure S1:** A schematic depicting the individual roles of the diffusion and reactions kinetics incorporated into the heterochromatin organization model. (a) Conservative diffusion of water which can redistribute molecules of water within the nucleus without changing the total amount of water, or total amount of hetero- or euchromatin in the nucleus. (b) Non-conservative reaction kinetics of histone acetylation and methylation, which allows an interconversion of chromatin phases. This changes the individual amounts of heterochromatin and euchromatin in the nucleus without changing the total amount of DNA. The reaction rates determine the ratio of heterochromatin to euchromatin at steady state. (c) The chromatin-chromatin interactions contribute to the reaction kinetics effectively driving a conservative evolution of epigenetic marks, which we call 'diffusion of epigenetic marks'. (d) The diffusion of epigenetic marks is an overall result of reaction kinetics coupled with preference of like-like neighbor over unlike ones, thereby effectively rendering reactions ate certain sites more probable than others. Adapted from Ref[4].

The kinetics of chromatin evolution (Eq S7) involves epigenetic regulation via the methylation or acetylation reactions and gives rise to the reaction kinetics (first term in Eq S7). The reaction kinetics are non-conservative as they involve interconversion of acetylated histones to methylated histones (methylation kinetics, $\Gamma_{me}$) and vice versa (acetylation kinetics, $\Gamma_{ac}$) as shown in Fig S1.

However, the reaction kinetics given by the first term in Eq S7 does not take into account the energetics of chromatin-chromatin interactions. As described in Eq S1.2, the chromatin energy landscape results in heterochromatin-heterochromatin or euchromatin-euchromatin neighbors being more stable than neighbors that are not like marked. As a result, if the methylation reaction results in conversion of a heterochromatin-heterochromatin neighbor into a heterochromatin-euchromatin neighbor, such conversion is energetically unfavorable. On the other hand, if reactions convert heterochromatin-euchromatin neighbors into a heterochromatin-heterochromatin or a euchromatin-euchromatin pair it is energetically favorable. Thus, the effective rates of epigenetic reactions are determined by the specific locations of the nucleosome where reactions occur dependent on the epigenetic marks of the neighboring nucleosomes.

A schematic evolution of epigenetic marks driven by such neighborhood dependent reaction rates is shown in Fig S1. At a given time, a portion of chromatin polymer has epigenetic marks as shown in Fig S1 (red: heterochromatin, blue: euchromatin). Let the first step involve acetylation reaction – converting a heterochromatic nucleosome into euchromatic form. While such conversion could happen at any of the nucleosomes, energetic interactions favor like-marked neighbors such that the configuration shown in Fig S1 has the highest probability of occurrence. If the next step involves further acetylation, in our model it will be captured by the non-conservative first term in Eq S7. So we only consider the case if the next step involves methylation – such that the number of heterochromatic and euchromatic nucleosomes are individually conserved. Again, due to energetic interactions favoring like-marked nucleosomes the highest probability of occurrence is for the configuration as shown in Fig S1. Thus, over the course of two steps in chromatin kinetics, we see that neighborhood dependent reaction kinetics can effectively result in a 'conservative' spatiotemporal evolution which leads to the coarsening of the two phases of chromatin. Such conservative evolution is effectively captured in our coarse-grained model via the second term in Eq S7 called the 'diffusion of epigenetic marks'.

Note that the sequence of events described here results in a cis-diffusion of epigenetic marks along the chain of chromatin polymer, as shown in Fig S1. Similar effective diffusion can occur between nucleosomes that are not neighbors along the polymer but are spatially close in the 3D space. This is called trans-diffusion of epigenetic marks (Fig S1). The diffusion term in our coarse-grained model (Eq S7, second term) is isotropic and involves both cis- and trans- diffusion mechanisms. The quantitative equivalence of the diffusion term used in Eq S7 and the neighborhood dependent reaction kinetics has been previously shown by us[4].

#### *S1.5 Rescaling the equations for numerical implementation*

The governing equations derived in Section S1.2 and S1.3 show an intrinsic length and time scale, which can be used for rescaling the equations. The rescaled equations are used in the numerical implementation of the solution of the governing equations. Here, we demonstrate the steps involved in obtaining a rescaled, non-dimensional set of governing equations by leveraging these inherent scales.

The reaction-diffusion kinetics described in Eq S7 yield a characteristic length scale determined by interplay of diffusion and reaction rates i.e. $\ell_{RD} = \sqrt{M_d/\Gamma_{ac}} = \sqrt{D/\Gamma_{ac}}$. Remarkably, another intrinsic length scale arises from the competition between interfacial and bulk mixing energies from Eq S1, i.e. the width of the interface $\ell_{int} = \sqrt{\kappa/c}$. In our simulations and theoretical analyses (discussed in Section S3), we find that the reaction-diffusion length significantly influences the sizes and spacing of heterochromatin domains. Consequently, we opt to rescale all lengths relative to $\ell_{RD}$, such that $\tilde{x} = x/\ell_{RD}$. Furthermore, the reaction rates provide an inherent time scale for the system. Therefore, we rescale all times as $\tilde{t} = t\Gamma_{ac}$. The coefficient $c$ , representing the energy of chromatin-chromatin interactions in Eq (S1), serves as an energy scaling factor. Thus, all energy densities are rescaled as $\tilde{f} = f/c$.

Rescaling Eq S1, we obtain

$$\tilde{f} = \underbrace{\frac{1}{2}\left[\phi_e^2 + \phi_h^2(\phi_{\mathrm{h}}^{\max} - \phi_h)^2\right]}_{\text{chromatin-chromatin interactions}} - \underbrace{\frac{\tilde{V}(\phi_h)}{d_0}\phi_h e^{-\frac{d}{d_0}}}_{\text{chromatin-lamina interactions}} + \underbrace{\frac{\delta^2}{2}|\nabla\phi_n|^2 + \frac{\delta^2}{2}|\nabla\phi_d|^2}_{\text{Interfacial energy}} \tag{S8}$$

Here, $\tilde{V}(\phi_h) = V(\phi_h)/c$ is the rescaled strength of chromatin-lamina anchoring interactions. The parameter $\delta = \frac{\ell_{int}}{\ell_{RD}}$ is a rescaled measure of the width of the interface.

The rescaled chemical potentials from Eq S3 expand as,

$$\tilde{\mu}_n(\tilde{\boldsymbol{x}},\tilde{t}) = -\phi_e - \phi_h(\phi_{h0} - \phi_h)(\phi_{h0} - 2\phi_h) - \frac{1}{2}\frac{\tilde{V}(\phi_h)}{d_0} e^{-\frac{d}{d_0}} - \delta^2\nabla^2\phi_n \tag{S9}$$

$$\tilde{\mu}_d(\tilde{\boldsymbol{x}},\tilde{t}) = -\phi_e + \phi_h(\phi_{h0} - \phi_h)(\phi_{h0} - 2\phi_h) - \frac{1}{2}\frac{\tilde{V}(\phi_h)}{d_0} e^{-\frac{d}{d_0}} - \delta^2\nabla^2\phi_d$$

Lastly, we rescale the kinetics equations such that,

$$\frac{\partial \phi_n}{\partial \tilde{t}} = \underbrace{\nabla^2\tilde{\mu}_n}_{\text{diffusion}} \tag{S10}$$

$$\frac{\partial \phi_d}{\partial \tilde{t}} = \underbrace{\nabla^2\tilde{\mu}_d}_{\text{diffusion}} + \underbrace{2(\tilde{\Gamma}_{me}\phi_e - \phi_h)}_{\substack{\text{epigenetic}\\ \text{regulation}}}$$

Here, all reaction rates have been rescaled with respect to the time scale such that $\tilde{\Gamma}_{me} = \frac{\Gamma_{me}}{\Gamma_{ac}}$. These rescaled equations, Eq (S8) and Eq (S10), are simultaneously solved numerically using COMSOL Multiphysics, subject to boundary conditions ensuring no flux of nucleoplasm or epigenetic marks across all boundaries (see section S6 for more details).

### S2 Average chromatin phase contents is determined by epigenetic reactions

Here we show that the average amount of eu- and heterochromatin in nucleus is determined solely by the epigenetic reactions. The spatiotemporal evolution of order parameter $\phi_d$ due to diffusion of epigenetic marks and kinetics of interconversion of eu- and heterochromatin phases, as described by Eq S7 as,

$$\frac{\partial \phi_d}{\partial t} = M_d\nabla^2\mu_d + 2(\Gamma_{me}\phi_e - \Gamma_{ac}\phi_h) \tag{S11}$$

At steady state, $\frac{\partial \phi_d}{\partial t} = 0$, the relation between the average euchromatin content $\bar{\phi}_e$ and the average heterochromatin $\bar{\phi}_h$ content in the nucleus can be obtained by averaging Eq S11 as,

$$0 = M_d \frac{\int_{\mathrm{V_n}} \nabla^2\mu_d \, dV}{\int_{\mathrm{V_n}} dV} + 2(\Gamma_{me}\bar{\phi}_e - \Gamma_{ac}\bar{\phi}_h)$$

The term $\nabla^2\mu_d$ (first term on the right-hand side) on averaging over the entire volume of the nucleus can be approximated to zero. Thus,

$$0 = 2(\Gamma_{me}\bar{\phi}_e - \Gamma_{ac}\bar{\phi}_h)$$

Given that $\phi_e + \phi_h + \phi_n = 1$, we obtain the following relationship:

$$(\Gamma_{me} + \Gamma_{ac})\bar{\phi}_h = \Gamma_{me}(1 - \bar{\phi}_n)$$

Or,

$$\bar{\phi}_h = \frac{\Gamma_{me}\,(1 - \bar{\phi}_n)}{\Gamma_{me} + \Gamma_{ac}}, \qquad \bar{\phi}_e = \frac{\Gamma_{ac}\,(1 - \bar{\phi}_n)}{\Gamma_{me} + \Gamma_{ac}} \tag{S12}$$

After rescaling, the average eu- and heterochromatin contents obtained via Eq S12 can be expressed as

$$\bar{\phi}_h = \frac{\tilde{\Gamma}_{me}\,(1-\bar{\phi}_n)}{\tilde{\Gamma}_{me}+1}, \qquad \bar{\phi}_e = \frac{(1-\bar{\phi}_n)}{\tilde{\Gamma}_{me}+1} \tag{S13}$$

### S3 Characteristic Heterochromatin Domain size determination in presence of Reactions

The reaction kinetic parameters $(\Gamma_{me}, \Gamma_{ac})$ not only determine the average amounts of heterochromatin and euchromatin in the nucleus (Eq S12, S13), but also influence the sizes of the heterochromatin domains, as shown in Section 3.1 of the main manuscript. Here, we show the detailed derivation of interior heterochromatin domain size away from nuclear periphery. To analyze the steady state size of the heterochromatin domain, we first examine the chromatin composition within and around the droplet (Fig S2). As discussed in Section S2, the acetylation and methylation together determine the mean heterochromatin (and euchromatin) content in the nucleus, given by Eq S12. In absence of any energetic considerations, this would give rise to a homogeneous mean chromatin composition $(\bar{\phi}_h, \bar{\phi}_e)$. However, this composition (Fig 1C; white circle) lies in neither of the energy wells and is thus energetically unfavorable. To reduce the total free energy, the system phase separates via nucleation of heterochromatin droplets (Fig S2). Due to phase separation, the heterochromatin volume fraction immediately outside the droplet is $\phi_h \approx 0$ corresponding to the euchromatic energy well. Far away from the droplet, the mean composition $(\bar{\phi}_h, \bar{\phi}_e)$ remains undisturbed. However, when there is a significant peripheral sequestering of methylated histones, the far-field concentration of heterochromatin $\bar{\phi}_h^*$ (Fig S2), drops from the average concentration $\bar{\phi}_h$ (discussed in Section S4).

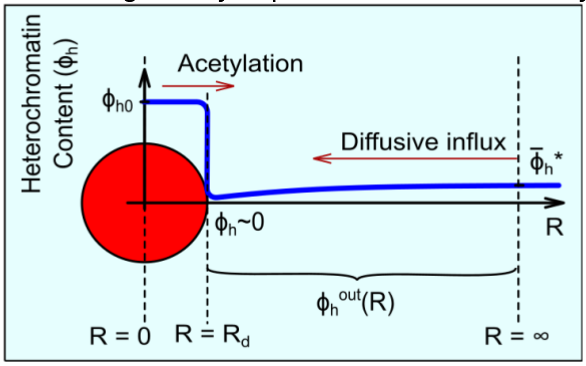


**Figure S2:** The competition of diffusion driven influx of heterochromatin with the epigenetic reaction driven outflux of heterochromatin from the heterochromatin domain determines its steady state size. The figure also shows the radial distribution of heterochromatin volume fraction $\phi_h$ in and around the domain.

In the dilute limit, where there is significantly more euchromatin than heterochromatin, the droplet size can be considered much smaller than the spacing between domains such that neighboring droplets are far enough to not interact with each other. Under such assumption the heterochromatin distribution becomes spherically symmetric. By adopting a polar coordinate system with origin at the droplet center, we can write the steady-state concentration field $\phi_h(R)$ as a function of distance to the center of the droplet, $R$ as

$$\phi_h(R) = \begin{cases} \phi_{h0} & R < R_d \\ \phi_h^+ = \dfrac{\kappa}{R_d} & R = R_d^+ \\ \bar{\phi}_h^* & R = \infty \end{cases}$$

where $R_d$ is the radius of the droplet and $\kappa$ is the measure of interfacial energy. Fig S2 shows the distribution of heterochromatin content $\phi_h(R)$ around a spherical heterochromatin domain (red) of radius $R_d$ as it grows surrounded by euchromatin phase (in light blue). At steady state, heterochromatin composition outside the droplet follows the equation,

$$D_h \nabla^2 \phi_h^{out} - \Gamma_{ac}\phi_h^{out} + \Gamma_{me}\phi_e = 0 \tag{S14}$$

Here $D_h = M_h c$ is the diffusivity of heterochromatin in the nucleoplasm. By solving Eq S14 outside the heterochromatin domain, with boundary conditions $\phi_h|_{R_d^+} = \kappa/R_d$ and $\phi_h|_\infty = \bar{\phi}_h^*$, we get

$$\phi_h^{out}(R) = \bar{\phi}_h^* + (\phi_h^+ - \bar{\phi}_h^*)\frac{R_d}{R}e^{\frac{R_d - R}{\ell_{RD}}}$$

where $\ell_{RD}$ is the characteristic reaction-diffusion length scale given under a dilute limit as $\ell_{RD} = \sqrt{\frac{D_h}{\Gamma_{ac}}}$. Thus,

$$\frac{\partial \phi_h^{out}}{\partial R} = (\bar{\phi}_h^* - \phi_h^+)\frac{R_d(\ell_{RD} + R)}{\ell_{RD}R^2}e^{\frac{R_d - R}{\ell_{RD}}}$$

Thus,

$$\left.\frac{\partial \phi_h^{out}}{\partial R}\right|_{R \to R_d} = \frac{\bar{\phi}_h^*}{R_d} - \frac{\kappa}{R_d^2} \qquad \left(when \frac{R_d}{\ell_{RD}} \ll 1\right) \tag{S15}$$

Having obtained the heterochromatin concentration field, we next discuss the growth dynamics of heterochromatin droplet. The droplet grows due to the reaction-diffusion driven influx (blue curve in Fig S2) of heterochromatin. On the other hand, within the heterochromatin droplet (with $\phi_h = \phi_{h0}$) histone acetylation reactions will allow conversion of heterochromatin inside the droplet into euchromatin outside. This acetylation driven outflux oppose the diffusive influx of heterochromatin and thereby reduce the size of the droplet (Fig S2). Thus, the rate of change of the volume of the droplet $V_d$ can be written as,

$$\frac{dV_d}{dt} = \frac{4}{3}\pi\frac{dR_d^3}{dt} = \underbrace{J^{in}}_{\text{inwards diffusion}} - \underbrace{\Gamma_{ac} \times \frac{4}{3}\pi R_d^3 \phi_{h0}}_{\text{Acetylation}} \tag{S16}$$

where the diffusive influx of heterochromatin is,

$$J^{in} = 4\pi R_d^2 D_h \left.\frac{\partial \phi_h^{out}}{\partial R}\right|_{R \to R_d} \tag{S17}$$

Using the spatial gradient from Eq S15, we simplify Eq S16 as,

$$4\pi R_d^2 \frac{dR_d}{dt} = 4\pi D_h(\bar{\phi}_h^* R_d - \kappa) - \Gamma_{ac} \times \frac{4}{3}\pi R_d^3 \phi_{h0}$$

$$\frac{dR_d}{dt} = D_h\left(\frac{\bar{\phi}_h^*}{R_d} - \frac{\kappa}{R_d^2}\right) - \frac{\Gamma_{ac}}{3}R_d\phi_{h0} \tag{S18}$$

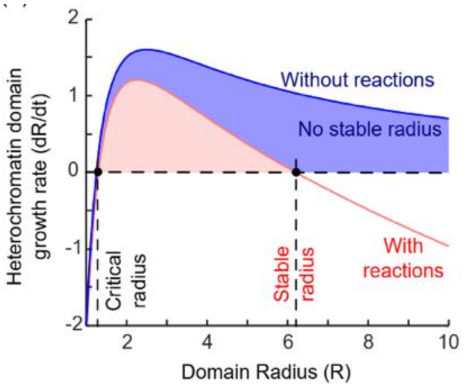


**Figure S3:** The growth rate of the heterochromatin domain.

Using Eq S18, we plot the rate of change of heterochromatin domain size with respect to the instantaneous domain radius (Fig S3). Above a critical radius, all heterochromatin domains grow ($dR_d/dt > 0$). In the absence of reactions ($\Gamma_{ac} = 0$, blue curve), the rate of change in the heterochromatin domain radius is always positive indicating that the domain will keep growing as long as its radius is larger than the critical radius. However, in the presence of the reactions, the domains grow until their growth rate reaches a zero value. This gives the stable size of heterochromatin domains. The domains larger than the stable radius will shrink back to the stable radius. The stable radius ($R_d^s$) can be obtained by setting $dR_d/dt = 0$ in Eq S18 such that,

$$0 = \frac{D_h \bar{\phi}_h^*}{\phi_{h0}} - \frac{\Gamma_{ac}}{3} {R_d^s}^2$$

Thus,

$$R_d^s = \sqrt{\frac{3D_h}{\Gamma_{ac}\phi_{h0}} \bar{\phi}_h^*} \qquad \text{(S19)}$$

On rescaling, Eq S19 become,

$$\tilde{R}_d^s = \sqrt{\frac{3}{\phi_{h0}} \bar{\phi}_h^*} \qquad \text{(S20)}$$

All lengths are rescaled with respect to $\ell_{RD}$, while all times with respect to $1/\Gamma_{ac}$. Eq S20 give a non-dimensional dependence of heterochromatin domain size on the epigenetic kinetics.

In deriving the domain size scaling (Eq S19), we have considered a 3-dimensional scenario where spheroids of heterochromatin domains are formed. However, for consistency with our simulations and the STORM images, the derivation can be easily repeated for the 2D case – to obtain the same scaling relations as in 3D case. In 2D, Eq. S14 can be rewritten as

$$D_h \left( \frac{\partial^2 \phi_h}{\partial r^2} + \frac{1}{r}\frac{\partial \phi_h}{\partial r} \right) - \Gamma_{ac}\phi_h + \Gamma_{me}\phi_e = 0$$

which is the modified Bessel's equation of order 0. The solution to this equation is

$$\phi_h = \bar{\phi}_h^* + (\phi_h^+ - \phi_{h0}) \frac{K_0(r/l_{RD})}{K_0(R_d/l_{RD})}$$

where $K_0$ denotes the zeroth order Bessel K function. Also, in 2D Eq. S16 becomes

$$\frac{dV_d}{dt} = 2\pi R_d \frac{dR_d}{dt} = \underbrace{-2\pi R_d D_h \frac{\partial \phi_h^{out}}{\partial r}}_{diffusive\ flux} - \underbrace{\pi R_d^2 \Gamma_{ac}}_{acetylation} \phi_{h0} \qquad \text{(S21)}$$

Solving Eq. S21 at the steady state $\frac{dV_d}{dt} = 0$ yields a scaling relation between the radius of heterochromatin domain and the level of methylation,

$$\frac{K_1(R_d^s/l_{RD})}{K_0(R_d^s/l_{RD})} \sim \frac{2D_h}{\Gamma_{ac}\phi_{h0}}\ \bar{\phi}_h^*$$

Thus, we obtain the same scaling relation $R_d^s = f\left(\frac{D_h}{\Gamma_{ac}\phi_{h0}}\bar{\phi}_h^*\right)$, where $f$ is a function determined from Eq. S21. In the range of our parameters, $R_d^s \sim \sqrt{\frac{D_h}{\Gamma_{ac}\phi_{ho}}\bar{\phi}_h^*}$ provides a good approximation for the scaling of the size of heterochromatin domains, as confirmed by our numerical simulations (Fig 2E).

As discussed in the beginning of this section, $\bar{\phi}_h^*$ is the far-field heterochromatin concentration for the interior of the nucleus. The peripheral sequestering of heterochromatin towards the nuclear lamina governs this value, as we discuss next.

### S4 Far-field Heterochromatin concentration is influenced by Peripheral HC Sequestering

The competition between interior and peripheral heterochromatin domains for the available DNA pool within the nucleus shapes the overall heterochromatin distribution. As discussed in Section 3.1 of the main manuscript, this competition is expressed as:

$$Total\ HC = HC^{interior} + HC^{Periphery} \tag{S22}$$

Therefore, when there is a significant peripheral sequestering of heterochromatin, the far-field concentration of heterochromatin $\bar{\phi}_h^*$ (Fig S2), drops from the reaction rate determined average concentration $\bar{\phi}_h \approx \frac{\Gamma_{me}\,(1-\bar{\phi}_n)}{\Gamma_{me}+\Gamma_{ac}}$.

In Eq S22, 'Total HC' is the average heterochromatin content in the nucleus $\bar{\phi}_h$ times the total nuclear area, $HC^{interior}$ is obtained by total number of interior domain times the area of one domain and $HC^{Periphery}$ is an estimate of heterochromatin accumulated at periphery, such that

$$\underbrace{\pi R_{nuc}^2}_{\substack{Area\ of\\ Nucleus}} * \underbrace{\bar{\phi}_h}_{\substack{Average\ HC\\ in\\ Nucleus}} = \underbrace{\frac{\alpha\pi R_{nuc}^2}{\pi l_{RD}^2}}_{\substack{Number\ of\\ interior\ HC\\ domains}} * \underbrace{\pi R_d^2}_{\substack{Area\ of\ one\\ interior\ HC\\ domain}} + \underbrace{2\pi R_{nuc}}_{\substack{Periphery\ of\\ Nucleus}} * \underbrace{T_{LAD}}_{\substack{Thickness\ of\\ peripheral\ HC}} \tag{S23}$$

Here, $R_{nuc}$ is the radius of the nucleus and $T_{LAD}$ is the size of peripheral heterochromatin domain (details in Section S5). The parameter $\alpha$, analogous to the atomic packing factor, gives an estimate of fraction of volume occupied by the heterochromatin in nucleus interior. On using Eq 19, Eq S23 gives

$$\bar{\phi}_h = \frac{3\alpha}{\phi_{ho}}\ \bar{\phi}_h^* + \frac{2T_{LAD}}{R_{nuc}}$$

Here $\frac{3\alpha}{\phi_{ho}} \approx 1$, thus

$$\bar{\phi}_h^* = \bar{\phi}_h - \frac{2T_{LAD}}{R_{nuc}} \tag{S24}$$

On rescaling, Eq S24 becomes,

$$\bar{\phi}_h^* = \frac{\tilde{\Gamma}_{me}}{\tilde{\Gamma}_{me}+1} - \frac{2\tilde{T}_{LAD}}{\tilde{R}_{nuc}} \tag{S25}$$

Here, $\tilde{T}_{LAD}$ is the rescaled size of peripheral heterochromatin domain and $\tilde{R}_{nuc}$ is the normalized radius of the nucleus. Note that when the LAD thickness is very small $\left(\tilde{T}_{LAD} \ll \tilde{R}_{nuc}\right)$, the influence of peripheral domains on interior domains is insignificant and $\bar{\phi}_h^* = \bar{\phi}_h$.

### S5 LAD morphology is regulated by chromatin-lamina affinity and epigenetic reactions

#### *S5.1 LAD shapes are determined by chromatin-lamina affinity*

As discussed in section 3.1, LADs shapes are determined by the competition between chromatin-chromatin interactions and chromatin -lamina interactions as

$$\theta = \cos^{-1}\left(\frac{\gamma_{lh} - \gamma_{le}}{\gamma_{he}}\right)$$

where $\theta$ is the contact angle of LAD with nuclear lamina, $\gamma_{lh}$ and $\gamma_{le}$ are the surface tension at lamina-heterochromatin and lamina-euchromatin interfaces and $\gamma_{he}$ is the surface tension at heterochromatin-euchromatin interface (Fig S4). Next, we drive the relation between surface tension at lamina-chromatin interface and lamina-chromatin interaction strength.

The total free energy over the entire volume from Eq S1 can be written as

$$\Pi = \int_V \left[\Delta F(\phi_h, \phi_e) + \kappa\left[\left(\frac{d\phi_h}{dx}\right)^2 + \left(\frac{d\phi_e}{dx}\right)^2\right] + \underbrace{V(\phi_h^s)}_{energy/area}\ \delta(0)\right] dV \tag{S26}$$

Here $\phi_h^s$ is the volume fraction of heterochromatin at the lamina interface. $V(\phi_h^s)$ is a general function, a measure of the chromatin-lamina interaction energy per unit area, with value $V_{EC}$ when in euchromatin phase (when $\phi_h^s = 0$) and $V_{HC}$ when in heterochromatin phase (when $\phi_h^s = 1$). We consider two regions – (i) where heterochromatin interacts with lamina, and (ii) where euchromatin interacts with lamina.

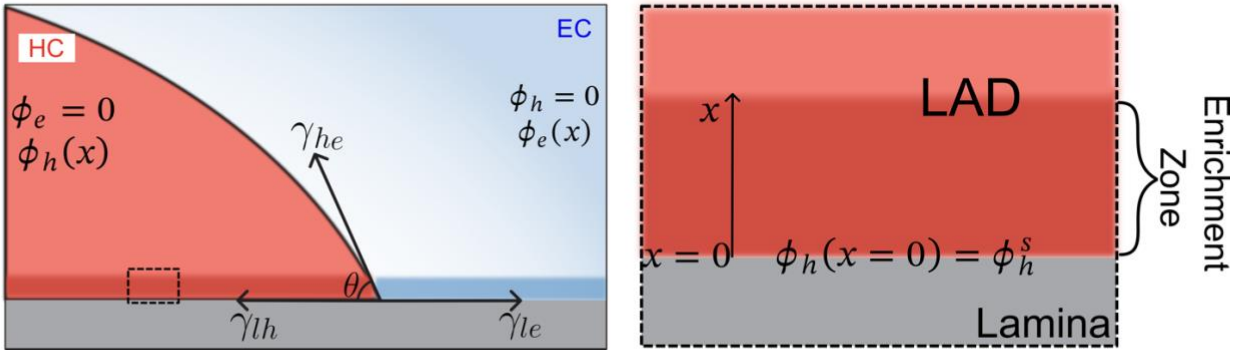


**Figure S4:** The balance of surface tensions at the interface of chromatin-lamina ($\gamma_{lh}$and $\gamma_{le}$) and the two phases of chromatin ($\gamma_{he}$) gives rise to a stable morphology of LADs, which have a characteristic contact angle with the lamina($\theta$). The right panel shows the zoomed in view of heterochromatin enriched zone at heterochromatin-lamina interface.

First consider the region (i) where $\phi_e = 0$. The total free energy in this region from Eq S26 is

$$\Pi = \int_0^x \left[\Delta F(\phi_h) + \kappa\left[\left(\frac{d\phi_h}{dx}\right)^2\right] + V(\phi_h^s)\delta(0)\right] dx \tag{S27}$$

On simplification,

$$\Pi = V(\phi_h^s) + \int_0^x \left[\Delta F(\phi_h) + \kappa\left[\left(\frac{d\phi_h}{dx}\right)^2\right]\right] dx$$

In region (i), the minima of total free energy $\Pi$ occurs when,

$$\Delta F(\phi_h) = \kappa\left(\frac{d\phi_h}{dx}\right)^2 \tag{S28}$$

Thus, everywhere in the region (i):

$$\frac{d\phi_h}{dx} = \pm\sqrt{\frac{\Delta F(\phi_h)}{\kappa}} \tag{S29}$$

With boundary conditions (at x=0),

$$\frac{dV(\phi_h^s)}{d\phi_h^s} = 2\kappa\left.\frac{d\phi_h}{dx}\right|_{x=0} = \pm 2\sqrt{\kappa\Delta F(\phi_h^s)}$$

Now the surface tension i.e the free energy at lamina-heterochromatin interface

$$\gamma_{lh} = \underbrace{V(\phi_h^s)}_{=V^{HC}} + \int_0^\infty \left[\Delta F(\phi_h) + \kappa\left[\left(\frac{d\phi_h}{dx}\right)^2\right]\right] dx$$

Use Eq S28 and S29:

$$\gamma_{lh} = V^{HC} + \int_1^{\phi_h^s} 2\Delta F(\phi_h)\sqrt{\frac{\kappa}{\Delta F(\phi_h)}}\, d\phi_h$$

$$\gamma_{lh} = \underbrace{V^{HC}}_{lamina\ interaction\ strength} + \underbrace{2\int_1^{\phi_h^s} \sqrt{\kappa\Delta F(\phi_h)}\, d\phi_h}_{Excess\ energy\ due\ to\ enrichment} \tag{S30}$$

Similarly in region (ii),

$$\gamma_{le} = \underbrace{V^{EC}}_{lamina\ interaction\ strength} + \underbrace{2\int_1^{\phi_e^s} \sqrt{\kappa\Delta F(\phi_e)}\, d\phi_e}_{Excess\ energy\ due\ to\ enrichment} \tag{S31}$$

Balancing the surface tensions:

$$\gamma_{he}\cos\theta = \gamma_{lh} - \gamma_{le}$$

$$\gamma_{he}\cos\theta = V_{HC} - V_{EC} = V_{LAD}$$

Notably the integral in right side in Eq S30 and S31 accounting for the excess energy due to enrichment at interface is negligible. We are extracting this difference $V_{LAD}$ – the preferential interactions of lamina with the heterochromatin over euchromatin.

*S5.2* *LADs thickness is regulated by both chromatin-lamina affinity and epigenetic reactions*

As discussed in Section 3.1, chromatin-lamina affinity and epigenetic reactions synergistically determine the characteristic length-scales of the peripheral heterochromatin domains.

As in the interior of the nucleus, the free energy of chromatin organization can be lowered by nucleation of compacted heterochromatin domains ($\phi_h = \phi_{h0}$) at the nuclear periphery, immediately surrounded by euchromatin ($\phi_h = 0$). We consider the nucleation and growth of a single peripheral heterochromatin domain (Fig 2F) of radius, $R_{LAD}$, at nuclear periphery forming a contact angle $\theta$ with lamina. Far from the peripheral heterochromatin domain, the average heterochromatin volume fraction remains undisturbed at the reaction determined level $\bar{\phi}_h^*$. Similar to interior domains, at steady state, heterochromatin composition outside the peripheral domain follows the equation

$$D_h\left(\frac{\partial^2\phi_h}{\partial r^2} + \frac{1}{r}\frac{\partial\phi_h}{\partial r}\right) - (\Gamma_{ac} + \Gamma_{me})\phi_h + \Gamma_{me} = 0 \tag{S32}$$

with solution

$$\phi_h = \bar{\phi}_h^* + (\phi_h^+ - \phi_{h0})\frac{K_0(r/l_{RD})}{K_0\left(\frac{R_{LAD}}{l_{RD}}\right)} \approx \bar{\phi}_h^*\left[1 - f\left(\frac{r}{l_{RD}}, \frac{R_{LAD}}{l_{RD}}\right)\right]$$

Thus,

$$\frac{\partial\phi_h}{\partial r}|_{R_{LAD}} = \bar{\phi}_h^*\frac{\partial f}{\partial r}|_{R_{LAD}}$$

Similar to interior domains, the concentration gradient driven diffusive influx (blue curve in Fig 2F) is opposed by acetylation-driven effective outflux of methyl marks from within the peripheral domains. Thus, the rate of change of the LAD size $A_{LAD}$ can be written as

$$\frac{dA_{LAD}}{dt} = \underbrace{D_h\int_0^\theta \frac{\partial\phi_h}{\partial r}|_{R_{LAD}}\,d\theta}_{influx} - \underbrace{A_{LAD}\Gamma_{ac}}_{outflux} \tag{S33}$$

Note that, the reaction-diffusion influx depends on the heterochromatin-lamina contact angle $\theta$ and the peripheral heterochromatin domain size $R_{LAD}/l_{RD}$. Thus, at steady state

$$\frac{dA_{LAD}}{dt}|_{ss} = 0 = D_h\int_0^\theta \bar{\phi}_h^*\frac{\partial f}{\partial r}|_{R_{LAD}}\,d\theta - A_{LAD}\Gamma_{ac}$$

On simplification,

$$0 = D_h\bar{\phi}_h^* F\left(\frac{R_{LAD}}{l_{RD}}, \theta\right) - R_{LAD}^2(\theta - sin\theta cos\theta)\Gamma_{ac}$$

Thus

$$R_{LAD} = \sqrt{\frac{D_h}{\Gamma_{ac}} \bar{\phi}_h^* \frac{F\left(\frac{R_{LAD}}{l_{RD}}, \theta\right)}{a(\theta)}} \quad \text{(S34)}$$

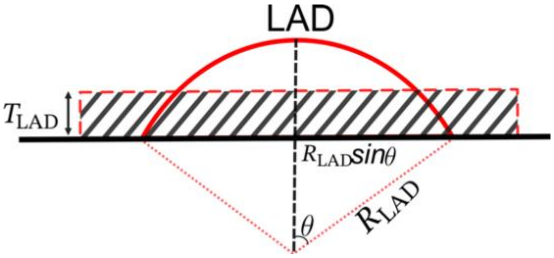

**Figure S5:** A schematic illustrating the transformation of LAD radius $R_{LAD}$ into LAD thickness $T_{LAD}$.

Here, the function $a(\theta) = (\theta - sin\theta cos\theta)$ quantifies the area of the stable LAD. For a unified quantification of the individual peripheral domains, we define the length-scale of a LAD $T_{LAD}$, as its height averaged over its span as shown in Fig S5. Note that $T_{LAD}$ is equivalent to the average thickness of the LAD and is inclusive of the previously defined LAD morphometric parameters $\theta$ and $R_{LAD}$ such that

$$T_{LAD} = \frac{R_{LAD}^2 a(\theta)}{l_{RD}} \quad \text{(S35)}$$

So

$$T_{LAD} = \sqrt{\frac{D_h}{\Gamma_{ac}} \bar{\phi}_h^* F\left(\frac{R_{LAD}}{l_{RD}}, \theta\right)}$$

The unified quantification of LADs via its length-scale $T_{LAD}$ is particularly advantageous for quantifying the LADs observed via STORM images, where due to resolution limits, there may be a significant level of stochasticity in the measurement of individual contact angles of each LAD. Using $\bar{\phi}_h^*$ from Eq S24

$$T_{LAD} = \sqrt{\frac{D_h}{\Gamma_{ac}} F\left(\frac{R_{LAD}}{l_{RD}}, \theta\right) \left(\frac{\Gamma_{me}}{\Gamma_{me} + \Gamma_{ac}} - \frac{2T_{LAD}}{R_{nuc}}\right)}$$

Thus

$$T_{LAD} = \sqrt{\frac{D_h}{\Gamma_{ac}} \frac{\Gamma_{me}}{\Gamma_{me} + \Gamma_{ac}} \frac{F\left(\frac{R_{LAD}}{l_{RD}}, \theta\right)}{\left(1 + \frac{2l_{RD}}{R_{Nuc}} F\left(\frac{R_{LAD}}{l_{RD}}, \theta\right)\right)}}$$

On simplification

$$T_{LAD} \sim \sqrt{\frac{D_h}{\Gamma_{ac}} \frac{\Gamma_{me}}{\Gamma_{me} + \Gamma_{ac}} G\left(\frac{R_{LAD}}{l_{RD}}, \theta\right)} \quad \text{(S36)}$$

Here, $G\left(\frac{R_{LAD}}{l_{RD}}, \theta\right)$ is a function that measures the total diffusive influx of methyl marks and is contingent on both the LAD-lamina contact angle $\theta$ and the LAD radius $R_{LAD}/l_{RD}$. The function

$G\left(\frac{R_{LAD}}{l_{RD}},\theta\right)$ can be obtained accurately numerically to obtain the dependence of LAD thickness on epigenetic rates $(\Gamma_{me},\Gamma_{ac})$ and chromatin-lamina affinity $(V_{LAD} \sim \gamma_{he}\cos\theta)$ as shown in next section.

### S6 Numerical Solution for LAD Thickness

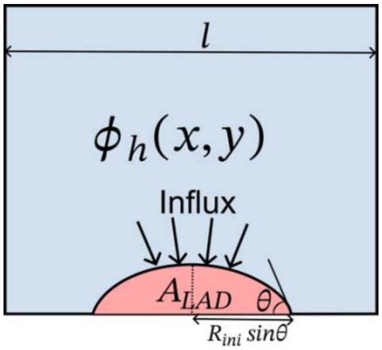


**Figure S6:** A schematic representation of geometry used to obtain LAD thickness numerically.

To get the precise dependence of LAD thickness $T_{LAD}$ on histone methylation rate $\Gamma_{me}$ and chromatin-lamina affinity relative to chromatin-chromatin interaction strength $V_{LAD}$ $(\sim \gamma_{\mathrm{he}}\cos\theta)$, we start with a LAD domain of initial guess radius $R_{ini}$ at a given histone methylation rate $\Gamma_{me}$ and a constant LAD-lamina contact angle $\theta$ (Fig S6). The rate of change of the LAD size $A_{LAD}$ is described by the equation:

$$\frac{dA_{LAD}}{dt} = D_h\nabla^2\phi_h - (\Gamma_{ac}+\Gamma_{me})\phi_h + \Gamma_{me} \quad \text{(S36)}$$

We solve Eq S36 numerically for $\phi_h$ to determine the flux. At steady-state, the balance between influx and outflux-given $D_h\nabla^2\phi_h$ for influx and $A_{LAD}\Gamma_{ac}$ for outflux-yields $A_{LAD}$. The updated LAD radius $R'$ is calculated from $A_{LAD}$ using: $R' = \sqrt{\frac{A_{LAD}}{\theta - 0.5 sin2\theta}}$. We then update the $R_{ini}$ with $R'$ and repeat the iterations (Fig S7) until convergence is achieved at an error threshold of 0.001. The final converged radius $R_{LAD}$, is used to calculate LAD thickness $T_{LAD}$ via

$$T_{LAD} = \frac{R_{LAD}^2(\theta - \sin\theta\cos\theta)}{l} \quad \text{(S37)}$$

To obtain the dependence of $T_{LAD}$ on $V_{LAD}$ and $\Gamma_{me}$, we conducted a series of above discussed numerical simulations varying $\theta$ and $\Gamma_{me}$. The results, depicted in Fig S8, show three distinguishable regimes in LAD thickness variation (Fig S8, inset) as the strength or number of chromatin anchoring proteins increases. In the next section we discuss these three regimes in detail.

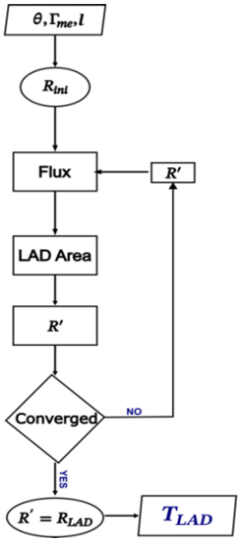


**Figure S7:** A flowchart showing the steps involved in numerical estimation of LAD thickness $T_{LAD}$.

### S7 Three regimes of LAD thickness $T_{LAD}$ variation with chromatin-lamina affinity $V_{LAD}$

As discussed in Section S6, the dependence of LAD thickness on chromatin-lamina affinity shows three regimes:

Regime I: Weak chromatin-lamina affinity: When the chromatin-lamina interactions at the periphery are weak, the interactions between distinctly and like-marked histones dominate, resulting in formation of bead-like peripheral heterochromatin domains with contact angles $\theta \approx 90°$ (Fig 2C). Due to a sparse distribution of peripheral domains along the lamina, owing to low chromatin-lamina affinity, the average LAD thickness (Fig 2G, top panel) is small. While a small increase in chromatin-lamina affinity will increase the spreading of LADs, their sparse distribution with regions of lamina not associating with chromatin results in a relatively negligible change in the LAD thickness. Indeed, as shown in Fig S8, within the Regime I the numerically predicted average LAD thickness does not change significantly with increase in chromatin-lamina affinity. On the other hand, as methylation rate increases (or acetylation rate decreases), there is an overall augmentation in heterochromatin content in the nucleus and at its periphery, leading to an increase in LAD thickness (Fig S8). Within regime I (i.e. when $\theta \sim 90°$), under dilute limit condition where LADs are very distant from each other, the discrete LADs are much smaller than the relevant reaction-diffusion length scale $R_{LAD} \ll l_{RD}$, $G\left(\frac{R_{LAD}}{l_{RD}}, \theta\right) \sim G(0, \theta)$ is a finite function of the contact angle $\theta$, independent of $R_{LAD}$. In this limit, the dependence of LAD thickness on epigenetic reaction rate $\Gamma_{me}$ scales as

$$T_{LAD} \sim \sqrt{\frac{D_h}{\Gamma_{ac}} \frac{\Gamma_{me}}{\Gamma_{ac} + \Gamma_{me}}} \tag{S37}$$

and shows excellent agreement with the numerical solution of Eq S36 (Fig S9).

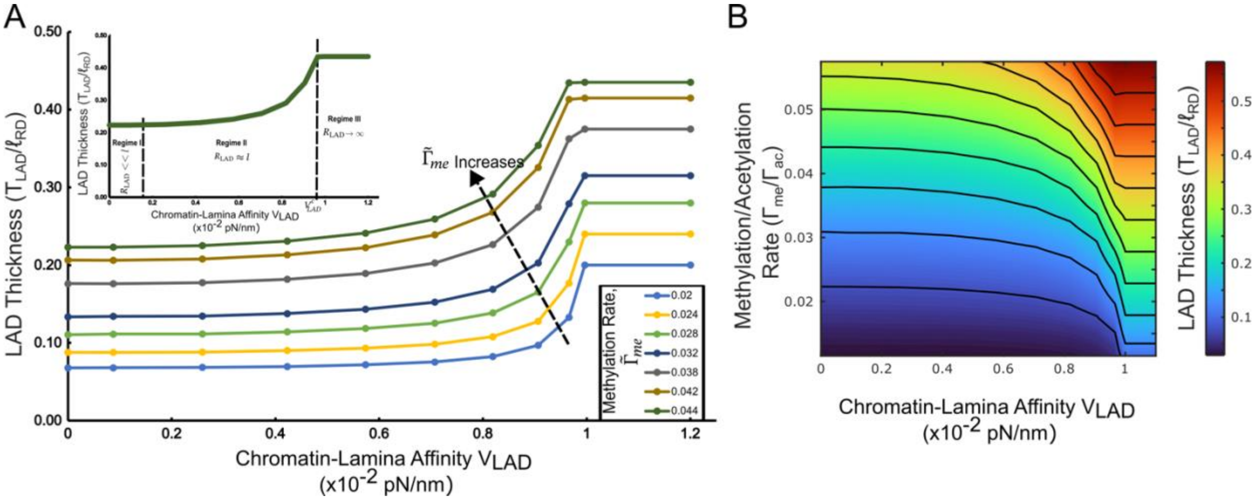


**Figure S8: The epigenetic reaction rates $(\Gamma_{me}/\Gamma_{ac})$ and the chromatin-lamina affinity $(V_{LAD})$ together determine the thickness of the LADs $(T_{LAD})$.** The numerically obtained dependance of LAD thickness on chromatin-lamina affinity and histone methylation rate. For a given $V_{LAD}$ as methylation increases the LAD thickness increases. For a given methylation rate the LAD thickness increases with chromatin-lamina affinity. However, beyond a value of $V_{LAD}$ any increase in methylation rate does not affect $T_{LAD}$. (A) The isolines shown on the phase diagram are for a constant methylation rate. The inset shows the three regimes of LAD thickness variation with chromatin-lamina affinity. (B) The isolines shown on the phase diagram are for a constant LAD thickness.

Regime II: Intermediate chromatin-lamina affinity: With increase in chromatin-lamina affinity, more chromatin is sequestered to the periphery, resulting in increased spreading of LADs along the lamina. Indeed, parametric increase in the chromatin-lamina affinity in the phase-field model shows the formation of elongated LADs (Fig 2C) shaped like a circular segment (Fig 2G) with radius comparable to the reaction-diffusion length scale $(R_{LAD} \sim l_{RD})$ and contact angles smaller than those in Regime I $(\theta < 90°)$. The growth of LADs is also accompanied by reduced spacing between neighboring LADs (Fig 2G). The dependence of LAD thickness on epigenetic rates and chromatin-lamina affinity in Regime II becomes non-linear (Eq S36) but can be evaluated numerically (Fig S8). As the chromatin-lamina affinity increases the LADs spread further, coming closer to their neighbors and reducing the LAD-free region along the lamina, thereby effectively increasing the average LAD thickness (as seen in Fig S8, Regime II). Moreover, we observe that increase in methylation rate increases the heterochromatin content at the periphery, thereby forming LADs of larger radius further amplifying the LAD thickness (Fig S8).

Regime III: High chromatin-lamina affinity: Within Regime II the elevation in chromatin-lamina affinity results in increased spreading of chromatin along the lamina bringing the neighboring LADs closer. Beyond a critical limit ($V_{LAD}^{c}$), we enter the Regime III where the LADs form a near continuous layer along the lamina (contact angle $\theta \sim 0$, Fig 2G, lower panel). The effective radius of such LAD approaches a large value ($R_{LAD} \sim \infty$, Fig 2G). Beyond the critical limit ($V_{LAD}^{c}$), any additional increase in the chromatin-lamina affinity alone does not alter the LAD thickness (Fig S8) since a continuous layer of LAD has already formed with maximal interaction between chromatin and lamina. On the other hand, as methylation rate increases (Fig S8) additional availability of heterochromatin increases the LAD thickness. The increased LAD thickness with increase in methylation facilitates an easier interaction between neighboring LADs, promoting the formation of a continuous layer of LADs. Hence, we observe (Fig S8) that with increasing methylation rates the transition into Regime III, occurs at a lower critical limit of chromatin-lamina affinity, i.e. $\gamma_{lh}^{c}$ reduces as methylation rate increases. In this regime, as contact angle $\theta \sim 0$ and $R_{LAD} \sim \infty$, consequently, $G\left(\frac{R_{LAD}}{l_{RD}}, \theta\right) \sim G(\infty, 0)$ is again independent of $R_{LAD}$. Thus, the dependence of thickness of LADs in Regime III on histone methylation rate $\Gamma_{me}$ scales as

$$T_{LAD} \sim \sqrt{\frac{D_h}{\Gamma_{ac}} \frac{\Gamma_{me}}{\Gamma_{ac} + \Gamma_{me}}} \qquad \text{(S38)}$$

and, as in Regime I, shows excellent agreement that observed numerically (Fig S9).

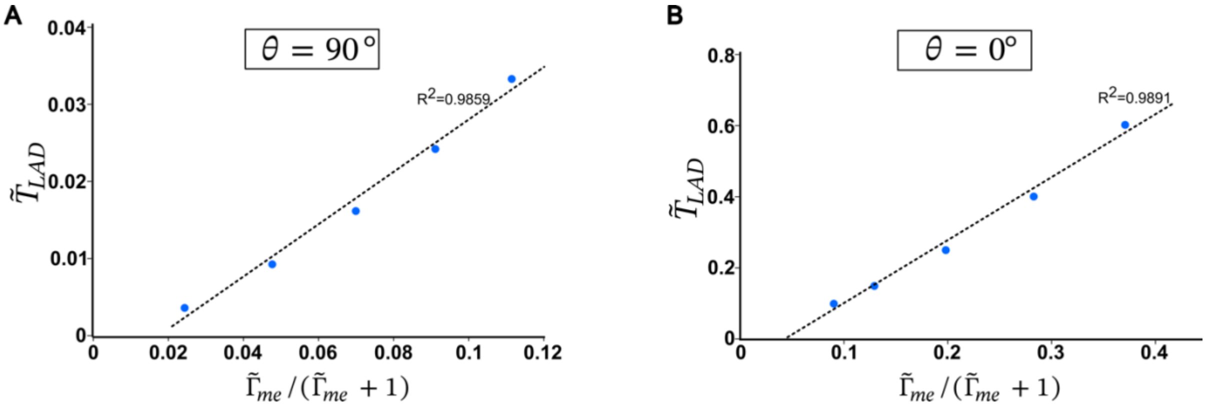


**Figure S9**: The scaling relation between $T_{LAD}$ and methylation rate obtained numerically for (A) small chromatin-lamina affinity and (B) high chromatin-lamina affinity.

For the specific morphologies of for the flat heterochromatin layer ($\theta \sim 0°$, $R_{LAD} \sim \infty$), the LAD thickness can also be derived from the balance between the reaction-diffusion driven influx and acetylation driven outflux to obtain the scaling relations equivalent to Eq S38 as follows:

Flat heterochromatin layer: Here we provide the derivation for thickness scaling that directly considers the layer of heterochromatin near the border. The reaction-diffusion equation Eq. S32 can be simplified to an 1D equation,

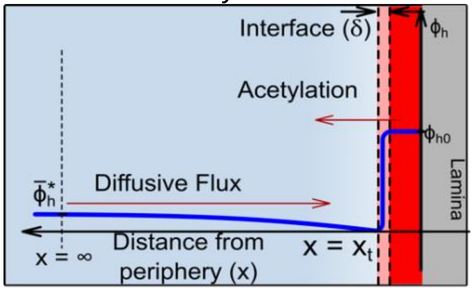


**Figure S10:** The competition of diffusion driven influx of heterochromatin with the epigenetic reaction driven outflux of heterochromatin from the LAD determines its steady state size.

$$D_h \frac{d^2\phi_h}{dx^2} - \Gamma_{ac}\phi_h + \Gamma_{me}\phi_e = 0 \tag{S39}$$

where $x$ denotes the distance from the interface between heterochromatin and euchromatin (Fig S10). Two different volume fractions $\phi_h^-$ and $\phi_h^+$ coexist immediately inside and outside the boundary of heterochromatin, respectively. Solving Eq S38 with boundary conditions $\phi_h|_{x_t^+} = \phi_h^+ = 0$ and $\phi_h|_{x=\infty} = \bar{\phi}_h^*$ gives,

$$\phi_h = -\bar{\phi}_h^* \, e^{-x/l_{RD}} + \bar{\phi}_h^* \tag{S40}$$

where $l_{RD}$ is the characteristic reaction-diffusion length scale given under a dilute limit as $l_{RD} = \sqrt{\frac{D_h}{\Gamma_{ac}}}$. Similar to Eq. S33, the change in the volume of the layer of heterochromatin can be written as,

$$\frac{dV}{dt} = S\frac{dT_{LAD}}{dt} = SD_h \frac{\partial\phi_h}{\partial x} - ST_{LAD}\Gamma_{ac} \tag{S41}$$

where S denotes the surface area of the heterochromatin layer. Solving Eq. S41 at the steady state by setting $\frac{dT_{LAD}}{dt} = 0$ yields

$$T_{LAD} = \frac{D_h\bar{\phi}_h^*}{\Gamma_{ac} l_{RD}} \sim \sqrt{\frac{D_h}{\Gamma_{ac}}}\,\bar{\phi}_h \tag{S42}$$

The relation in Eq. S42 agrees with Eq. S38.

On rescaling Eq S36 and S38

$$\tilde{T}_{LAD} \sim \frac{\tilde{\Gamma}_{me}}{\tilde{\Gamma}_{me}+1}\, G(\tilde{R}_{LAD}, \theta) \tag{S38}$$

$$\tilde{T}_{LAD} \sim \frac{\tilde{\Gamma}_{me}}{\tilde{\Gamma}_{me}+1} \tag{S39}$$

### S8 Model parameters, Initial and boundary conditions

To capture the spatiotemporal organization of chromatin in the nucleus, we numerically solve Eq S9 and S10. We consider an initial spatially homogenous distribution of chromatin in the nucleus with a no flux boundary condition of the order parameter $(\nabla\mu_d . \hat{\boldsymbol{n}}|_{\text{boundary}} = 0)$ ensuring the conservation of epigenetic marks. To mimic the intrinsic heterogeneities present in the nucleus, we add a random uniform noise to the initial chromatin configuration. In the model all length and time scales are normalized against the reaction-diffusion length scale $l_{RD}$ and $\Gamma_{ac}$ respectively. In rescaled model, the heterochromatin distribution is determined by rescaled methylation rate $\tilde{\Gamma}_{me}$ and rescaled chromatin-lamina affinity $\tilde{V}_{LAD}$. The initial values of these two parameters are chosen such that the simulation generate a distribution of HC domains comparable to the control nuclei (Fig S11). To represent the spatial heterogeneity of epigenetic reactions and chromatin-lamina interactions, Gaussian noise was applied to methylation reaction rates and chromatin-lamina affinity. All initial parameters are listed in Table S1.

**Table S1**: Values of the parameters used in simulation.

| | PARAMETER | DESCRIPTION | VALUE |
|---|---|---|---|
| **INITIAL CONDITIONS** | $\phi_h^{initial}$ | Initial heterochromatin content in the nucleus | 0.3 |
| | $\phi_e^{initial}$ | Initial euchromatin content in the nucleus | 0.7 |
| | $\phi_n^{initial}$ | Initial nucleoplasm content in the nucleus | $1 - \phi_h^{initial} - \phi_e^{initial}$ |
| | $\sigma_\phi^{noise}$ | Range of uniform perturbation due to heterogeneities in initial conditions | 0.01 |
| **EPIGENETIC KINETICS** | $\tilde{\Gamma}_{me}$ | Reaction rate of histone methylation (non-dimensionalised) | 0.38 |
| | $\sigma_\Gamma^{noise}$ | Variance in spatial distribution of reaction rates | 0.2 |
| **CHROMATIN-LAMINA INTERACTIONS** | $\tilde{V}_{LAD}$ | Chromatin-lamina interaction strength (non-dimensionalised) | 0.01 |
| | $\sigma_{\tilde{V}_{LAD}}^{noise}$ | Variance in spatial distribution of $\tilde{V}_{LAD}$ | 0.3 |

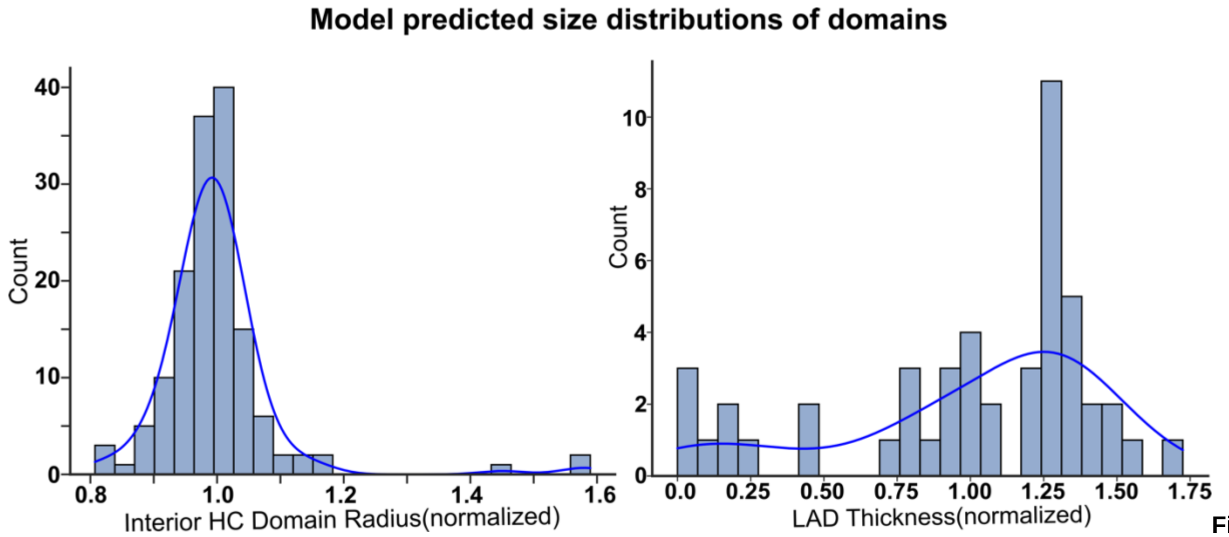


**Figure S11:** Size distribution of the heterochromatin domains in the interior of the control nucleus obtained numerically (left) and from STORM image(right).

To simulate the change in chromatin organization in chemo-mechanical altered environments, we vary the parameter $\tilde{\Gamma}_{me}$ and $\tilde{V}_{LAD}$ according to extracted parameter from the theoretical framework (listed in Table S2). The numerically observed change in heterochromatin domain sizes in interior and at the periphery of the nucleus (Fig S12) using the extracted biophysical parameters from STORM images agrees with the in-vitro observed chromatin reorganization.

**Table S2:** Values of parameters used to simulate nuclei undergoing pharmacological and environmental interventions.

| | $\tilde{\Gamma}_{me}$ | $\tilde{\gamma}_{lh}$ |
|---|---|---|
| **Control** | 0.038 | 0.01 |
| **GSK** | 0.024 | 0.0103 |
| **TSA** | 0.029 | 0.0088 |
| **Y27** | 0.043 | 0.0126 |
| **Stiff** | 0.032 | 0.0089 |
| **Glass** | 0.030 | 0.0085 |
| **Healthy Tenocytes** | 0.026 | 0.0089 |

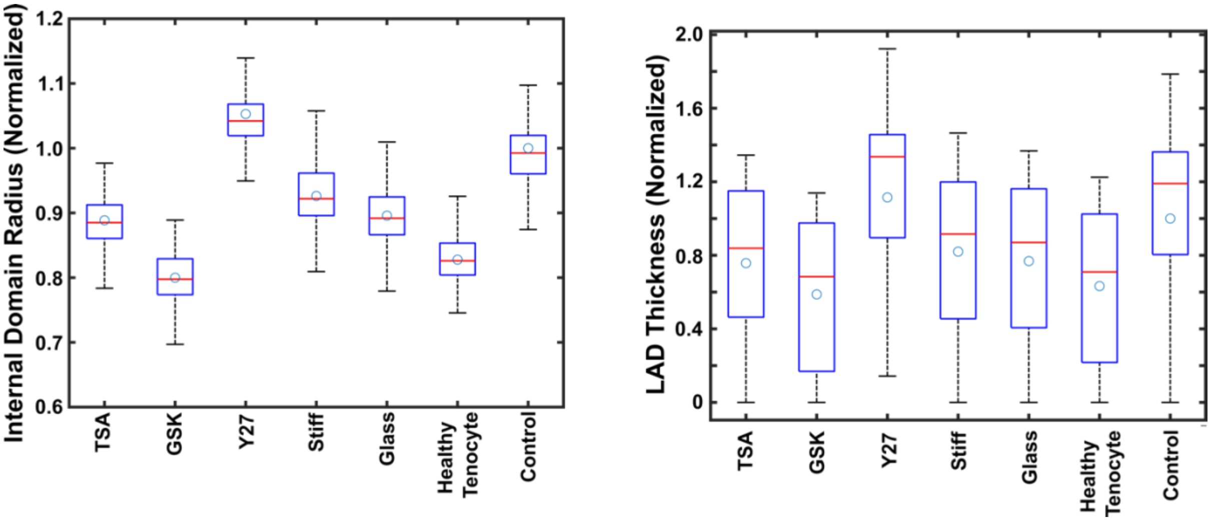


**Figure S12:** Numerical predicted change in heterochromatin domain size (A) and LAD thickness (B) in nuclei undergoing chemical-mechanical alterations.

### S9 Physiological values of model parameters

The numerical simulations are performed with rescaled parameters as discussed in Section S1.5. The rescaling of the mathematical model is done with respect to the intrinsic energy scale (chromatin-chromatin interaction strength), time-scale (reaction rates) and length-scale (reaction-diffusion length). To physically interpret our simulation results, the physiological values of these model parameters are evaluated as listed in Table S3.

**Table S3:** Value of non-dimensional parameters in physical dimensions

| | PARAMETER | DESCRIPTION | VALUE | REFERENCE |
|---|---|---|---|---|
| **ENERGY SCALE** | $c\sim\left(\frac{2-4k_BT}{V_N}\right)$ | Scale of chromatin-chromatin interactions | $\sim 8\times10^{-3} - 16\times10^{-3} pN/nm^2$ | [5] |
| | $l_{int}$ | Interface width of heterochromatin domains | $10-50\ nm$ Observed from ChromSTEM imageing | [4] |
| | $\kappa=(l_{int}^2\times c)$ | the penalty associated with the interface formation | $\sim 0.8pN$ | |
| | $\gamma=\left(\frac{1}{6}\sqrt{\kappa c}\right)$ | Surface tension | $\sim 10^{-2}pN/nm$ | |
| | $V_{LAD}^{max} = (\gamma cos\theta)$ | Maximum strength of chromatin-lamina interactions per unit area | $\sim 10^{-2}pN/nm$ | [6] |
| **TIME SCALE** | $\Gamma_{ac}$ | Reaction rate of histone acetylation | $\sim 10^{-2}\ s^{-1}$ | [7] |

| | | | | |
|---|---|---|---|---|
| | $\Gamma_{me}$ | Reaction rate of histone methylation | $\sim 10^{-3}\ s^{-1}$ | [8] |
| **LENGTH SCALE** | $D$ | Diffusivity of nucleosome | $\sim 10^{-3}\ \mu m^2/s$ | [9] |
| | $l_{RD} \sim \sqrt{\frac{D}{\Gamma_{ac}}}$ | Reaction-diffusion length scale | $\sim 300\ nm$ | |

### S10 Example of extraction process of biophysical parameters in control nuclei from STORM images using theory

Here, we present the extraction process for hMSCs nuclei cultured on glass under control conditions. We start by generating H2B density heatmaps from segmented super-resolution images using Voronoi tessellation (Fig S13 top panel). These images are processed using a custom MATLAB code (section2.4)[10] to obtain the statistical distributions of the interior domain radii $R_d$ and the LAD thickness $T_{LAD}$ for each nucleus (Fig S13 middle panel). The derived interior domain and LAD sizes are locally correlated with the methylation rate $\Gamma_{me}$ via Eqs S19 and S25, enabling the extraction of nucleus-wide distribution of $\Gamma_{me}$ (Fig S13 bottom panel). Finally, leveraging the extracted distributions of $\Gamma_{me}$ and LAD thickness, we apply Eq S36 (numerically depicted in Fig S8) to infer the distribution of chromatin-lamina affinity relative to chromatin-chromatin interaction strength $V_{LAD}$ along the nuclear periphery (Fig S13 bottom panel). Notably, the distributions of extracted parameters in individual nuclei feature characteristic means (red dot) that are comparable to the mean across all nuclei (blue dot).

In a similar fashion, we extract the distribution of biophysical parameters for all nuclei undergoing known pharmacological treatments and biomechanical stimuli. The quantitative changes in the mean values of these parameters, relative to the mean values of control nuclei for each treatment, are listed in Table S4.

The general trend observed from Table S4 indicates that a decrease (or increase) in the methylation rate $\Gamma_{me}$ is accompanied by a corresponding decrease (or increase) in the interior domain radii $R_d$. This aligns with our theoretical predictions in Eq S19 which evaluates the size of heterochromatin domains based on the kinetic balance between methylation-diffusion driven heterochromatin influx and acetylation driven euchromatin outflux.

However, as discussed in Section 3.1 (Eq 6) in the main manuscript, our theory also predicts that increased chromatin-lamina interactions can lead to substantial sequestering of chromatin towards the periphery, reducing the heterochromatin available in the nucleus interior. In such a scenario, even with increased methylation, some heterochromatin may move towards the periphery, resulting in reduction of interior heterochromatin domain sizes.

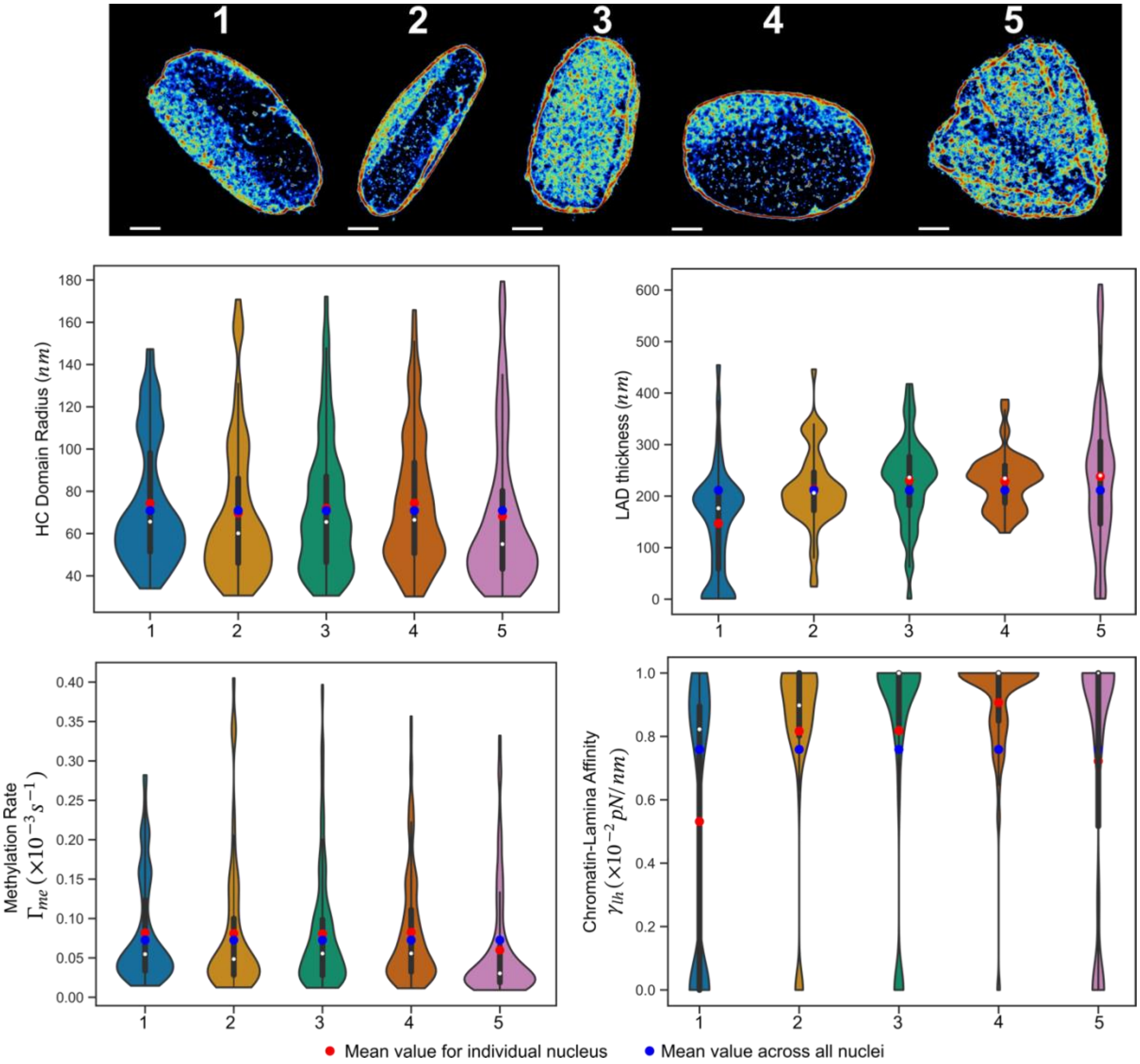


**Figure S13:** Top Panel: Voronoi density plots of H2B STORM images of hMSCs on glass nuclei cultured in control condition. Middle Panel: Quantification of STORM images showing distribution of domain radius and LAD thickness for individual nucleus. Bottom Panel: Distribution of histone methylation rate and chromatin-lamina affinity obtained by our integrative framework.

**Table S4**: Change in biophysical parameters relative to control nuclei

| | **QUNATIFIED FROM STORM IMAGES** | | **PREDICTED BY INTERGATIVE FRAMEWORK** | |
|---|---|---|---|---|
| | $R_d$ | $T_{LAD}$ | $\Gamma_{me}$ | $V_{LAD}$ |
| **GSK** | 18% (↓) | 20% (↓) | 36% (↓) | - |
| **TSA** | 11% (↓) | 26% (↓) | 22% (↓) | 11% (↓) |
| **Y27** | 4.3% (↓) | 19% (↑) | 12% (↑) | 26% (↑) |
| **STIFF** | 6.8% (↓) | 14% (↓) | 17 % (↓) | 10 % (↓) |

| | | | | |
|---|---|---|---|---|
| **GLASS** | 8% (↓) | 21.5% (↓) | 19.9% (↓) | 14.5 % (↓) |
| **TENDINOSIS** | 5.8% (↑) | 10.2% (↑) | 30% (↑) | 11.6% (↑) |

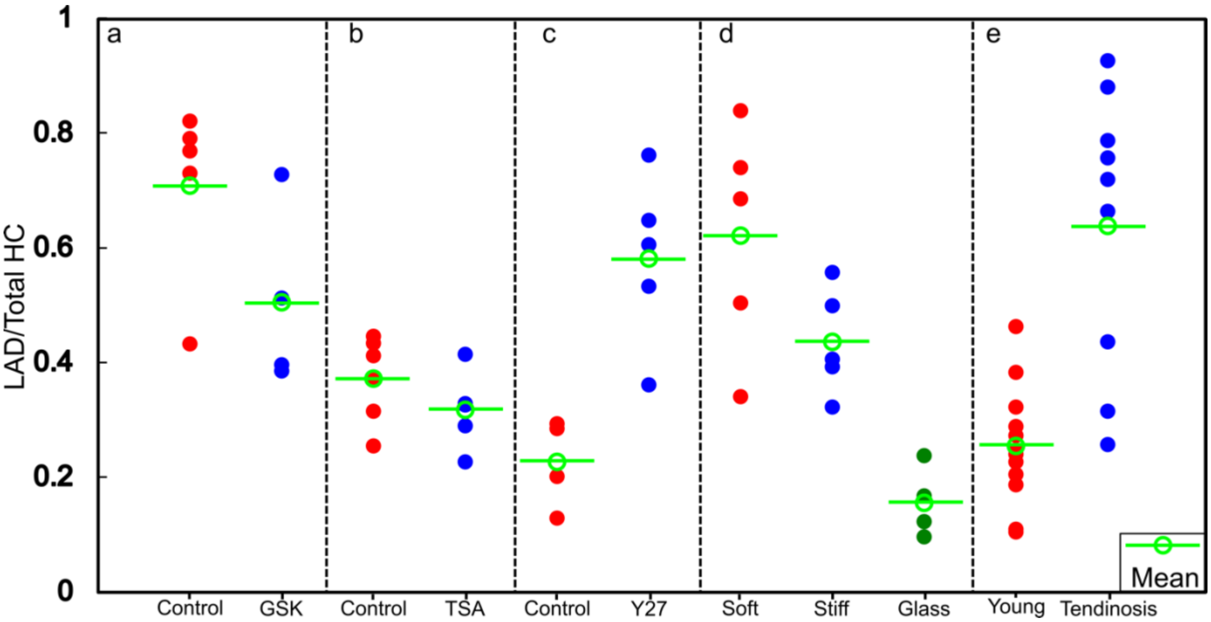

**Figure S14:** The quantification of heterochromatin localization to periphery relative to total heterochromatin in nucleus undergoing different treatments and control of each case (red plots).

This effect is observed in the case of Y27 treatment where methylation increases by 12% (Table S4), but chromatin-lamina affinity increases more significantly (by 26%), resulting in a slight (4.3%) reduction in the mean size of the interior domains. To confirm the significant sequestering of heterochromatin towards the periphery, we plot the ratio of peripheral heterochromatin to total heterochromatin for each nucleus after control and Y27 treatment (Fig S14c). Indeed, there was a significant increase in the relative amount of heterochromatin at the periphery after Y27 treatment.

Interestingly, similar sequestering of heterochromatin from nucleus interior to the periphery is also observed in some other treatments viz. on soft substrates (Fig S14d), after tendinosis (Fig S14e) and in the control case of GSK treatment (Fig S14a). However, Table S4 shows that in these cases the increase in chromatin-lamina affinity (which promotes chromatin sequestering to the lamina) was countered by the increase in methylation rate (which enlarges the interior domains). Thus, a balance between chromatin sequestering and histone methylation rate regulates interior heterochromatin domain sizes.

It is important to note that in all treatments, the effect of chromatin sequestering cannot be overlooked. For example, a 36% change in methylation rate (GSK treatment) results in 18% change in domain sizes when the chromatin lamina affinity is unchanged. However, after tendinosis a 30% change in methylation rate only resulted in 5.8% change in domain sizes. This is because tendinosis also resulted in 11.6% increase in chromatin-lamina affinity causing some heterochromatin to move towards the periphery thereby limiting the growth of interior domains.

### REFERENCES


1. Briand, N. and P. Collas, Lamina-associated domains: peripheral matters and internal affairs. Genome Biol, 2020. **21**(1): p. 85.

2. Kiseleva, A.A. and A. Poleshko, The secret life of chromatin tethers. FEBS Lett, 2023. **597**(22): p. 2782-2790.
3. Sanulli, S., et al., HP1 reshapes nucleosome core to promote phase separation of heterochromatin. Nature, 2019. **575**(7782): p. 390-394.
4. Kant, A., et al., Active transcription and epigenetic reactions synergistically regulate meso-scale genomic organization. Nature Communications, 2024. **15**(1): p. 4338.
5. Moller, J., J. Lequieu, and J.J. de Pablo, The free energy landscape of internucleosome interactions and its relation to chromatin fiber structure. ACS central science, 2019. **5**(2): p. 341-348.
6. Tolokh, I.S., et al., Strong interactions between highly dynamic lamina-associated domains and the nuclear envelope stabilize the 3D architecture of Drosophila interphase chromatin. Epigenetics Chromatin, 2023. **16**(1): p. 21.
7. Waterborg, J.H., Dynamics of histone acetylation in vivo. A function for acetylation turnover? Biochemistry and cell biology, 2002. **80**(3): p. 363-378.
8. Haws, S.A., et al., Intrinsic catalytic properties of histone H3 lysine-9 methyltransferases preserve monomethylation levels under low S-adenosylmethionine. Journal of Biological Chemistry, 2023. **299**(7).
9. Nozaki, T., et al., Condensed but liquid-like domain organization of active chromatin regions in living human cells. Science Advances, 2023. **9**(14): p. eadf1488.
10. Heo, S.-J., et al., Aberrant chromatin reorganization in cells from diseased fibrous connective tissue in response to altered chemomechanical cues. Nature biomedical engineering, 2023. **7**(2): p. 177-191.
